# A computational framework for mapping isoform landscape and regulatory mechanisms from spatial transcriptomics data

**DOI:** 10.1101/2025.05.02.651907

**Authors:** Jiayu Su, Yiming Qu, Megan Schertzer, Haochen Yang, Jiahao Jiang, Tenzin Lhakhang, Theodore M. Nelson, Stella Park, Qiliang Lai, Xi Fu, Seung-won Choi, David A. Knowles, Raul Rabadan

## Abstract

Transcript diversity including splicing and alternative 3’ end usage is crucial for cellular identity and adaptation, yet its spatial coordination remains poorly understood. Here, we present SPLISOSM (SpatiaL ISOform Statistical Modeling), a computational framework for detecting isoform-resolution patterns from spatial transcriptomics data. SPLISOSM leverages multivariate testing to account for spot- and isoform-level dependencies, demonstrating robust and theoretically grounded performance on sparse data. In the mouse brain, we identify over 1,000 spatially variable transcript diversity events, primarily in synaptic signaling pathways linked to neuropsychiatric disorders, and uncover both known and novel regulatory relationships with region-specific RNA binding proteins. We further show that these patterns are evolutionarily conserved between mouse and human prefrontal cortex. Analysis of human glioblastoma highlights pervasive transcript diversity in antigen presentation and adhesion genes associated with specific microenvironmental conditions. Together, we present a comprehensive spatial splicing analysis in the brain under normal and neoplastic conditions.

## Introduction

Almost all human genes undergo orchestrated pre-mRNA processing with precise spatiotemporal regulation^1,2^. The brain exhibits particularly high transcript diversity^3^, where region-specific expression of RNA binding proteins (RBPs) drives specialized local transcript landscapes that control neurogenesis^4,5^ and synaptic plasticity^6,7^. Disruption of this delicate regulatory balance contributes to numerous neurological disorders^8–10^ and cancer progression^11–13^. Despite its critical importance, our understanding of how RNA isoforms are spatially organized within tissues remains limited.

Modern spatial transcriptomics (ST) platforms naturally provide some isoform-specific information, whether through 3’ fragments in short-read approaches^14–16^, full-length transcripts in long-read sequencing^17,18^, or exon probes in imaging-based platforms^19,20^ (Figure 1A). Despite this, most ST analysis pipelines aggregate expression at the gene level, discarding rich transcript information. Few computational approaches^21–23^ detect spatial patterns in isoform distribution, creating a significant gap in our understanding of the molecular complexity and its regulation in shaping tissue architecture.

**Figure 1:**
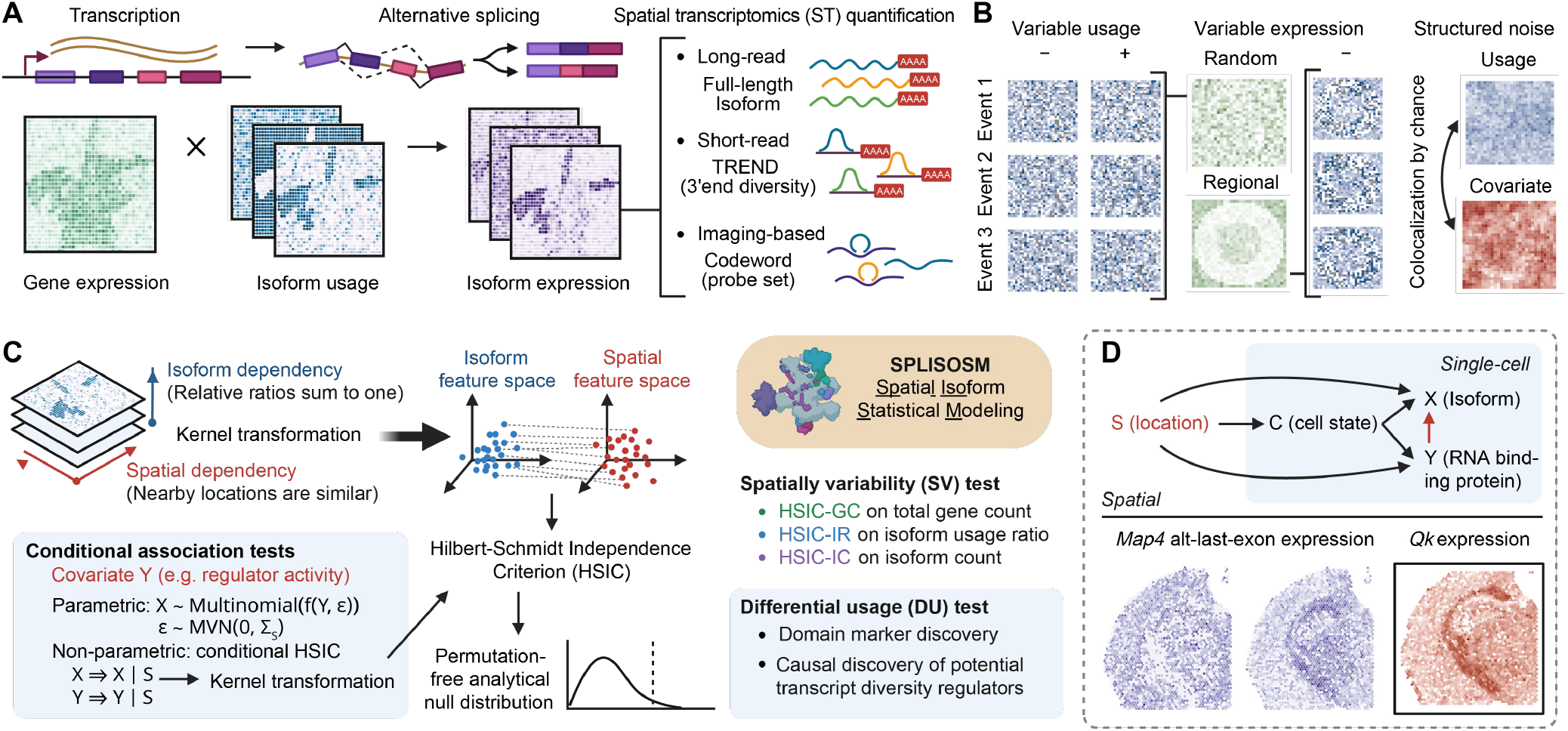
A computational toolbox for spatial isoform pattern discovery. **(A)** Alternative pre-mRNA processing generates multiple isoforms from the same gene, creating varying molecular transcript diversity detectable by major spatial transcriptomics (ST) platforms. **(B)** Isoform quantification from ST data is multivariate, sparse, and confounded by gene expression and spatial autocorrelation. **(C)** SPLISOSM addresses challenges in spatial isoform analysis by recasting spatial variability (SV) and differential isoform usage (DU) detections as multivariate independence tests. For SV testing, isoform usage and spatial coordinates are projected into feature spaces to assess nonlinear dependencies using the Hilbert-Schmidt Independence Criterion (HSIC) with optimized spatial and isoform kernels. For DU testing, nonparametric conditional HSIC test and parametric generalized linear mixed models (GLMM) control for confounding spatial autocorrelation and reduce spurious associations. **(D)** SPLISOSM enables cell-type-agnostic detection of spatially variably processed (SVP) genes and their potential regulators.

Analyzing spatial isoform patterns presents several formidable challenges (Figure 1B). First, isoforms from the same gene are interdependent, with expression constrained by total transcribed pre-mRNAs. This multivariate and compositional nature invalidates standard statistical approaches designed for gene expression analysis. Early attempts either inappropriately treated isoforms as independent^18^ or relied on oversimplified univariate metrics^21,22^, significantly compromising statistical power (Supplementary Note). Second, data sparsity – already problematic in ST – is exacerbated at the isoform level when gene counts are subdivided across multiple transcripts. Regional expression variation can lead to zero total UMI counts at many locations, making isoform ratios undefined. Even when detected, these ratios frequently appear binary despite representing continuous biological preferences^24,25^, complicating operations like log-ratio transformations. Common workarounds such as pseudo-counts introduce additional biases that distort relationships between observations^26^. Finally, spatial autocorrelation violates the independent observation assumptions fundamental to most differential association tests, generating false positives from spurious associations that merely reflect mutual dependence on spatial coordinates.

To address these challenges, we present SPLISOSM (SPatiaL ISOform Statistical Modeling), a computational framework for analyzing isoform usage in ST data. Across multiple datasets of adult mouse and human brain, we identify thousands of genes underwent spatially variable processing. To uncover regulatory mechanisms, we test for conditional association between isoform ratios and potential regulators like RBPs, confirming known connections and discovering novel region-specific regulation. Finally, we extend our analysis to glioblastoma cohorts, revealing how transcript diversity is shaped by both normal brain architecture and disease-specific alterations in the tumor microenvironment.

## Results

### Designing isoform-level spatial variability and association tests

Spatial gene variability arises from both cell type distribution patterns and location-specific variation within cell types. While several methods exist for identifying spatially variably expressed genes^27–31^, they prioritize different spatial patterns, resulting in limited consensus^32^. Moreover, these approaches cannot be easily adapted for multivariate isoform analysis due to their reliance on specific model assumptions (Supplementary Note). Drawing on non-parametric statistics’ effectiveness in modeling total expression from sparse ST data^30^, we reformulated spatial variability detection and differential usage analysis as multivariate independence testing. Specifically, SPLISOSM transforms isoform compositions, spatial coordinates and other covariates of interest into feature spaces, capturing their complex relationships through the kernel-based Hilbert-Schmidt Independence Criterion (HSIC)^33,34^ (Figure 1C). Significant patterns are identified by comparing against the null hypothesis that a gene’s isoform preference is independent of either spatial locations or a given covariate such as RBP expression (Methods).

Our approach introduces two key technical innovations: First, we mathematically prove that all low-rank approximations of kernel tests – including those in SPARK-X^30^ – inevitably sacrifice statistical power (Theorem 1, Supplementary Note). This insight led us to develop enhanced spatial kernels^35^ using spectral graph theory that prioritize signals by their spatial frequency, limiting power losses to high-frequency fluctuations representing noise when processing large-scale data. Second, we established a rigorous framework for sparse isoform data with a new compositional kernel that handles undefined ratios (Theorem 2, Supplementary Note), optimizing efficiency while maintaining test validity through a mean replacement strategy that utilizes all spatial observations.

Based on these theoretical advances, SPLISOSM implements three complementary spatial variability tests: HSIC-GC for total gene expression, HSIC-IR for isoform usage ratios, and HSIC-IC for isoform counts. For genes showing spatial variability in isoform preference, we apply conditional differential usage tests to identify potential RBP regulators (Methods). Using spatial variability, SPLISOSM enables cell-type-agnostic discovery (Figure 1D), complementing existing single-cell isoform studies^36–38^ while offering unique advantages in contexts where cell typing is challenging, such as in tumor microenvironments.

### SPLISOSM produces well-calibrated and permutation-free p-values in simulation

We first validated SPLISOSM through carefully designed simulations reflecting both gene expression and isoform preference variation (Figure 2A, Methods). Our spatial variability tests target distinct biological signals with well-calibrated statistics (Figure 2B). Among them, HSIC-IR correctly identified spatial patterns in isoform usage while ignoring changes in total gene expression (Scenarios 4 vs 5). The modified linear kernel with mean replacement provided optimal power without introducing artifacts (Figure S1A-B). Complementarily, HSIC-IC detected both isoform usage (Scenarios 1 vs 3) and gene expression spatial variation (Scenarios 1 vs 4). We also observed improvements in the gene-level HSIC-GC compared to SPARK-X through the optimized spatial kernel (Scenarios 4-6), validating our theoretical predictions.

**Figure 2:**
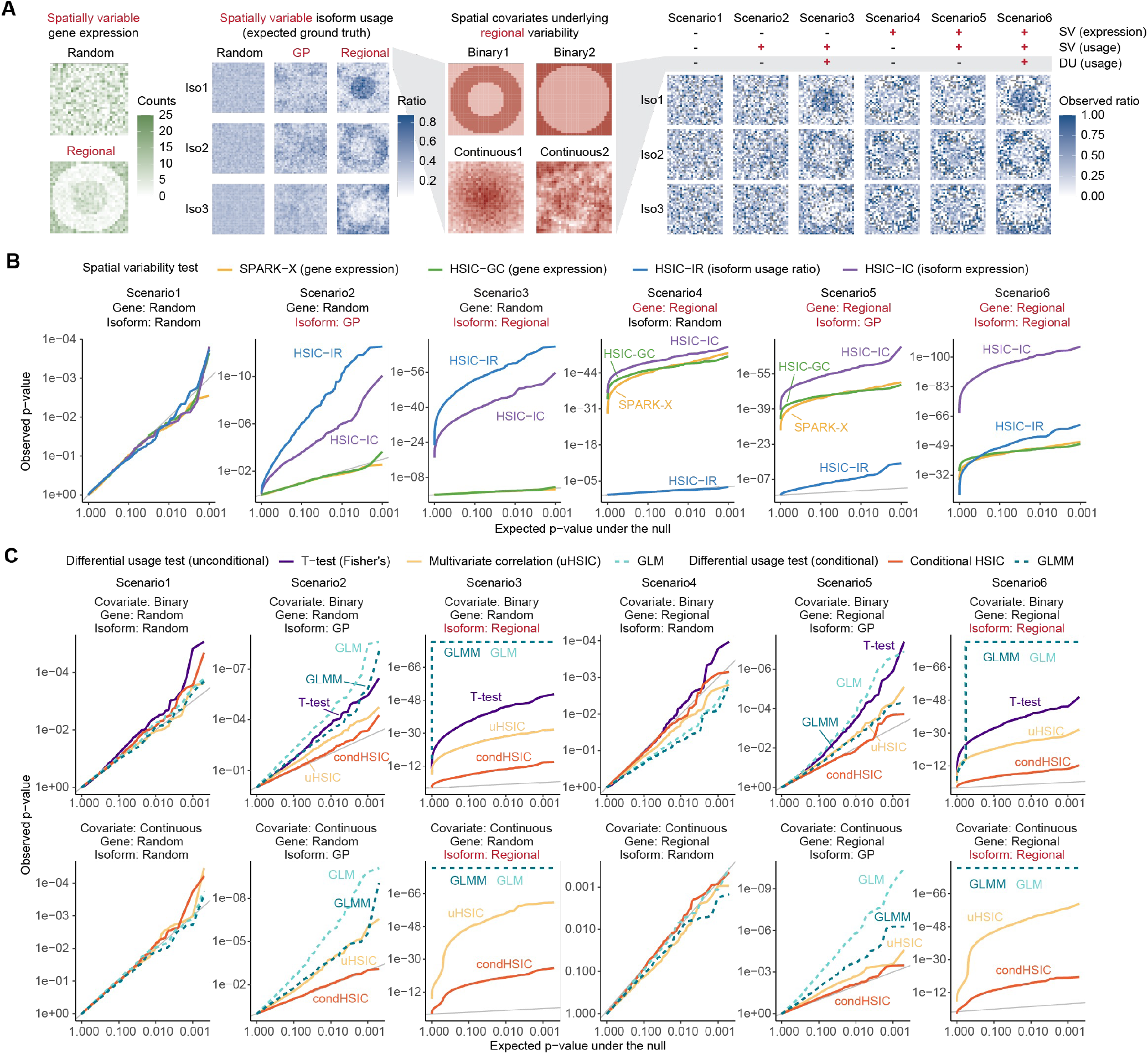
SPLISOSM produces well-calibrated and permutation-free p-values in simulation. **(A)** Six simulation scenarios with spatial variability at different layers. Regional variability was generated from two binary and two continuous spatial covariates. **(B)** Q-Q plots of SV test p-values across 1,000 simulated genes per scenario. Regional gene expression (Scenarios 4-6) was included to introduce artifacts in the observed ratios. **(C)** Q-Q plots of DU test p-values aggregated from 2,000 tests (1,000 genes × 2 covariates) per covariate type. Variable isoform usage independent of covariates (Scenarios 2 and 5) was sampled from a Gaussian process (GP) to mimic spurious differential associations. For binary covariates, T-tests were performed on individual isoforms with p-values combined at gene-level using Fisher’s method. For GLM and GLMM, null models with zero effect size were fitted and tested using the score statistic.

For association analysis, we focus on the challenge of correlated noise, which creates patterns that falsely align with covariates (Scenarios 1 vs 2). Conventional approaches, including discrete T-tests, continuous multivariate correlation, and generalized linear models (GLMs), all exhibited significant p-value inflation under spatial confounding (Figure 2C, Scenarios 1 vs 2 and 4 vs 5). SPLISOSM overcomes this limitation by introducing spatial conditioning within existing differential analysis frameworks (Methods). Simulations confirmed that our non-parametric conditional HSIC maintains proper null calibration while preserving sensitivity to subtle covariate effects (Figure 2C, Figure S1C-E), and that our parametric generalized linear mixed model (GLMM) partially controlled false positives compared to standard GLMs.

### Integrative analysis reveals spatial alternative splicing programs in adult mouse brain enriched for synaptic and membrane trafficking functions

Recent studies^17,18^ have generated spatial isoform maps of postnatal and adult mouse brains by combining 10X Visium spatial barcoding with Oxford Nanopore (ONT) sequencing. While able to resolve full-length transcripts, these long-read ST technologies are limited by throughput and data sparsity, providing a natural testbed to demonstrate SPLISOSM’s utility in biological discovery. We reanalyzed two SiT^18^ coronal brain section samples (CBS1, CBS2) from adult mouse (Figure 3A). SPLISOSM detected 150 spatially variably processed (SVP) genes shared by both replicates (adjusted p-value < 0.05, Figure 3B and Figures S2-3) – more than double the originally reported 61 genes – without relying on region annotations. Our spatial variability tests demonstrated strong consistency between replicates (Spearman’s ρ of p-values = 0.52, 0.39, 0.43 for HSIC-IR, HSIC-GC, and HSIC-IC, respectively), while SPARK-X biased toward the sample with higher sequencing coverage (Spearman’s ρ = 0.26). These SVP genes were enriched in membrane trafficking and synaptic signaling (*Clta, Cltb, Snap25, Stxbp1, Ap2a1, Atp6v1e1, Napa*), with many implicated in neurodegenerative diseases (Figure 3C). We also observed widespread isoform variability in ribosomal genes (*Rps24, Rps6, Rpl13a, Rpl5, Rps9*), aligning with previous studies showing cell-type and tissue-specific ribosome composition^39^. Importantly, transcript usage variability alone proved sufficient to define brain regions with comparable spatial resolution to expression-based clusters (Figure 3D, Methods), and isoform preference could be grouped into region-specific programs by gene-wise clustering (Figure 3E, Methods).

**Figure 3:**
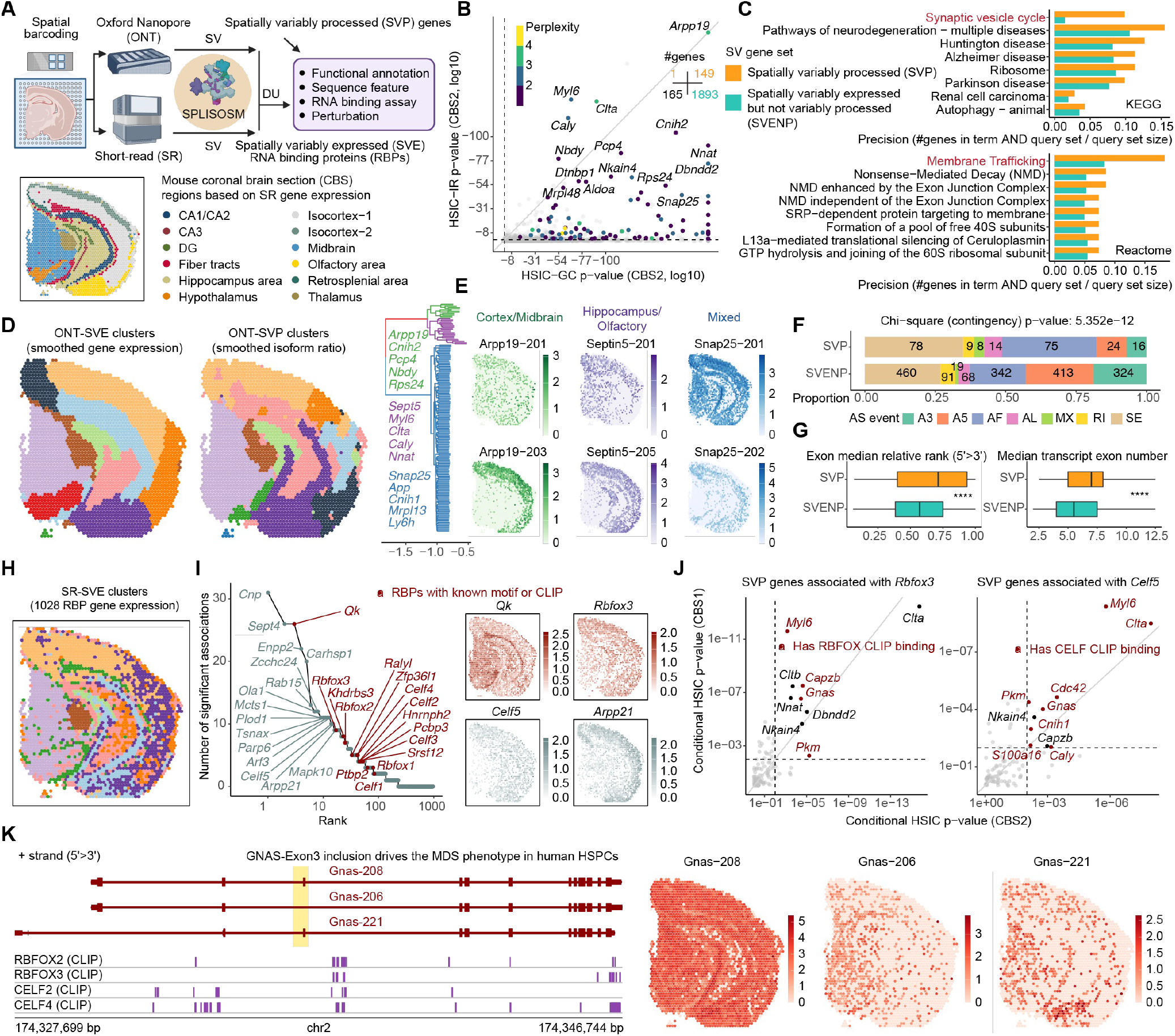
Integrative analysis reveals spatial alternative splicing programs in adult mouse brain enriched for synaptic and membrane trafficking functions. **(A)** Workflow overview. **(B)** Comparison of spatial variability in gene expression (x-axis) versus isoform usage (y-axis) in the CBS2 sample. Points colored by expression perplexity (effective number of isoforms per gene). Inset shows number of events significant in both CBS1 and CBS2 replicates (adjusted p-value < 0.05), categorized as spatially variably processed (SVP) or spatially variably expressed but not variably processed (SVENP) genes. **(C)** Pathway enrichment analysis comparing SVP versus SVENP genes, with terms arranged by precision differences between the two groups. **(D)** Unsupervised clustering based on gene-level expression of SVE genes (left) or relative isoform ratios of SVP genes (right). Isoform counts were locally smoothed to compensate for data sparsity. **(E)** Selected isoform usage programs identified through hierarchical clustering of SVP genes. **(F)** Distributions of alternative splicing event types among SVP and SVE genes. **(G)** Selected exon features of skipping exons and mutually exclusive exons in SVP and SVENP genes. Boxplots show median relative rank (center line), interquartile range (box), and 1.5× interquartile range (whiskers); ****: T-test p < 0.0001. **(H)** Unsupervised clustering based on 1,028 SVE RBP expression from short-read sequencing data. **(I)** Top RBPs ranked by number of significantly associated targets, color-coded by availability of documented motifs (CISBP-RNA) or CLIP data (POSTAR3). Log-normalized expression of selected RBPs shown at right. **(J)** Differential isoform usage p-values for Rbfox3 (left) and Celf5 (right) in two replicates. Red points indicate validated targets with RBP binding near regulated events in brain-CLIP data. **(K)** *Gnas* transcript structure and spatial log-normalized expression patterns.

At the structural level, SVP genes showed enrichment for exon skipping particularly near the 3’ end, as well as alternative first exons (Figure 3F-G). To identify potential modulators behind these events, we systematically searched for differential association with spatially variably expressed RBPs, which collectively reproduced brain major compartments (Figure 3H and Figure S4) and may act through direct binding or as indirect cofactors to drive transcript regional specificity. Top RBPs prioritized in our analysis included well-established neural splicing regulators like QKI, RBFOX, and CELF families and less-studied ones like ARPP21 (Figure 3I). To evaluate these relationships, we examined available brain CLIP datasets^40^, confirming 4 of 9 significant *Rbfox3* associations. Although CELF5-specific CLIP data remains unavailable, 8 of 10 of its associated target genes showed binding by other CELF family members, supporting CELF5’s role as a major neural splicing modulator in adult mouse brain (Figure 3J).

Among SVP genes, *Gnas* showed particularly intriguing associations (Figure 3K). Its exon 3, which is spliced out specifically in midbrain and fiber tracts, generates a hyperactive G-alpha protein driving differentiation defects and myeloid malignancies when included in SRSF2 and U2AF1 mutants^41^. Our analysis reveals that RBFOX and CELF families potentially regulate *Gna*s exon 3 skipping in the brain, raising the possibility that neural RBP dysregulation in non-neural tissues could contribute to disease pathogenesis.

### Cross-platform validation of spatial transcript variability in adult mouse brain

The substantial spatial transcript processing identified from long-read data prompted us to examine whether these patterns could be detected across different ST platforms, each recognizing distinct aspects of molecular variation. In addition to alternative splicing, mammalian nervous systems exhibit active alternative polyadenylation (APA)^42^ that generates transcript diversity near the 3’ end (TREND^43^), which can be captured by 3’-based ST sequencing protocols. In a separate 10X Visium sample^44^ of adult mouse brain, we identified 815 SVP genes with spatially variable TREND usage (adjusted p-value < 0.01, Figure 4A-B, Methods). The increased detection power reflected the higher sequencing depth of short-read Visium, as confirmed through down-sampling (Figure S5A). Analysis of a near-single-cell Slide-seqV2 dataset of mouse hippocampus^16^ revealed similar albeit fewer TREND patterns due to its sparsity (Extended Data Figure 1A). Remarkably, 95 SVP genes overlapped between long-read and short-read datasets despite their technical differences (Extended Data Figure 2A-B). Furthermore, genes using variable isoforms across space also show significantly higher spatial variability in their 3’ end regions detectable by short-read sequencing (Extended Data Figure 2C, Figure S6), suggesting coordinated transcript processing across the full transcript length^45^.

**Figure 4:**
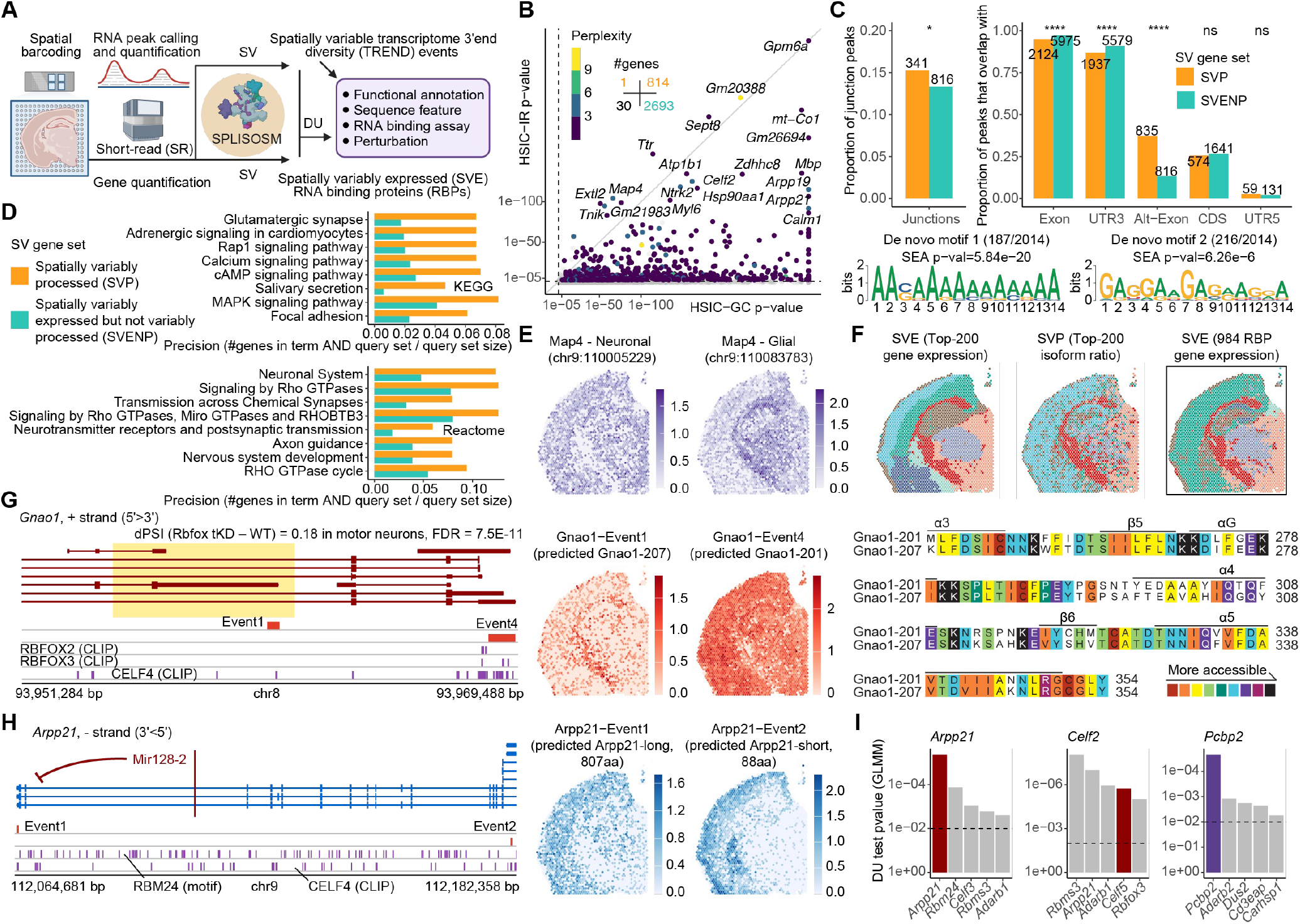
Spatial 3’end transcript diversity in adult mouse brain extends beyond alternative polyadenylation and shows functional convergence on signaling pathways. **(A)** Workflow overview. **(B)** Comparison of spatial variability in gene expression (x-axis) versus TREND usage (y-axis) in the short-read coronal brain section sample. Points colored by expression perplexity (effective number of isoforms per gene). Inset quantifies significant events (adjusted p-value < 0.01). **(C)** TREND event annotation with overlapping with genomic features. Alternative exons (Alt-Exon) are defined as exons located within introns of other transcript variants. De novo motif enrichment results shown at bottom. **(D)** Pathway enrichment analysis comparing SVP and SVENP genes, arranged by precision. **(E)** Spatial log-normalized expression of the neuronal (chr9:110005229) and glial (chr9:110083783) *Map4* variants. **(F)** Unsupervised clustering based on gene-level expression of top 200 SVE genes (left), relative TREND ratios of top 200 SVP genes (middle), and gene expression of all SVE RBPs (right). **(G)** *Gnao1* transcript structure and spatial log-normalized expression patterns. The likely protein sequence variation, color-coded by solvent-accessible surface area, is shown on the right. **(H)** *Arpp21* transcript structure and spatial log-normalized expression patterns. **(I)** Top RBPs associated with TREND usage in *Arpp21, Celf2* and *Pcbp2*. Dashed line indicates p = 0.01.

To obtain orthogonal validation, we reanalyzed a 10X Xenium Prime 5K dataset^46^ of adult mouse brain with subcellular spatial resolution through multiplexed in situ hybridization (Methods). Although most predesigned Xenium probes do not target sequencing-based variable regions, we observed strong gene-level agreement: 55.6% (20/36) of ONT-detected and 39.8% (100/251) of short-read-detected SVP genes showed significant spatial exon usage variability, both far exceeding the baseline rate of 14.9% (Extended Data Figure 2D-G). At single-cell resolution, we could pinpoint cellular origins of spatial transcript patterns. For example, Xenium confirmed that *Dtnbp1* isoform switching detected by long-read in specific brain regions was driven by distinct neuronal subpopulations: *Nptxr*+/*Slc17a7*+ excitatory neurons in in olfactory and hippocampus-CA3 and GABAergic interneurons in isocortex (Extended Data Figure 3). Similarly, we validated the *Zdhhc8* 3’ end variation and its isocortex and hippocampus-DG specificity (Figure S7).

This cross-platform convergence demonstrates that spatial transcript diversity represents a fundamental organizing principle of brain architecture, detectable across diverse technical approaches (Extended Data Figure 4, Figure S8). It also establishes SPLISOSM as a robust and cell-type-agnostic framework for uncovering transcript patterns invisible at gene level from ST data.

### Spatial 3’ end transcript diversity in adult mouse brain extends beyond alternative polyadenylation and shows functional convergence on signaling pathways

Although over 80% of spatially variable TREND events mapped to 3’ UTRs, we found significant localization at junctions and alternative exons (Figure 4C). Some non-UTR events represented internal poly(A) priming sites, supported by A-rich motif enrichment, allowing us to detect upstream diversity deeper into transcripts. The functional importance of TREND became evident as 25% of events overlapped coding regions. Of 7 previously reported protein-altering poly(A) site shifts between neurons and glia^47^, we recapitulated 6 (*Map4, Atp2a2, Cdc42, Klc1, Itsn1, Ptpn2*) while providing new insights into their spatial distribution (Figure 4E and Extended Data Figure 1B). SVP genes converged on key neural signaling pathways, including glutamatergic and adrenergic synaptic transmission and Rap1, Ca^2+^, cAMP, and MAPK cascades (Figure 4D). We observed additional protein-coding changes in genes involved in diverse cellular functions (Extended Data Figure 1C), including proteostasis (*Hsp90aa1, Hsp90ab1*), membrane trafficking and secretion (*Septin8, Septin11, Ap1s2, Olfm1, Lgi3*), and enzymatic activities (*Kalrn, Rexo2, Ppp2r5c*). Finally, major brain regions could be defined using TREND preference alone (Figure 4F, Figure S9), in agreement with our long-read analysis.

To understand the spatial regulation of TREND, we performed association analysis between event usage and RBP expression. Contrary to conventional thinking, spatial variability in core alternative polyadenylation machinery components was insufficient to explain the observed 3’ end diversity patterns (Extended Data Figure 1D). Instead, neural splicing regulators from our ONT analysis such as QKI, RBFOX, and CELF families also emerged as top potential regulators for APA, supporting their central role in shaping the adult mouse brain transcriptome^48^. A prominent RBFOX target is *Gnao1*, encoding a subunit of the heterotrimeric G proteins. Our analysis uncovered previously unrecognized spatial divergence in *Gnao1* last exon usage between cerebral nuclei and fiber tracts versus other brain regions, generating two protein isoforms with different C-termini (Figure 4G). Using public CLIP^40^ and knockout data^49^, we confirmed that RBFOX binds and promotes *Gnao1* downstream exon usage. Regional isoform specificity in *Gnas* and *Gnao1* extends to other G proteins (*Gnai2, Gnas, Gnal, Gnaq*) and G-protein-coupled receptors (*Gabbr1, Grm5, Adgrl1*), indicating a role of alternative transcript processing in fine-tuning signal transduction for spatially diverse physiological functions.

Our catalog also brought to light self-regulatory mechanisms driving transcript diversity in RBP genes themselves. For instance, the last exon usage of *Arpp21* correlated with its total expression (Figure 4H), consistent with a regulatory model where miR-128-2, produced from the longer isoform’s second-to-last intron, inhibits *Arpp21* expression^50^. We also verified self-regulation in the 3’ UTR of CELF family^51^ (Figure 4I) and discovered additional spatially variable 3’ UTR lengths across RBPs (*Elavl2, Elavl3, Qk, Rbfox1, Hnrnpa2b1, Hnrnpk*). One particularly intriguing new finding involves *Pcbp2*, which showed enriched usage of a minor upstream last exon in isocortex and thalamus. This alternative exon encodes a protein variant with an extended disordered C-terminus (Extended Data Figure 1E), whose usage inversely correlated with overall *Pcbp2* expression in both mouse brain and across human tissues (Extended Data Figure 1F). CLIP data^52^ revealed stronger human PCBP2 binding to this minor last exon compared to the canonical one, suggesting selective auto-regulatory feedback contributing to spatial specialization.

**Extended Data Figure 1:**
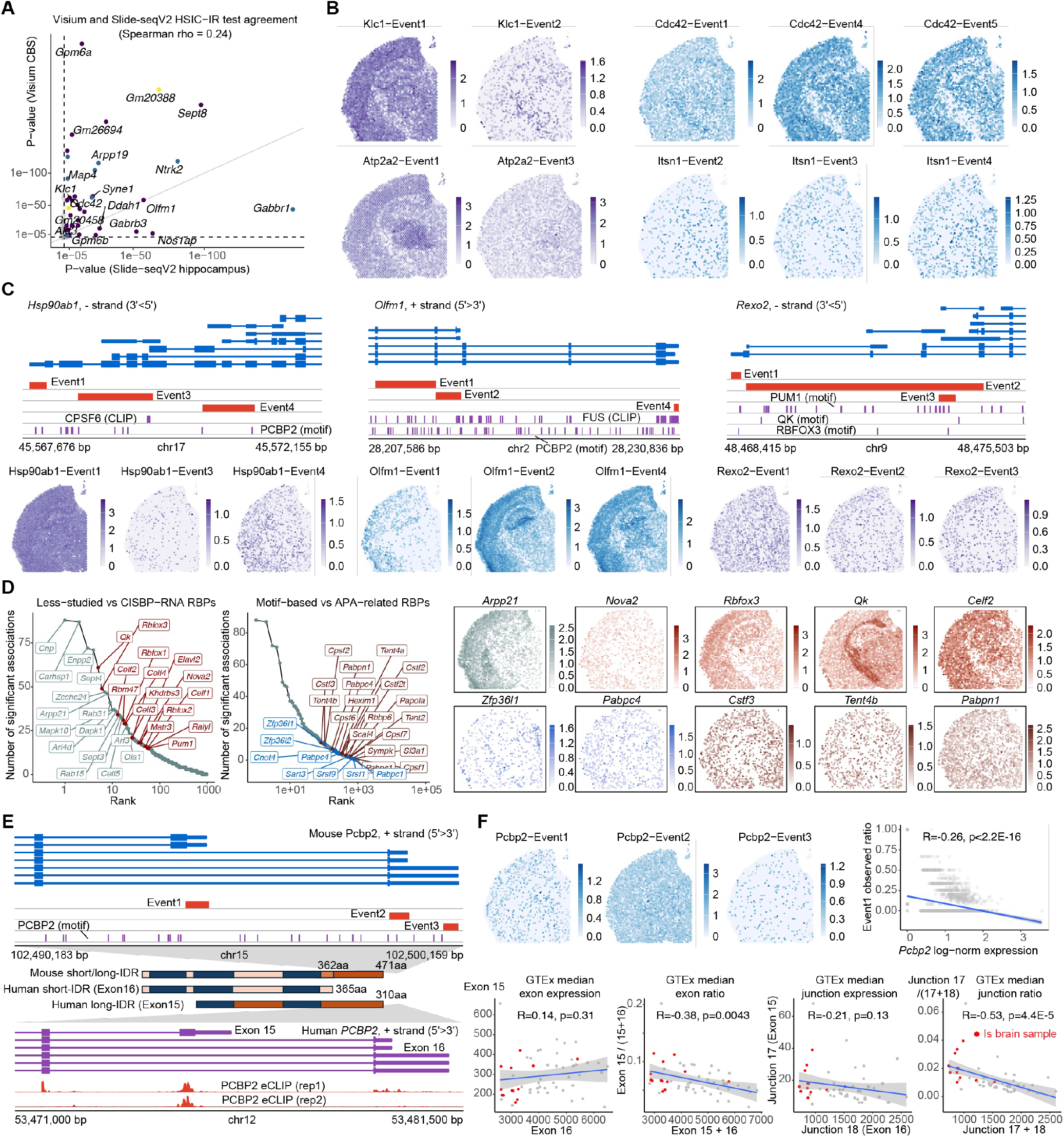
Spatially variable TREND events and their regulation in adult mouse brain, related to Figure 4. **(A)** HSIC-IR test agreements between Visium CBS (y-axis) and Slide-seqV2 hippocampus (x-axis) datasets. Points colored by expression perplexity. **(B)** Differential protein-altering alternative polyadenylation events between neurons and glial cells. Related to Figure 4E. **(C)** Additional protein-altering TREND events and their spatial log-normalized expression. **(D)** Top RBPs ranked by number of significantly associated targets, color-coded by available motifs (CISBP-RNA) or CLIP data (POSTAR3), core APA machinery involvement and motif enrichment near SVP TREND events. Spatial log-normalized expression of selected RBPs shown at right. **(E)** PCBP2 transcript structure in mouse and human, along with eCLIP data from ENCODE. **(F)** PCBP2 isoform expression in spatial mouse CBS data (top) and across GTEx human tissues (bottom).

**Extended Data Figure 2:**
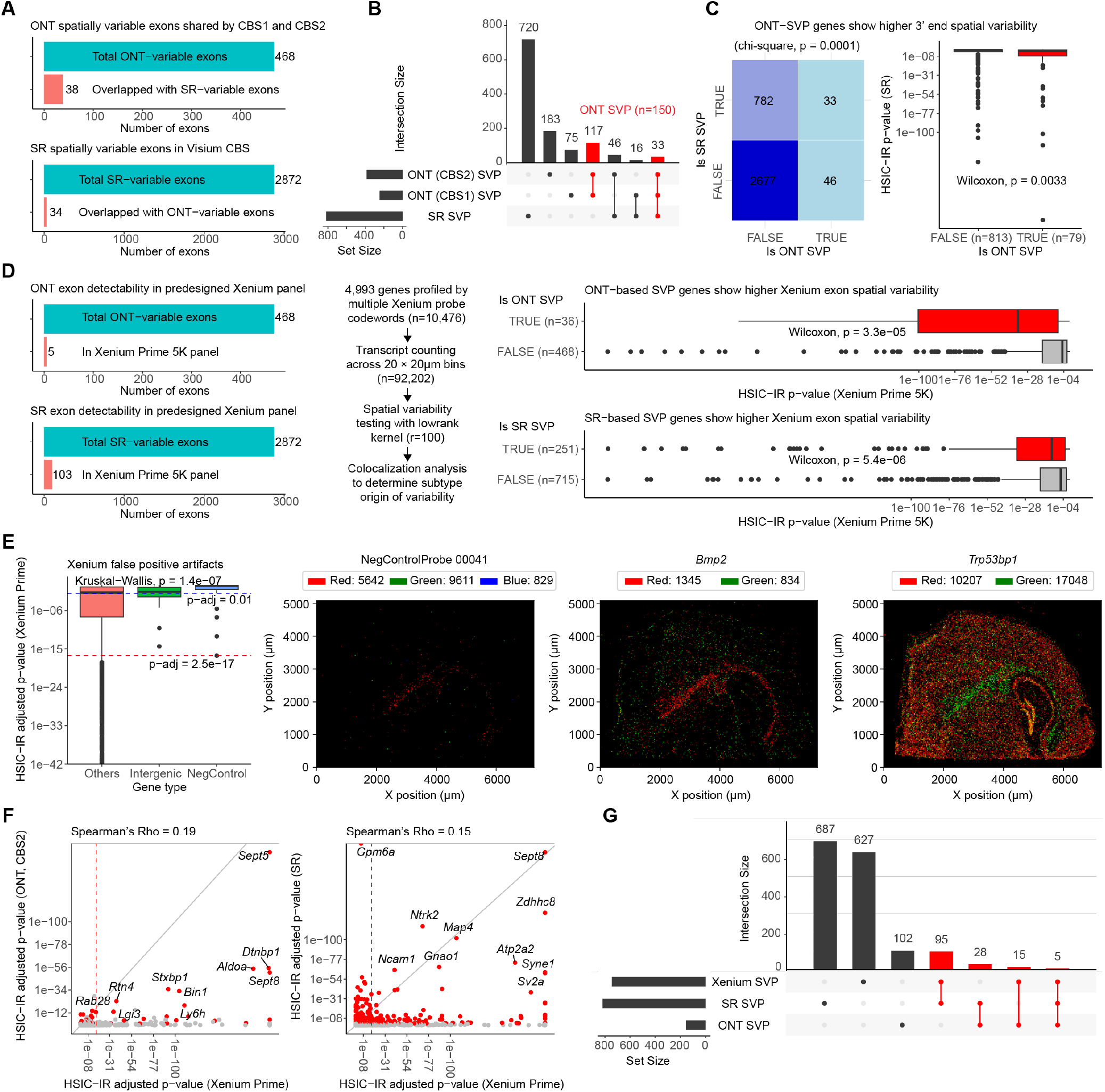
Comparison of spatial transcript diversity detection in adult mouse brain across ST platforms. **(A)** Limited exon overlap between ONT and short-read (SR) detected variable events. Isoforms from the 150 ONT-SVP genes in Figure 3 and TREND events from the 815 SR-SVP genes in Figure 4 were decomposed into 214 and 2,872 spatially variable exons, respectively. **(B)** SVP gene overlap between ONT and SR mouse CBS samples. Events were identified using adjusted p-value thresholds: ONT-SVP (p-adj ≤ 0.05), SR-SVP (p-adj ≤ 0.01). 150 genes shared by both ONT replicates are considered as ONT-SVP genes in following panels. **(C)** Gene-level SVP test agreements between ONT and SR. Left: Chi-square test applied to compare SR- and ONT-based SVP calling results. Right: SR HSIC-IR p-value distributions where boxplots show median (center line), interquartile range (box), and 1.5× interquartile range (whiskers). Wilcoxon rank-sum test was used to compare p-values across groups. Sample sizes and SVP annotation F (false) and T (true) indicated. Negative background numbers vary as not all genes detected/tested in both platforms. **(D)** Left: Limited exon overlaps across platforms. Right: HSIC-IR p-value distributions from the Xenium 5K adult mouse brain sample. **(E)** Assessment of technical artifacts in Xenium spatial patterns. Left: HSIC-IR adjusted p-values for non-gene features, showing that intergenic regions and negative control probes can generate false-positive spatial patterns (p-adj ≤ 0.01 threshold indicated by the blue dashed line). Kruskal-Wallis test was used to compare p-value distributions between gene groups. Y-axis was truncated (p-adj ≥ 1e-40) for visualization purpose. Right: Spatial transcript density of NegControlProbe_00041, *Bmp2*, and *Trp53bp1* showing spatially variable codeword usage. NegControlProbe_00041 variable usage is driven by codeword 5642, which shows a similar glial-cell-specific pattern as Bmp2-1345 and Trp53bp1-17048. **(F)** Cross-platform comparison of spatial variability significance. Scatter plots show HSIC-IR adjusted p-values for genes detected in both Xenium Prime 5K and either Visium-ONT (left, Spearman’s ρ = 0.19) or Visium-SR (right, Spearman’s ρ = 0.15). Red dots indicate genes identified as spatially variable in the Visium platforms. Dashed red lines mark significance thresholds. **(G)** Overlap of spatially variable processed genes across platforms. Venn diagram shows limited concordance between platforms, with Xenium-SVP filtered using stringent threshold (p-adj ≤ 2.5e-17) to minimize technical artifacts.

**Extended Data Figure 3:**
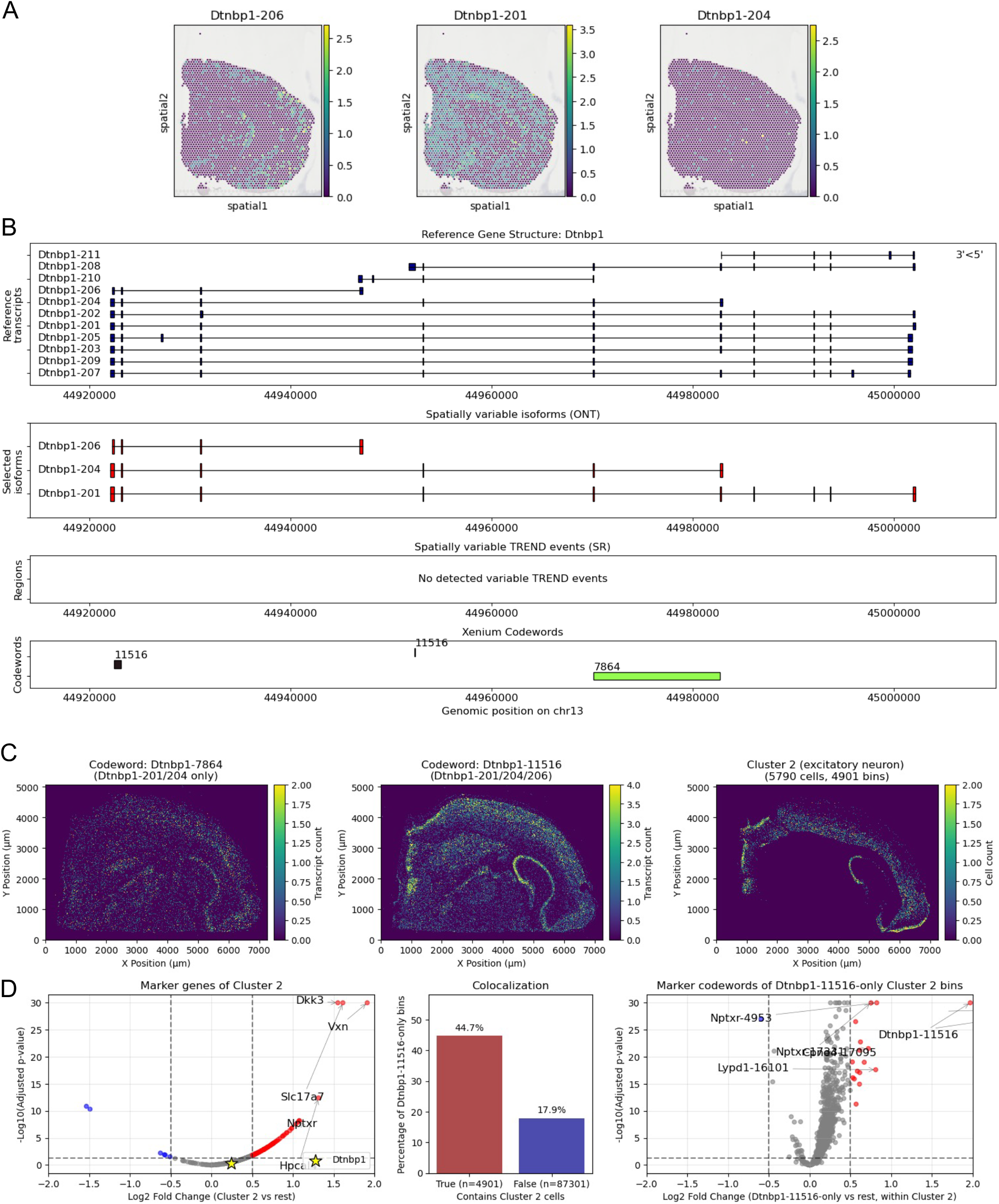
In situ validation of spatially variable isoform usage of *Dtnbp1*, related to Extended Data Figure 2. **(A)** Spatial distribution of log-normalized expression for *Dtnbp1* isoforms in SiT-ONT CBS2 sample. **(B)** Gene structure of *Dtnbp1* with annotated isoforms and Xenium Prime 5K probe sets. Among three ONT-detected *Dtnbp1* isoforms (201, 204, 206), codeword 7864 specifically targets 201 and 204, while codeword 11516 recognizes all three. **(C)** Spatial density of *Dtnbp1* codewords (left) and Cluster 2 cells (*Slc17a7*+ excitatory neurons, right) in Xenium Prime 5K mouse CBS sample. Cell segmentation, clustering and differential expression analysis were performed using Xenium Ranger with default parameters. **(D)** Dtnbp1-206 spatial variability driven by a subset of *Nptxr*+ excitatory neurons. Left: Volcano plot of Cluster 2 marker genes. Middle: Comparison of Dtnbp1-11516-only bins (20 × 20 μm) with and without Cluster 2 cells. Right: Volcano plot of marker codewords of Dtnbp1-11516-only Cluster 2 bins.

**Extended Data Figure 4:**
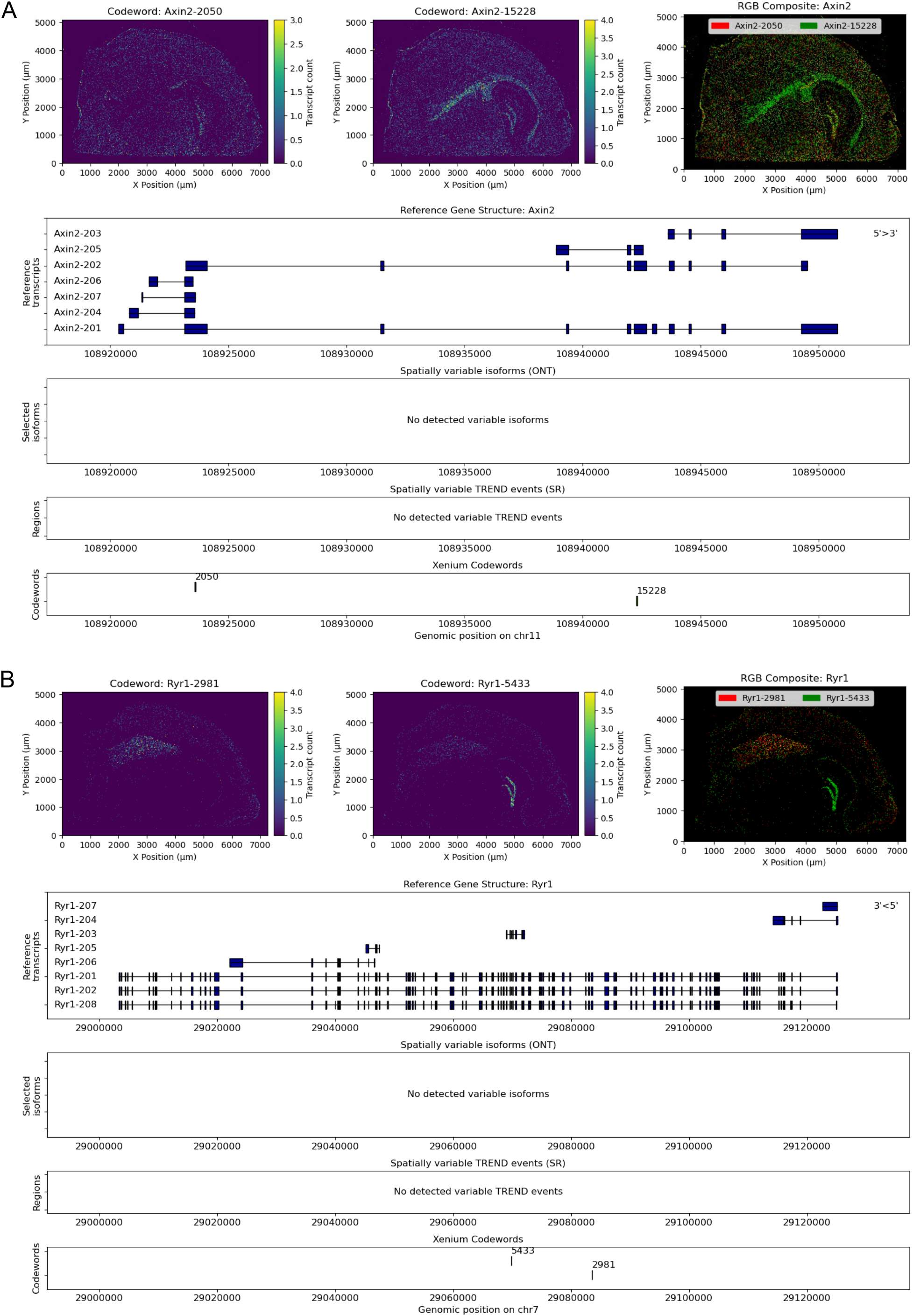
Examples of Xenium-only spatially variably processed genes identified in the mouse CBS dataset, related to Extended Data Figure 2.

### RBFOX and CELF families regulate neural transcript diversity cooperatively

Causal inference from observational studies faces drawbacks from unknown confounding factors. Since SPLISOSM’s differential test captures both direct and indirect associations in the general form, it is not immune to such confounding effects. By comparing spatial RBP associations with perturbation studies in specific cellular environments, we can nevertheless extract meaningful biological insights.

A revealing example is the spatial regulation of *Clta* isoforms (Extended Data Figure 5A). Knockout studies demonstrated that RBFOX proteins suppress *Clta* exons 5 and 6 inclusion in developing motor neurons^49^, yet our spatial test revealed a surprising positive correlation between *Rbfox3* expression and the Clta-205 isoform containing these exons. Analysis of exon inclusion in a single-cell neocortex atlas initially confirmed this positive relationship across brain cell types (Extended Data Figure 5B). The perceived contradiction resolved, however, when examining only neuronal subtypes: within GABAergic and far-projecting glutamatergic neurons, *Rbfox3* negatively correlates with exon inclusion, aligning with the knockout results. This context-dependent relationship implies the presence of other cell-type-specific factors, countering RBFOX, that drive up *Clta* exon 5 usage in neurons. We found several additional genes (*Myl6, Cltb, Tln1, Klc1*; Figure S10) where RBFOX knockout effects contradicted spatial associations. Supporting our hypothesis, previous experiments have shown that repressive CELF binding can override RBFOX enhancement at *MYL6* exon 6 in human T cells^53^.

To systematically identify potential RBFOX co-regulators, we assessed agreement between RBP differential usage results and spatial co-expression patterns (Extended Data Figure 5C). The top candidates included RBPs with varying reported RBFOX interactions: CELF family proteins, known to act antagonistically with RBFOX in muscle and heart tissue^53^; ELAVL2, sharing targets with RBFOX1 in human neurons^54^; and QKI, predicted to affect RBFOX function^55^. Expression of these collaborating RBPs remained stable after RBFOX depletion (Figure S10A), pointing toward direct cooperation rather than cross-regulation feedback. Sequence motif analysis provided further validation, showing significant enrichment of ELAVL, CELF, and QKI binding motifs near Rbfox-associated variable exons detected in the ONT data, while KHDRBS3 (a QKI homolog) and ELAVL motifs were enriched near Rbfox-associated TREND regions in short-read data (Extended Data Figure 5D). Consistently, we observed position-dependent overrepresentation of CELF and ELAVL motifs around exons responding to RBFOX triple knockout in the perturbation data^49^, supporting the collaborative regulation between multiple RBPs on the same targets (Extended Data Figure 5E).

### Evolutionarily conserved transcript diversity in synaptic genes across the human prefrontal cortex

Given the critical role of splicing and APA in brain functions, we hypothesized that spatial transcript diversity patterns would be highly preserved throughout mammalian evolution. To investigate this, we reanalyzed 12 10X Visium samples^56^ from healthy human brain (Figure 5A, Methods), identifying 861 SVP genes in the dorsolateral prefrontal cortex (DLPFC, adjusted p-value < 0.05). Power to detect spatial variability has not yet saturated with respect to sequencing depth (Figure 5B, Extended Data Figure 6A), so improved RNA capture per spot would likely reveal additional variable events. When evaluating performance across replicates, SPLISOSM’s expression-based tests (HSIC-GC and HSIC-IC) demonstrated better power and consistency compared to SPARK-X, which was hampered by suboptimal kernel design (Figure 5C). The HSIC-IR test showed weaker concordance, primarily due to data sparsity and noise, though this improved significantly for genes with higher expression.

**Figure 5:**
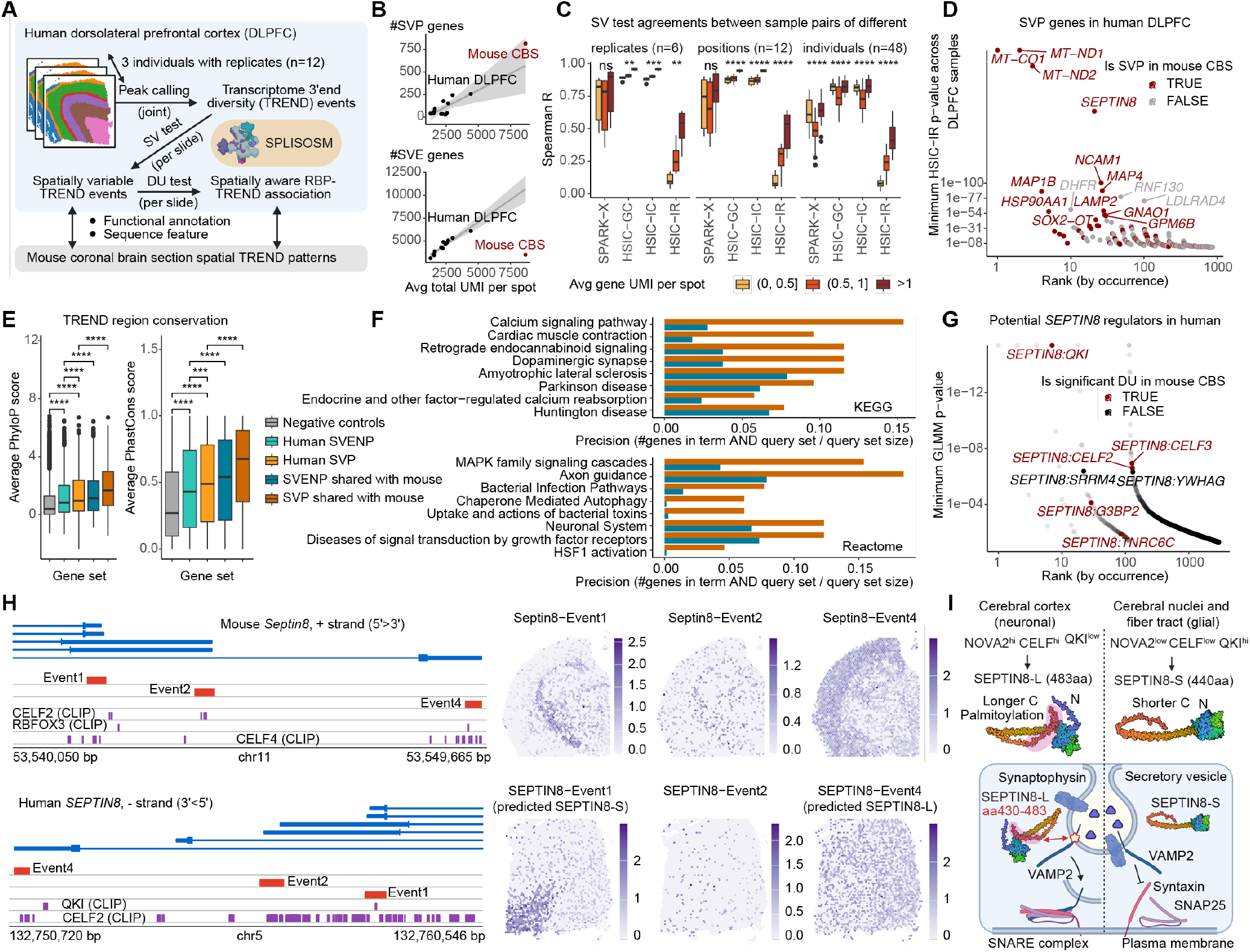
Evolutionarily conserved transcript diversity in synaptic genes across the human prefrontal cortex. **(A)** Workflow overview. **(B)** Relationship between total read coverage in TREND regions per spot (x-axis) and number of significant spatially variably processed (SVP, top, adjusted p-value < 0.01) or variably expressed (SVE, bottom, adjusted p-value < 0.01) genes across datasets. **(C)** P-value correlation distributions between technical replicates (left), different sections from same donor (middle), and different donors (right). Genes are stratified by average TREND read counts per spot and group means are compared using Kruskal-Wallis test. ns: p>0.05; **: p < 0.01; ***: p < 0.001; ****: p < 0.0001. Boxplots show median (center line), interquartile range (box), and 1.5× interquartile range (whiskers). Same below. **(D)** SVP genes ranked by recurrence (number of significant samples, x-axis) and the minimal HSIC-IR p-values across samples (y-axis). Highlighted genes have mouse homologs that are also spatially variably spliced. **(E)** Distributions of average conservation scores in TREND regions across gene categories: non-variable controls, human-specific SVENP/SVP, and human-mouse shared SVENP/SVP. Pairwise group means are compared using T-test. ***: p < 0.001; ****: p < 0.0001. **(F)** Pathway enrichment analysis comparing SVP versus SVENP genes conserved between human and mouse. **(G)** RBP-SVP associations ranked by recurrence (x-axis) and minimal GLMM p-values across samples (y-axis). Top potential regulators of *SEPTIN8* are highlighted and colored by whether the association is conserved in mouse. **(H)** *SEPTIN8* transcript structure and spatial log-normalized expression in mouse (top) and human (bottom, sample 151673). **(I)** SEPTIN8-long and SEPTIN8-short protein variants with potentially different cellular functions.

Despite these technical limitations, we found strong evidence for cross-species conservation at both individual gene and functional levels. Of the human SVP genes, 178 had mouse homologs that also exhibited spatial variability (Figure 5D, Extended Data Figure 6B-C). Moreover, conserved TREND regions displayed significantly higher sequence conservation than invariant regions (Figure 5E). Functional enrichment analyses in both species highlighted the importance of isoform-level regulation in synaptic signaling and chaperone-mediated autophagy, processes implicated in various neurodegenerative disorders (Figure 5F, Extended Data Figure 6D).

The molecular regulation governing spatial isoform diversity also showed remarkable cross-species similarities. MAP4, which organizes microtubules in muscles and neurons, exhibited patterns corresponding to neuronal-vs-glial usage in both species (Figure 4E and Extended Data Figure 6G). While PTBP2 was previously identified as a regulator of *MAP4* last exon usage in embryonic brain cortex^47^, its limited spatial variability in adult brain suggested the involvement of additional factors in region-specific regulation (Extended Data Figure 6E). Our differential usage analyses pointed to RBFOX1, QKI, and CELF families as likely co-factors, with CLIP data revealing conserved binding sites supporting this hypothesis (Extended Data Figure 6F).

SEPTIN8 provides another compelling example of evolutionarily conserved regulation. This gene encodes a GTP-binding protein that polymerizes into filaments essential for cellular architecture. Our spatial analysis revealed a distinct usage of SEPTIN8 upstream last exon, presumably producing a shorter isoform predominating in mouse fiber tracts and human DLPFC white matter (Figure 5H). This truncated protein lacks the extra C-terminal region that interacts with VAMP2 to facilitate SNARE-complex assembly^57^, which also features a filopodia-inducing motif (FIM) that enhances dendritic arborization when palmitoylated^58^ (Figure 5I). The downstream *Septin8* last exon is promoted by NOVA2 in mouse motor neurons^58^. Our analysis indicates that NOVA2 likely cooperates with QKI and CELF proteins jointly to achieve the precise spatial patterning in the brain (Figure 5G).

**Extended Data Figure 5:**
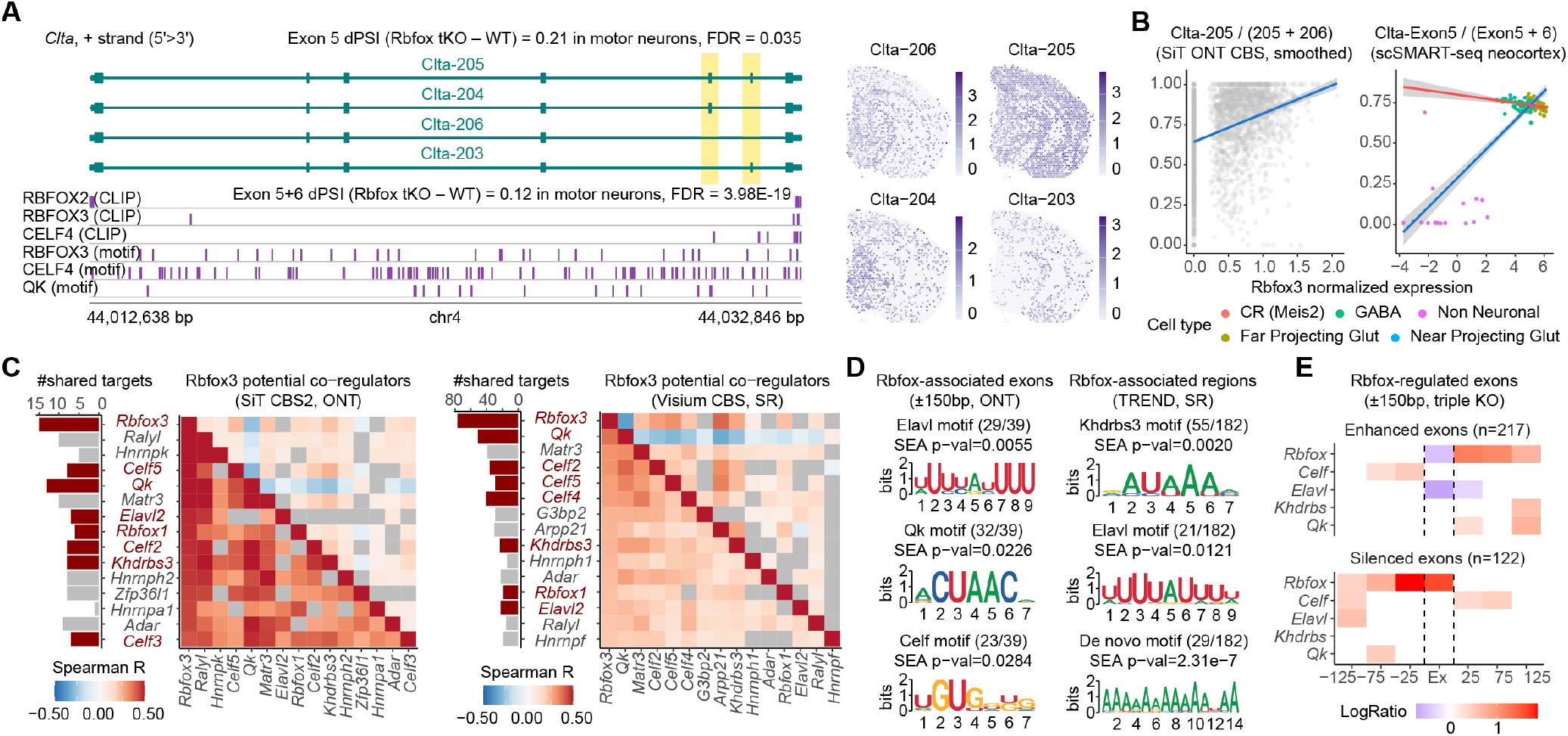
RBFOX regulates neural transcript usage with other splicing factors cooperatively. **(A)** *Clta* transcript structure and spatial log-normalized expression in the ONT-CBS2 sample. **(B)** Relationship between normalized *Rbfox3* expression and *Clta* isoform ratio (left) or cell-type-aggregated exon PSI values (right). Spatial data was locally smoothed before calculating isoform ratios for visualization. **(C)** Identification of Rbfox3 co-regulators in ONT (left) and SR (right) samples. Heatmaps display pairwise correlations of gene expression (upper triangle) or conditional HSIC test p-values (lower triangle). RBPs with reported RBFOX interaction are highlighted. **(D)** Motif enrichment in Rbfox1/2/3 associated SVP targets using alternative exons and 150bp flanking introns (left) or TREND regions (right). **(E)** Motif enrichment in exons with ≥25% PSI changes following RBFOX triple knockout in Day 10 motor neurons, categorized as silenced (increased PSI after KD) or enhanced (decreased PSI). Sequences were binned by distance to exon boundaries (x-axis) to show positional enrichment.

### Spatial transcript diversity in human glioma is shaped by immune infiltration and microenvironment composition

The capability to detect patterns without cell-type or region annotations makes SPLISOSM especially valuable for complex tissues with ambiguous cellular states. Glioblastomas are characterized by their infiltrative nature and heterogeneous spatial structures^59^. To explore isoform distributions in this challenging context, we systematically mapped spatial transcript usage across 24 human glioma samples (Figure 6A, Methods), identifying 306 spatially variably processed (SVP) genes in 11 ONT samples^60^ and 2,828 SVP genes in 13 SR samples^59^ (adjusted p-value < 0.05, Figure 6B). The most frequently recurring SR-SVP genes overlapped substantially with those found in healthy brain tissue, reflecting compositional changes of non-tumor cells (Figure 6C, Extended Data Figure 7A-B). As in healthy brain, approximately 25% of all variable transcript events in glioma had potential implications for protein structure, with alternating exon accounting for much of this variation (Extended Data Figure 7D). Beyond RNA processing, glioma-specific transcript diversity also emerged from genomic alterations. In one pediatric sample, we detected variable immunoglobulin transcripts resulting from VDJ recombination, revealing clonality in B-cell infiltration (Extended Data Figure 7C).

**Figure 6:**
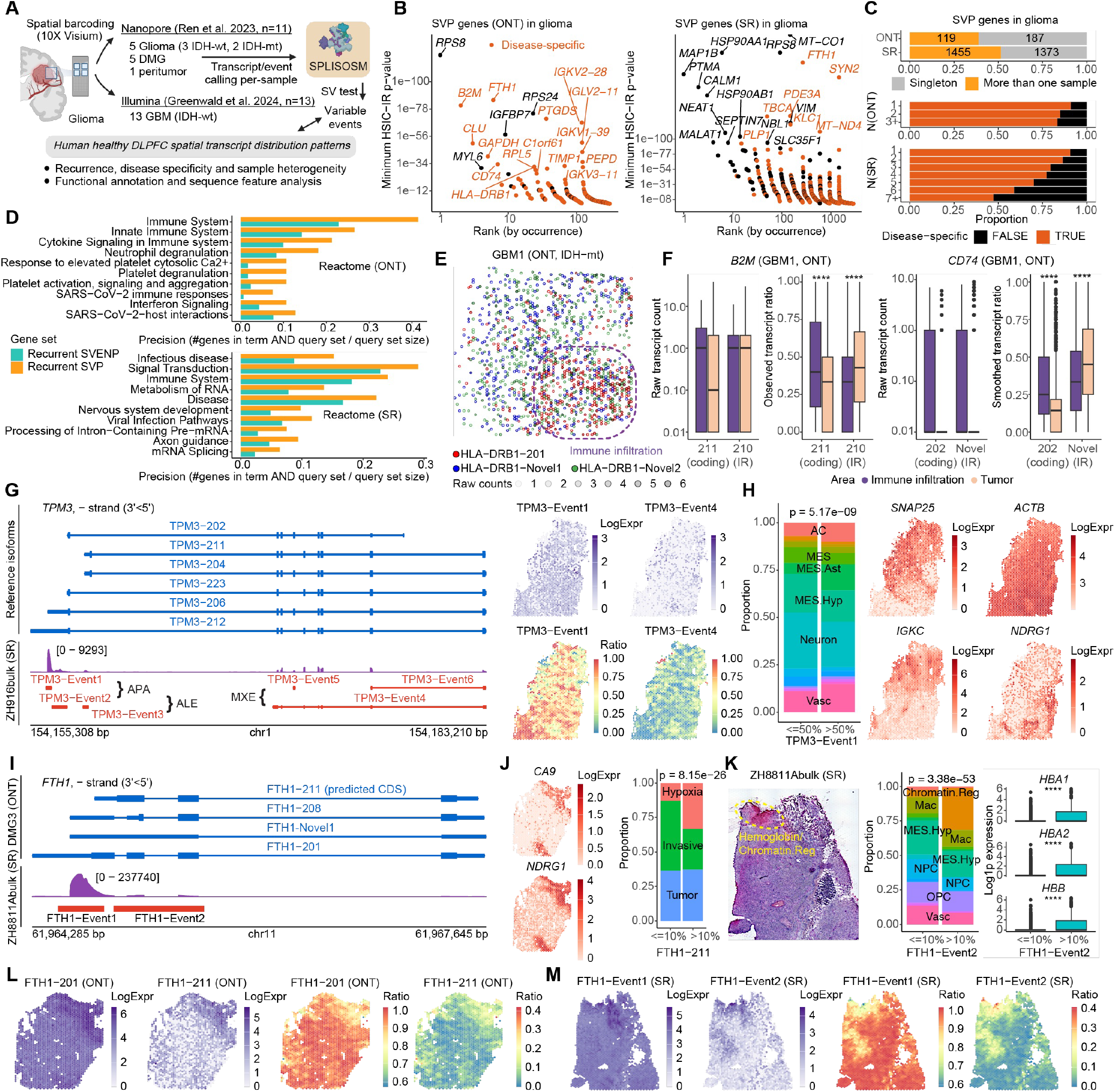
Spatial transcriptomic diversity in human glioma is shaped by microenvironment composition and immune infiltration. **(A)** Workflow overview. **(B)** SVP genes ranked by recurrence (x-axis) and the minimal HSIC-IR p-values across samples (y-axis) in ONT (left) and SR (right) cohorts. Orange indicates disease-specific genes not variable in healthy DLPFC. **(C)** Distributions of recurrent and glioma-specific SVP genes. **(D)** Pathway enrichment analysis comparing recurrent SVP (≥2 samples) genes versus recurrent SVENP (≥4 samples) genes. **(E)** Spatial distribution of *HLA-DRB1* isoforms in the ONT sample GBM1. **(F)** *B2M* and *CD74* isoform expression in GBM1, comparing tumor versus immune-infiltrated regions. Three novel *CD74* intron-retention isoforms are aggregated as one for visualization. Boxplots show median (center line), interquartile range (box), and 1.5× interquartile range (whiskers). Same below. ****: T-test p < 0.0001. **(G)** *TPM3* transcript structure, RNA-seq coverage, and spatial patterns in the SR sample ZH916bulk. Event counts were locally smoothed for ratio visualization, same in **(L)** and **(M). (H)** Glioblastoma metaprogram distribution in regions with high/low TPM3-Event1 usage. P-value from Chisquare test. Shown on the right are log-normalized expression of selected markers: *SNAP25* (neuron), *ACTB* (glial), *IGKC* (B cell), *NDRG1* (hypoxia). **(I)** *FTH1* transcript structure in ONT sample DMG3 and its RNA-seq coverage in SR sample ZH8811Abulk. **(J)** Hypoxia marker expression (left) and correlation with FTH1-211 usage in DMG3. P-value from Chi-square test. **(K)** Histology (left), metaprogram distribution (middle, p-value from Chi-square test) and hemoglobin expression (right) in regions with high/low FTH1-Event2 usage in ZH8811Abulk. Boxplot p-values from T-test: ****p < 0.0001. **(L)** Spatial distribution of *FTH1* isoforms in DMG3. **(M)** Spatial distribution of *FTH1* TREND usage in ZH8811Abulk.

The importance of immune interactions was further underscored by consistent variability in antigen presentation components across samples, often correlated with immune infiltration patterns (Figure 6D, Extended Data Figure 7E-G). This included HLA class I (*HLA-A, B, C*) and class II (*HLA-DRA, DRB1, DPB1*) genes, whose sequence variation likely stems from both alternative splicing and germline variability^61,62^ (Figure 6E). For non-polymorphic elements such as *B2M* and *CD74*, we found preferential usage of intron-retaining, potentially non-functional transcripts in tumor regions (Figure 6F). While the SR data could not capture identical variability events due to its 3’ end bias, it demonstrated similar functional convergence toward immune-related pathways, particularly in infectious disease response and viral infection pathways (Figure 6D, Extended Data Figure 7G).

Furthermore, we detected spatial variability in signal transduction, focal adhesion and cytoskeleton regulation pathways (Extended Data Figure 7H), reflecting the intricate cellular interactions and extracellular matrix remodeling within tumors. *TPM3*, a tropomyosin family member with well-documented cancer-associated splicing patterns across tumor types^63^, exhibited complex and distinct splicing variation (Figure 6G). We identified an upstream *TPM3* event (Event 4) that partially overlapped with neuronal and hypoxia signatures, while the canonical form (Event 1) associated with glial programs and increased actin expression while negatively correlating with B-cell infiltration (Figure 6H). These results resonates with recent reports of differential splicing of actin cytoskeleton components between peripheral and core glioblastoma cells^64^. Additionally, we observed spatially variable usage of the last exon of *GFAP*, encoding two variants of an astrocytic intermediate filament protein whose ratio is linked to glioma grade and malignancy^65,66^ (Extended Data Figure 7I). The location-specific distribution of these cytoskeleton gene isoforms indicates dynamic responses to local microenvironmental cues, modulating migration potentials and driving glioblastoma invasion.

Microenvironmental influences extended to ribosomal proteins as well. We observed spatial variability in transcripts encoding ribosomal proteins (including *RPS8, RPS24, RPL5, RPLP1, RPS6, RPS12*) associated with tumor infiltration, frequently involving retained introns or truncated 3’ ends (Figure S11). While some such as *RPS8* also exhibited variability in normal brain, the altered read distributions clearly displayed glioblastoma-specific changes (Figure S11A), aligning with prior findings that ribosomal isoform switching occurs in response to microenvironmental cues and contributes to metabolic reprogramming phenotypic plasticity in glioblastoma^67^.

**Extended Data Figure 6:**
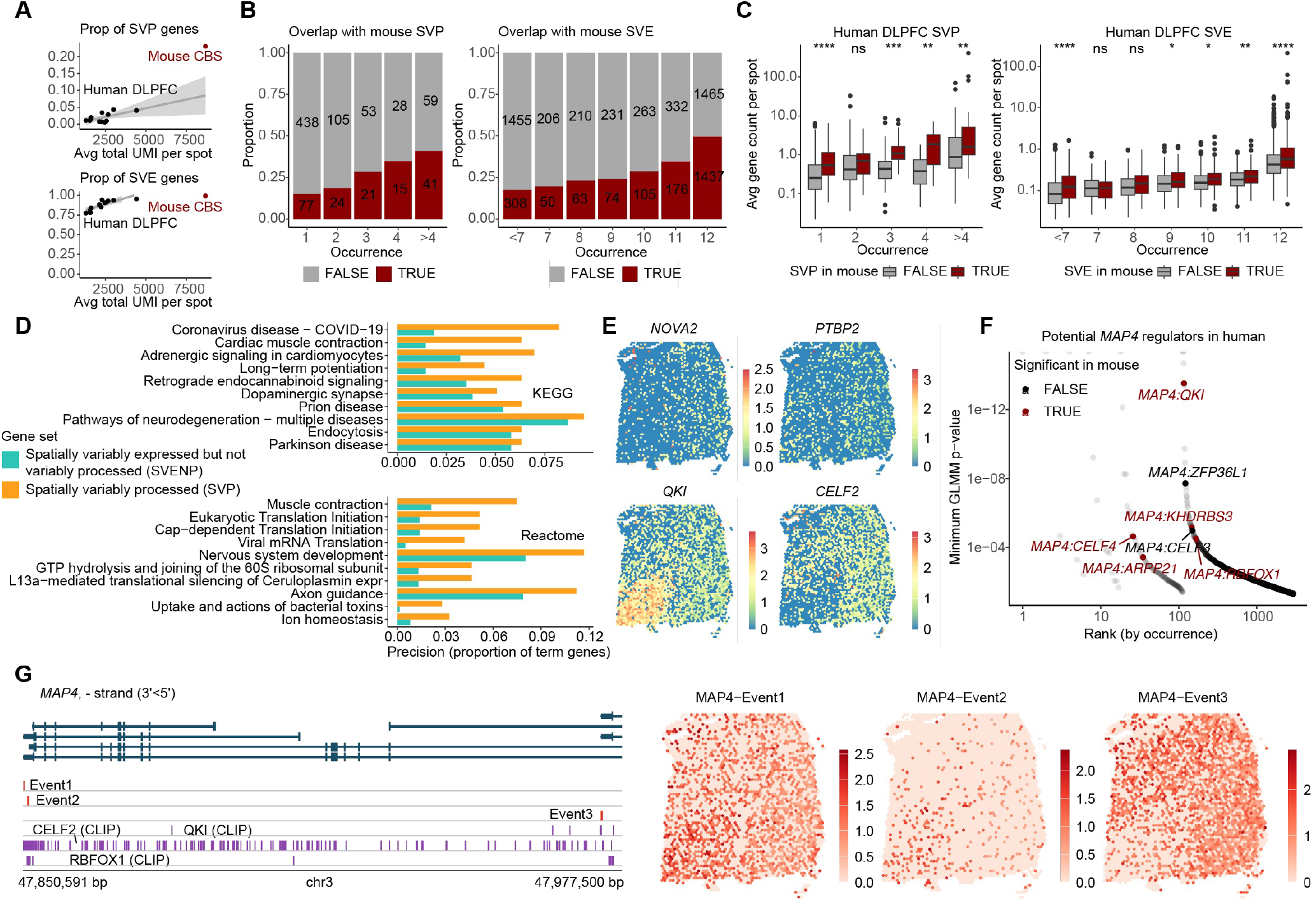
Additional examples of evolutionarily conserved SVP events in human DLPFC, related to Figure 5. **(A)** Relationship between total read coverage in TREND regions per spot (x-axis) and proportion of significant spatially variably processed (SVP, top) or variably expressed (SVE, bottom) genes in all tested genes. Related to Figure 5B. **(B)** Number and proportion of human DLPFC SV genes that have spatially variable mouse homologs, grouped by recurrence (number of significant DLPFC samples, x-axis). **(C)** Sequencing depth of SVP and SVE genes, grouped by conservation and recurrence. Boxplots show median (center line), interquartile range (box), and 1.5× interquartile range (whiskers). Group means are compared using T-test. ns: p>0.05; **: p < 0.01; ***: p < 0.001; ****: p < 0.0001. **(D)** Pathway enrichment analysis comparing human SVP versus human SVENP genes. Related to Figure 5F. **(E)** Spatial log-normalized expression of selected RBPs in sample 151673. **(F)** RBP-SVP associations ranked by recurrence (x-axis) and minimal GLMM p-values across samples (y-axis). Top potential regulators of *MAP4* are highlighted and colored by whether the association is conserved in mouse. **(G)** *MAP4* transcript structure and spatial log-normalized expression in human sample 151673.

**Extended Data Figure 7:**
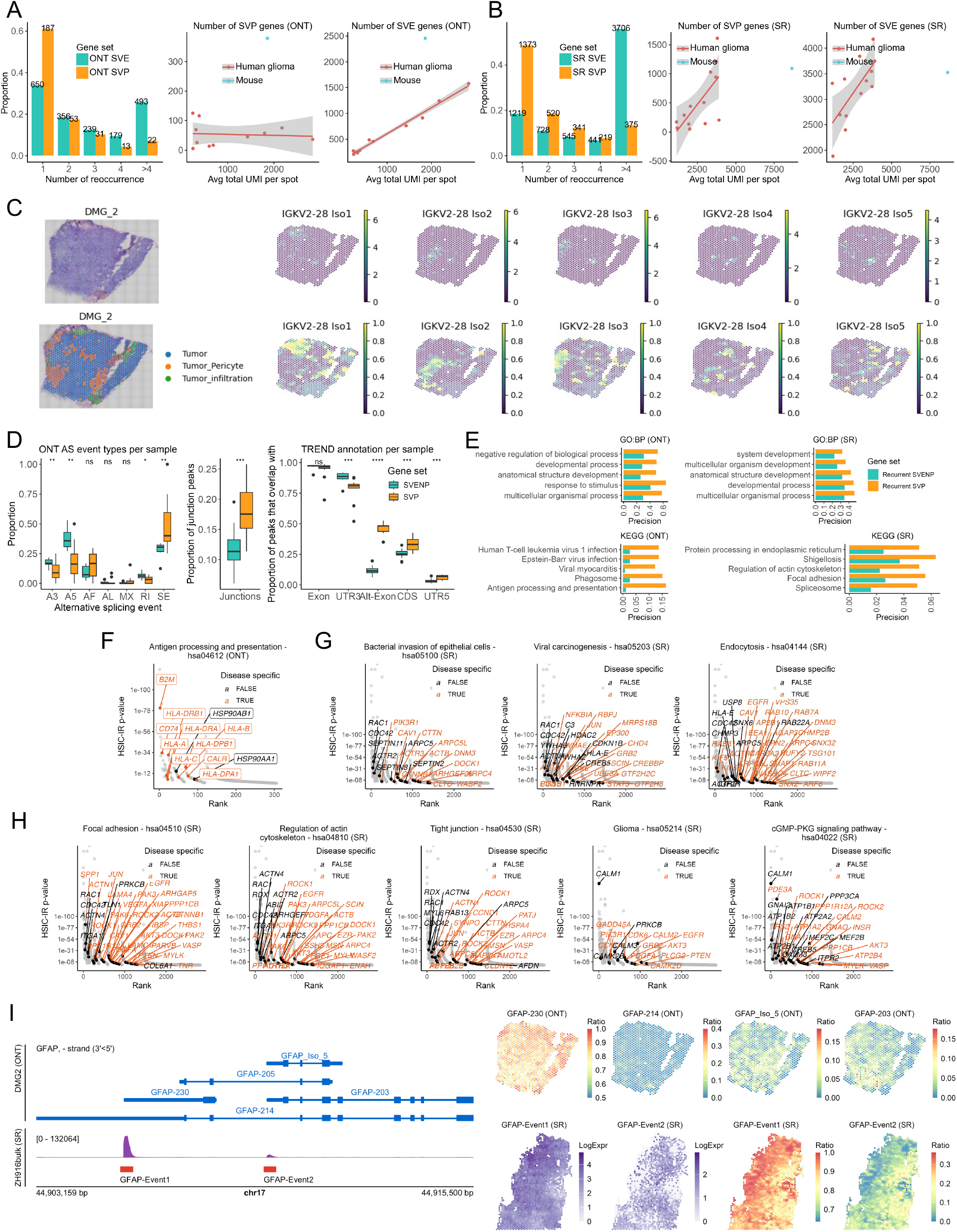
Additional examples of transcript diversity in human glioma, related to Figure 6. **(A)** Distribution of recurrent SVP genes (left) and relationships between total isoform read coverage per spot (x-axis) and number of SVP and SVE genes (y-axis) across ONT datasets. **(B)** Distribution of recurrent SVP genes (left) and relationships between total TREND read coverage per spot (x-axis) and number of SVP and SVE genes (y-axis) across SR datasets. **(C)** IG gene isoform diversity in the ONT sample DMG2. **(D)** Distributions of alternative splicing types in ONT-SV genes and TREND annotations in SR-SV genes. Boxplots show median (center line), interquartile range (box), and 1.5× interquartile range (whiskers). Group means are compared using T-test. ns: p>0.05; **: p < 0.01; ***: p < 0.001. **(E)** Pathway enrichment analysis comparing recurrent SVP versus recurrent SVENP genes. Related to Figure 6D. **(F-H)** SVP genes involved in selected KEGG pathways. **(I)** *GFAP* transcript structure in the ONT sample DMG2, read coverage in the short-read sample ZH916bulk, and the respective isoform or TREND spatial distribution in each sample (right).

*FTH1* offers another prominent example of microenvironment-driven transcript usage (Figure 6I-M). This iron-storage protein, upregulated under hypoxic conditions, protects cells from ferroptosis by reducing reactive iron^68^. We discovered that *FTH1* upregulation in hypoxic tumor regions coincided with increased expression of 3’ truncated transcripts lacking the final exon-exon junction (Figure 6J). In short-read data, we identified a minor event spanning *FTH1* exons 2 and 3, predominantly in regions with erythrocytes and overlapped with the chromatin regulation metaprogram (Figure 6K). Our findings suggest coordination between oxygen-sensing, iron metabolism, and stress response pathways through transcript-level modulation of FTH1.

## Discussion

Spatial omics techniques are transforming our understanding of tissue organization, yet few studies have explored molecular diversity at the isoform level. Here we present SPLISOSM, a computational framework designed for robust detection of spatial transcript processing and its regulation in ST data. Through theory-grounded non-parametric kernel tests, our approach achieves high statistical power in sparse data while delivering well-calibrated, permutation-free p-values. SPLISOSM demonstrates broad applicability across ST platforms and spatial resolutions. We show that standard 3’ end sequencing protocols and predesigned in situ panels can capture considerable transcript diversity, enabling researchers to extract novel insights from existing datasets. Our implementation of low-rank kernel approximation allows testing hundreds of thousands of locations within minutes on standard hardware with minimal power loss (Supplementary Note).

Using SPLISOSM, we have uncovered thousands of evolutionarily conserved spatial transcript diversity events in mouse and human brain. However, the spectrum of detectable events varies across ST platforms (Figure 1A). While SPLISOSM remains agnostic to event type, its performance depends on platform-specific limitations (Extended Data Figure 8). Sequencing-based platforms suffer primarily from false negatives due to 3’ bias, read length, and sequencing depth, particularly affecting detection of longer transcripts (Extended Data Figure 4B). Current depths have not saturated SVP detection, suggesting that improvements in RNA capture efficiency and isoform quantification especially for novel transcripts can further expand the landscape. In contrast, imaging-based technologies face false positives from off-target probe binding, exemplified by spurious glial-specific patterns in negative control probes (Extended Data Figure 2E). Nevertheless, our analysis revealed hundreds of novel variable exon usage events in mouse brain using predesigned Xenium 5K panel, though mapping these events to full-length isoforms requires future efforts.

**Extended Data Figure 8:**
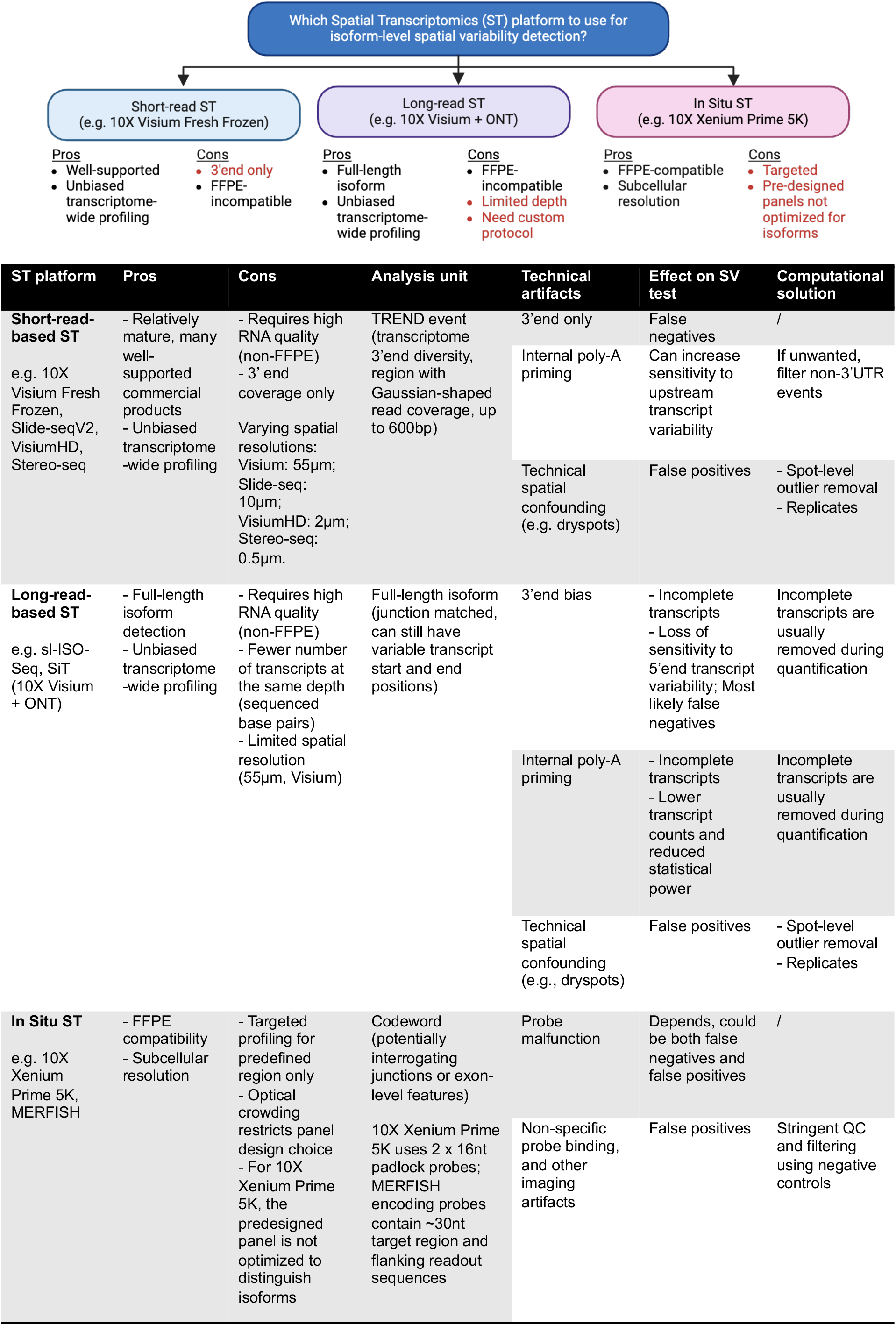
Comparison on the applicability of common ST platforms for isoform variability detection.

Our integrative regulatory analyses shed light on the complex network governing spatial splicing. Neural RBPs, including RBFOX and CELF family members, may orchestrate transcript diversity through precise spatiotemporal activation^69^. However, important limitations temper our conclusions. First, ST data sparsity limits association test power, potentially missing lowly expressed but functionally important RBPs. Second, tissue and developmental stage mismatches between spatial data (adult mouse brain) and available functional perturbation data (often in vitro cell culture) complicate direct validation. Evidence for cooperative regulation derived from co-expression, motif enrichment, and CLIP binding remains largely correlative. Future perturbation experiments in appropriate neural contexts are essential to validate and elucidate the molecular basis of the proposed cooperative mechanisms. Third, while our differential usage test accounts for spatial autocorrelation, they still assess correlation rather than causation and treat RBPs independently without explicitly controlling for co-expression. Future extensions could incorporate causal graphical models to test conditional independence among multiple regulators.

In glioblastoma, our analysis reveals how spatial transcript diversity is shaped by microenvironmental composition and metabolic conditions. The upregulation of truncated and intron-retaining transcripts in antigen presentation genes implies that tumors may employ alternative processing for immune evasion^62^. Targeting these events may enhance immune response and improve immunotherapy benefits. We also observed significant diversity in ribosomal transcripts, suggesting translational regulation could play an underappreciated role in tumor heterogeneity and thus exposing new therapeutic vulnerabilities^67^.

In summary, SPLISOSM introduces an isoform-centric perspective to ST data analysis. The spatial organization of transcript diversity we mapped in healthy and malignant brain tissues established a foundation for understanding how alternative isoform processing contributes to specialized functions and disease pathology.

## Supporting information

Supplementary Table 1: Datasets and sample metadata

## Acknowledgements

We thank Chaolin Zhang for input on the splicing regulatory analysis and Daniel Moakley for assistance on the preprocessing of single-cell exon splicing data. We thank Junwei Song for access to processed 10X Visium ONT glioblastoma data and transcript annotation. We thank Jun Hou Fung for feedback on the theoretical analysis. We thank Sanja Vickovic, David McKellar, and Karin Isaev for helpful discussions. This work was funded by the National Institutes of Health, National Cancer Institute (R35CA253126, U01CA243073, P01CA285250 to R.R.), the Edward P. Evans Center for MDS at Columbia University (to J.S. and R.R.), and the National Science Foundation (CAREER DBI2146398 to D.A.K.). Any opinions, findings, and conclusions or recommendations expressed in this material are those of the authors and do not necessarily reflect the views of the NSF or the NIH.

## Contributions

J.S. conceived the project, designed and implemented SPLISOSM, conducted the theoretical analysis, benchmark experiments, and applications on spatial transcript diversity in mouse and human samples. Y.Q. contributed to the parallelization and benchmark of the parametric models; J.S., M.S., and S.P. designed and conducted experimental validation; H.Y. applied SPLISOSM to the Slide-seqV2 hippocampus data; J.J. processed the short-read glioblastoma samples; J.S. and T.L. developed the website; Q.L. contributed to the analysis of 3’ end variability in neurodegenerative diseases; T.M.N., X.F. and S.C. contributed to the analysis and interpretation of transcript diversity in glioblastoma samples; D.A.K. and R.R supervised the project. J.S took lead in writing the manuscript with input from all authors. All authors read and approved the final manuscript.

## Data availability

This paper analyzes existing, publicly available data without generating new primary data.

References and accession details to raw data are provided in the Supplementary Table 1 and in the corresponding Methods sections. Per-cohort isoform quantification results and other data necessary to reproduce results described in this study can be downloaded from Zenodo (10.5281/zenodo.15556390). Processed and formatted per-sample data and SPLISOSM test results can be downloaded from Zenodo (10.5281/zenodo.16905935).

## Code availability

The SPLISOSM python package is available at https://github.com/JiayuSuPKU/SPLISOSM under BSD-3 License. Code and scripts to reproduce this paper, as well as Google Colab notebooks for interative data exploration and visualization, are available at https://github.com/JiayuSuPKU/SPLISOSM_paper.

## Methods

### SPLISOSM Overview

SPLISOSM analyzes isoform-level spatial transcriptomics data through a two-step approach: first detecting spatial variability (SV), then identifying differential usage (DU) by associating isoform patterns with potential regulatory factors. The framework accommodates quantification results of both full-length isoforms from long-read sequencing and local transcript diversity events from short-read sequencing platforms including 10X Visium and Slide-seqV2.

Unlike gene-level analysis, SPLISOSM recognizes that multiple isoforms (*q* > 1) from the same gene are interdependent. Their relative proportions form a multivariate statistical variable across the spatial random field of *n* locations

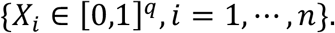

Together with the 2D spatial coordinates {*Y*_*i*_ ∈ ℝ^2^, *i* = 1, …, *n*} and covariates such as RNA binding protein expression {*Z*_*i*_ ∈ ℕ, *i* = 1, …, *n*}, we can formulate spatial variability detection and differential association analysis as multivariate independence tests T(*X, Y*) and T(*X*|*Y, Z*|*Y*). The spatial conditioning in DU tests is critical for eliminating spurious associations caused by spatial autocorrelated noise and hidden confounders.

Below we outline SPLISOSM’s key components, with comprehensive methodology, theoretical analysis and implementation details available in the Supplementary Note.

### Unconditional kernel independence test

A commonly used statistic for multivariate correlation analysis is the RV coefficient, defined as the normalized matrix norm of the q-by-2 cross-covariance matrix *C*_*X*F_ = *C*_*ij*_ = Cov;*X*^*i*^, *Y*^*j*^< between X and Y

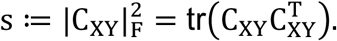

To capture complex, non-linear spatial patterns beyond linear correlation, SPLISOSM employs kernel transformations to map variables into reproducing kernel Hilbert spaces (RKHSs). In these potentially infinite-dimensional feature spaces, we measure statistical dependence using the Hilbert-Schmidt Independence Criterion (HSIC)^33^, defined as the Hilbert-Schmidt norm of the cross-covariance operator *C*_*X*F_: 𝒢 → ℱ that connects the respective feature spaces ℱ and 𝒢 of X and Y,

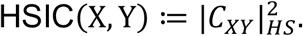

Similar to the univariate gene expression SV test SPARK-X^30^, we build calibrated, permutation-free testing procedures based on the empirical HSIC estimator,

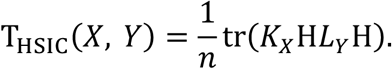

Here, *K*_*X*_ and *L*_*Y*_ are the n-by-n isoform and spatial kernel matrices that specify feature maps for X and Y, and 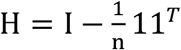 is the centering matrix. Under the null hypothesis of statistical independence, *T*_*HSIC*_ follows an asymptotic distribution of chi-square mixture^34^, with weights derived from the eigenvalues {*λ*_*i*_} and {*μ*_*j*_} of *K*_*X*_ and *L*_*Y*_,

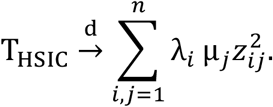

The statistical power of HSIC-based non-parametric test depends heavily on effective kernel design. To profile spatial isoform patterns in sparse data, SPLISOSM introduces two technical innovations:

First, we prove that any low-rank approximation of the spatial kernel significantly reduces test power (Theorem 1, Supplementary Note), highlighting a key limitation of existing SV tests like SPARK-X. Based on this insight, we developed a new full-rank spatial kernel using the Intrinsic Conditional Autoregressive (ICAR) model^35^,

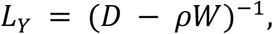

where *W* is the n-by-n binary adjacency weight matrix of the mutual k-nearest-neighbor spatial graph, *D* = diag{Σ_*j*_ *W*_*ij*_} is the degree matrix, and ρ ∈ [0,1) is the spatial autocorrelation coefficient ensuring invertibility.

In essence, this approach represents spatial patterns as signals on the spatial graph to be decomposed onto the eigenvectors of *L*_*Y*_ through Graph Fourier Transform (GFT)^70^, allowing us to prioritize patterns by frequency in the graph spectral domain without manually selecting spatial bandwidths. Additionally, the sparse structure of *D* and *W* enhances computational efficiency. For large-scale datasets (e.g. hundreds of thousands of spots) where full-rank spatial kernels become computationally prohibitive, we developed GFT-based low-rank kernels that approximate the kernel inverse *L*^−1^ through sparse eigen-decomposition, preserving biologically meaningful low-frequency patterns while limiting power loss to less important high-frequency signals.

Second, to handle sparse isoform data where many spots have zero gene coverage, we designed a Zero-Padded Centered (ZPC) kernel for isoform ratios *X*_*n*_ with *n* − *m* undefined values,

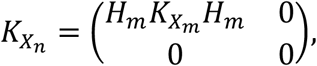

where *X*_*m*_ represents isoform ratios at *m* spots with non-zero gene coverage, and 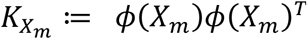 is the isoform kernel with transformation *ϕ* for compositional data (e.g. log-ratio transformations). Our empirical analysis showed that a simple linear kernel *k*(*x, y*) = ⟨*x, y*⟩ (with identity transformation) delivers optimal performance.

The ZPC version of a linear kernel is equivalent to a two-step procedure where the *n* − *m* undefined ratios in *X*_*n*_ are first replaced with global averages per isoform before constructing the n-by-n isoform kernel for independence test. The NA replacement allows us to reuse the n-by-n spatial kernel across genes with varying sparsity patterns, substantially improving computational efficiency. We mathematically prove that this approach provides a bounded approximation to testing procedures that simply omit spots with undefined ratios (Theorem 2, Supplementary Note).

Detailed implementations of the ICAR kernel and SPLISOSM’s three spatial variability tests --- HSIC-GC (gene expression), HSIC-IR (isoform usage ratio), and HSIC-IC (isoform expression) --- are available as Algorithms 1-4 in the Supplementary Note. The GFT-based spatial kernel construction is presented in Algorithm 5.

### Conditional kernel independence test

Linking isoform usage X with potential regulators Z requires addressing a fundamental challenge: X and Z may appear related simply because both follow similar spatial patterns irrelevant to splicing regulation. To learn the causal relationship, we need to condition on spatial coordinates Y and all other confounders if available. However, ST data provide only snapshots: at each location *Y*_*i*_, we observed only one (*X*_*i*_, *Z*_*i*_) pair, making direct estimation of conditional distributions for *X*|*Y* and *Z*|*Y* impossible.

We address this using the conditional independence framework from Zhang et al^34^, which shares information across observations by learning regression functions *f*_*X*_: *Y* → *X* and *f*_*Z*_: *Y* → *Z*, then testing kernel independence of the residuals.

For linear ridge regression with L2 regularization λ, we can compute the residuals as

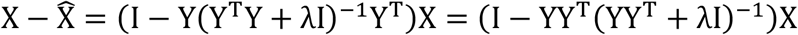

where *Y* ∈ *R*^*n*×2^ represents spatial coordinates and *X* ∈ *R*^*n*×d^ the spatial variable to predict. By substituting the linear kernel *YY*^*T*^ with general spatial kernels *K*_*Y*_, we extend the solution to non-linear relationships

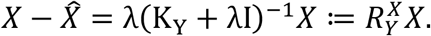

This allows us to construct a kernel for the conditional variable *X*|*Y* as 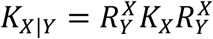 where 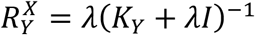 is learned from the data. The parameter *λ* controls conditioning strength: at *λ* = 0, X is fully explained by coordinates Y, leaving no information for *X*|*Y*; as λ → ∞, the residuals 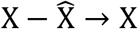, which gives the unconditional test. Since genes have diverse spatial patterns, we estimate *λ* and *K*_*Y*_ for each variable X (and Z) individually using Gaussian Process Regression.

Using linear kernels *K*_*X*_ and *K*_*Z*_, our conditional association test statistic becomes

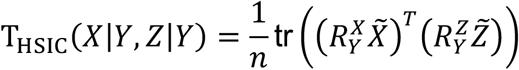

where 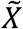 *and* 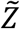 are column-wise centered isoform usage ratios and covariates, and 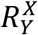 and 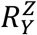 are the residual operators estimated from Gaussian Process regression.

Detailed implementations of the HSIC-based unconditional and conditional DU tests are provided as Algorithms 6 and 7 in the Supplementary Note. For parametric approaches, we can also introduce spatial conditioning through generalized linear mixed models (GLMMs) with spatial random effects. Specifically, we implemented the score test for differential usage where null GLMM models without covariates are first fitted to account for background spatial patterns. Covariate significance is then assessed by computing the gradient (i.e. score statistic) at zero effect size. Details on model fitting and hypothesis testing are available in Chapter 3 of the Supplementary Note.

### General test configurations

For all isoform spatial variability tests, we used raw UMI counts from isoform quantification results (i.e. isoform counts in ONT data and TREND event counts in SR data). Input data varied by method: HSIC-GC and SPARK-X used summed isoform UMI counts (a scalar) per gene per spot, HSIC-IC used the vector of isoform counts per gene per spot, and HSIC-IR used observed isoform ratios. For spots with zero gene coverage (undefined ratios), we substituted with the global mean ratio (justified in Theorem 2).

For differential usage testing with RBP expression, we used log1p-normalized Visium RBP gene expression (library size = 1e-4) as covariates. We first identified spatially variably expressed RBPs (adjusted p-values < 0.01 in both HSIC-GC and SPARK-X tests) to include as covariates, excluding ribosomal proteins and non-canonical RBPs. SVE tests were conducted on Visium-based RBP gene raw counts. For human DLPFC samples, we restricted our analysis to potential splicing modulators (RBPs with motif or CLIP binding data). Only SVP genes with average total ONT/TREND counts >0.5 per spot were investigated for DU with RBP. Specific numbers of tested RBPs and SVP genes per sample are detailed in Supplementary Table 1. For HSIC-based DU tests, we used observed isoform ratios with mean replacement for undefined values. Gaussian process regression was implemented using GaussianProcessRegressor from the scikit-learn^71^ package with learnable RBF and white noise kernels. For parametric models (GLM/GLMM), we used isoform count vectors as inputs, grouping genes with identical isoform numbers to enable parallel processing. We calculated p-values using score tests on fitted null models under asymptotic chi-square distribution.

Except otherwise noted, all SPLISOSM SV results were computed with the full-rank ICAR kernel with *ρ* = 0.99 and the number of mutual nearest neighbors *k* = 4 for constructing the spatial graph. SV results in the Xenium Prime 5K adult mouse brain sample were computed with the low-rank kernel (r=100) for computational efficiency. HSIC-based test p-values were calculated using the Liu’s method^72^ to approximate the chi-square mixture null distribution. Multiple testing correction was applied using the Benjamini-Hochberg method^73^.

### External datasets of RNA binding proteins

We obtained mouse and human RBP lists from EuRBPDB^74^. For motif analysis, we obtained RBP binding motifs from CISBP-RNA^75^ and refined them using the R package universalmotif^76^ to trim low-information peripheral positions before scanning with FIMO from the MEME suite^77^ (p-value threshold = 1e-3, background model computed using the default fasta-get-markov). RBP CLIP data was downloaded from the POSTAR3^40^ database and the ENCODE^78^ portal (for PCBP2 human eCLIP). In our downstream analysis, we prioritized RBPs with validated binding motifs or CLIP evidence as potential splicing regulators. APA-related RBPs were collected based on regulators previously characterized in a high-throughput screening study^79^.

### Data simulation

We generated *in silico* spatial isoform quantification data using a hierarchical generative model following two steps: first sampling total gene expression and expected isoform usage per spot, then sampling observed isoform counts from spot-specific multinomial distributions.

For total gene expression, we sampled from Poisson distributions with spot-specific means. In Scenarios 1-3 (without spatial gene expression variability), these means were drawn from independent Gaussian distributions (*μ* =5, *σ*^2^ =1). In Scenarios 4-6 (with regional gene expression patterns), means were sampled from multivariate Gaussian distributions with binary covariate-controlled means and covariance matrices combining spatial autocorrelation (ICAR)^35^ and white noise.

For expected isoform usage ratios, we sampled in the log-ratio space with *q* − 1 degrees of freedom where *q* = 3 is the number of isoforms per gene. In Scenarios 1 and 4 (without spatial isoform variability), log-ratios were sampled from independent Gaussian distributions (*μ*=0, *σ*^2^=0.2) and transformed via softmax to obtain expected ratios. In Scenarios 2 and 5 (with Gaussian process variability), log-ratios were sampled from multivariate Gaussian distributions with ICAR-based spatial covariance. In Scenarios 3 and 6 (with regional isoform usage), log-ratios were determined by binary covariates with added ICAR-based spatial autocorrelated noise.

To evaluate statistical power, we created intermediate scenarios by interpolating between key configurations. For the SV test HSIC-IR, we varied the proportion of spatially autocorrelated noise between Scenarios 4 (regional expression, no isoform usage variability) and Scenarios 5 (regional expression, 50% spatially autocorrelated noise + 50% white noise in the Gaussian process). For DU tests, we adjusted covariate effect sizes between Scenarios 5 (regional expression, no isoform usage association) to Scenarios 6 (regional expression, covariate effect size=0.5).

### SiT data of adult mouse brain

We downloaded processed spatial isoform transcriptomics (SiT)^18^ isoform quantification results as well as Visium gene expression data of mouse olfactory bulb (MOB) and two coronal brain section samples (CBS1, CBS2) from Gene Expression Omnibus (GSE153859). The SiT ONT datasets contain only reference isoforms mapped to Mouse Gencode vM24. We removed isoforms expressed in <1% of spots and genes with fewer than 2 detected isoforms.

To reduce sparsity for visualization and clustering, we applied local smoothing by borrowing 20% of UMIs from six nearest neighbors per spot, then calculated isoform ratios with mean-padding for remaining undefined values. The resulting ratios may amplify noise and artifacts and should therefore be interpreted with caution. For spatial clustering, we first identified significant SVE and SVP genes using unsmoothed raw data, then performed dimensionality reduction (PCA, 50 components) and Leiden clustering on either smoothed gene expression or isoform ratios. RBP-based clustering used log1p-normalized Visium expression of SVE RBPs. Clustering resolutions were individually chosen such that the resulting cluster number were similar. For SVP usage program hierarchical clustering, we computed pairwise gene similarity using RV coefficients between multivariate smoothed isoform usage ratios.

For splicing type annotation, we used SUPPA^80^ to split isoforms into local alternative splicing events. Skipping exons (SE), mutually exclusive exons (MX), and their flanking introns were analyzed for motif enrichment using MEME suite^77^ tools (XSTREME and SEA), with additional exon sequence features extracted using Matt^81^.

### Visium and Slide-seqV2 data of adult mouse brain

We downloaded RNA-seq bam files of the 10X Visium^44^ data of a fresh frozen adult mouse coronal brain section sample from https://www.10xgenomics.com/datasets/adult-mouse-brain-coronal-section-fresh-frozen-1-standard, and the Slide-seqV2^16^ data of adult mouse hippocampus from https://singlecell.broadinstitute.org/single_cell/study/SCP815/sensitive-spatial-genome-wide-expression-profiling-at-cellular-resolution#study-summary.

Transcriptome 3’ end diversity (TREND) was quantified using Sierra^82^ with Mouse Gencode vM10 reference annotation, with minimal peak cutoffs in FindPeaks set to zero to recover minor events in the quantification stage. TREND events expressed in <1% of spots and genes with fewer than 2 TREND regions were later filtered out. Events were annotated using Sierra’s AnnotatePeaksFromGTF with annotation_correction=FALSE to allow multiple annotations per region. Alternative exons (Alt-Exon) were defined as exons located within introns of other transcript variants. Motif enrichment analysis was performed on exons of TREND regions using MEME suite^77^ tools (XSTREME and SEA).

For spatial variability tests on gene expression (HSIC-GC and SPARK-X), we used the sum of TREND reads per gene per spot rather than Visium’s Space Ranger quantification (except for the RBP SVE test which we always used the latter) to maintain consistency with HSIC-IC/IR tests and ONT data analysis. Slide-seqV2 data was analyzed using HSIC-GC/IC/IR with full-rank spatial kernels. See Supplementary Note for low-rank test performance.

For spatial clustering, we use unsmoothed TREND-based gene expression and isoform usage from top 200 SVE and SVP genes for respective clusters, and Visium-based expression of all SVE RBPs for RBP clusters. We performed dimensionality reduction (PCA, 50 components) and Leiden clustering following the same approach as in our SiT ONT data analysis.

### Xenium Prime 5K data of adult mouse brain

We obtained the Xenium output bundle of a fresh frozen adult mouse brain hemisphere profiled using 10X Xenium Prime 5K Mouse Pan Tissue and Pathways Panel^46^ from https://www.10xgenomics.com/datasets/xenium-prime-fresh-frozen-mouse-brain and reran the Xenium Ranger (v3.1.1) relabel pipeline with default parameters. Transcripts passing sequence quality filter (Q ≥ 20) from genes with multiple detected codewords were spatially binned into 20 × 20μm regions. This yielded a sparse count matrix comprising 10,476 codewords from 4,993 genes across 92,202 spatial locations. SVP significance threshold (adjusted p-value < 2.5e-17) was determined by the minimum p-value observed in negative control and intergenic probe sets. Cell segmentation, clustering, and per-cluster differential gene expression analysis were performed as part of the Xenium Ranger pipeline with default parameters. For co-localization analysis, we quantified the number of cells from different clusters within each 20 × 20μm bin. To identify molecular heterogeneity within Cluster 2 (excitatory neurons), we performed an additional differential codeword expression analysis using T-test, comparing Dtnbp1-11516-only bins against the remaining Cluster 2 bins. This analysis revealed *Nptxr* as a marker for the Dtnbp1-206-positive excitatory neuron subpopulation.

### Single cell splicing quantification and RBFOX triple knockout data

We analyzed processed exon and splicing quantification data from the adult mouse neocortex single-cell atlas^83^ using results from Moakley et al. 2024^84^. Cassette exon PSI was quantified at cell-type level using Quantas^4^ by aggregating cells of identical types. Average RBP expression per cell type was log normalized and standardized to zero means.

For RBFOX triple knockout analysis, we used processed Quantas-based splicing quantification results of mouse Day 5 and 10 motor neurons in vitro differentiated from mouse embryonic stem cells from Jacko et al. 2018^49^. The alternative splicing event, PSI and statistical significance were all determined by the original study. All RBFOX tKO events shown in the paper are statistically significant after multi-test correction. Raw RNA-seq data was downloaded from the Sequence Read Archive (SRP128054) to run RBP map and positional motif enrichment analysis using Matt^81^.

### Visium data of human DLPFC

We downloaded Visium RNA-seq bam files of 12 human dorsolateral pre-frontal cortex (DLPFC) samples^56^ through Globus (endpoint: jhpce#HumanPilot10x). Transcriptome 3’ end diversity (TREND) was quantified using Sierra^82^ with 10X’s GRCh38-2024-A reference annotation and default parameters. To create a shared set of TREND events across samples, we merged per-sample peaks using MergePeakCoordinates with default settings and quantified them with CountPeaks. We obtained conservation scores (phyloP100way and phastCons20way) from the UCSC Genome Browser and calculated average conservation scores specifically for exons within the TREND regions.

### ONT and Visium data of human glioma samples

We analyzed two independent cohorts of human glioma samples using long-read and short-read spatial transcriptomics platforms.

For the Ren et al. 2023 cohort^60^, we obtained processed ONT isoform quantification results, matched Visium gene expression data, and region annotations for 13 human glioma samples directly from the study authors, as the raw data was restricted by human genetic resources regulations. ONT transcripts were error-corrected, collapsed by shared exon-exon junctions, and quantified using DEMINERS^85^ by the original study. For spatial analysis, we included only reference isoforms matching Gencode v32 (hg38) and novel isoforms from known genes. We further filtered out isoforms expressed in <5% of spots and genes with fewer than 2 detected isoforms. SUPPA^80^ was used to categorize alternative isoform usage into local alternative splicing events.

For the Greenwald et al. 2024 cohort^59^, we processed Visium data from 13 IDH wild-type glioblastoma samples downloaded from SRA (PRJNA994130). Raw RNA-seq reads were aligned to the human reference genome (hg38) using Space Ranger v3.0.1. Metaprograms and spatial annotations were obtained from the authors’ GitHub repository (github.com/tiroshlab/Spatial_Glioma). Transcriptome 3’ end diversity (TREND) was quantified using Sierra^82^ with 10X’s GRCh38-2024-A reference annotation and default parameters. Due to sample heterogeneity, we processed each sample independently with sample-specific TREND regions rather than merging across samples. For spatial analysis, we filtered out TREND events expressed in <1% of spots, genes with fewer than 2 TREND regions, and novel genes with symbols beginning with ‘ENSG’.

**Figure S1:**
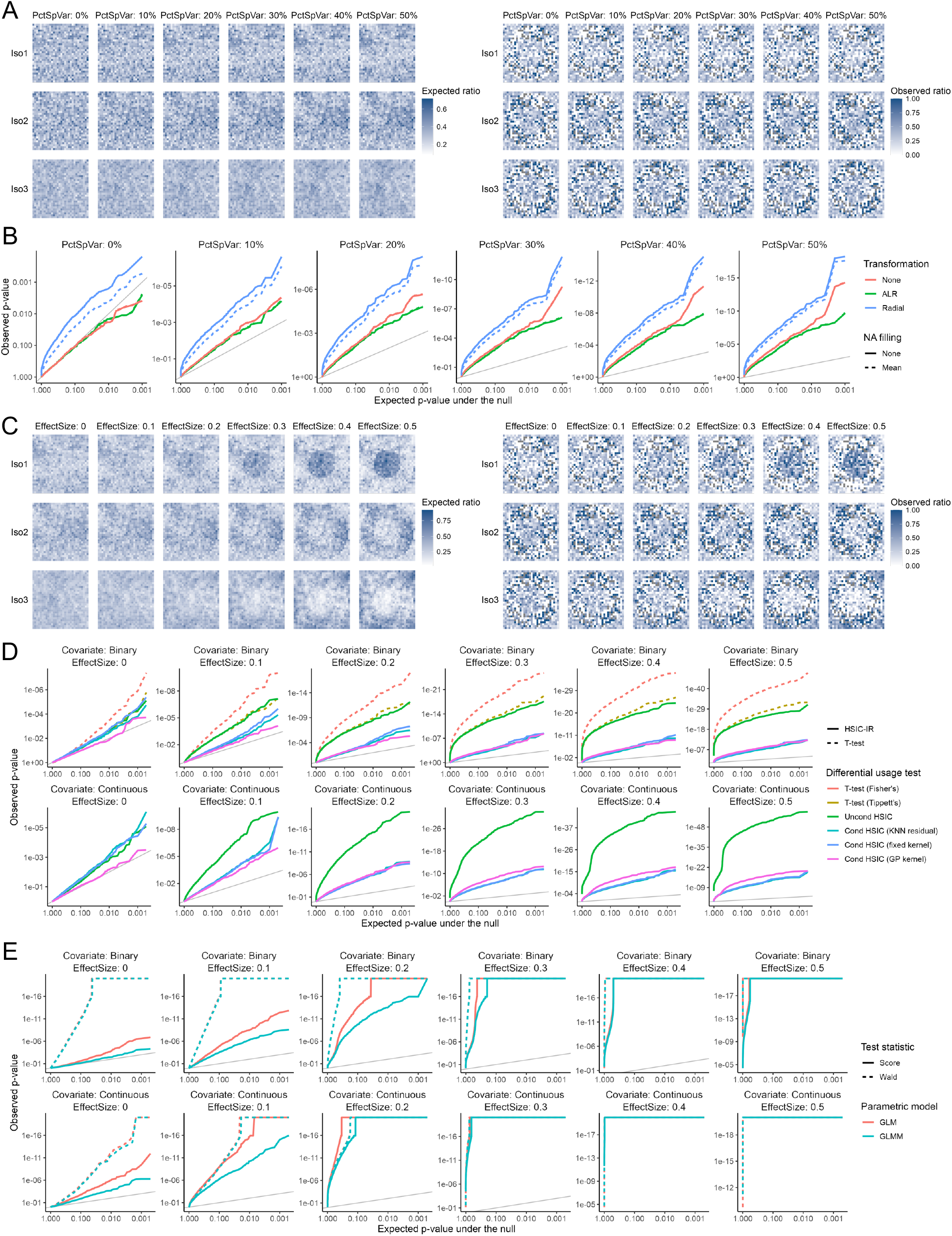
Power analysis for spatial variability and differential usage tests, related to Figure 2. **(A)** Expected and observed isoform ratios in simulations with varying degree of spatial variability, from Scenario 4 (no spatial variance) to Scenario 5 (50% spatial variance) in Figure 2. **(B)** Q-Q plots of HSIC-IR p-values with different ratio transformations and undefined ratio replacement strategies. None means spots with NA ratios are removed and ignored for SV testing. **(C)** Expected and observed isoform ratios in simulations with varying degree of covariate influence, from Scenario 5 (no association) to Scenario 6 (strong association) in Figure 2. **(D)** Q-Q plots of non-parametric DU test p-values. Per-isoform T-test p-values were combined using either Fisher’s or Tippett’s methods. Conditional HSIC with fixed spatial or KNN kernels are included as approximations of the learnable Gaussian process kernel (default). **(E)** Q-Q plots of parametric DU test p-values.

**Figure S2:**
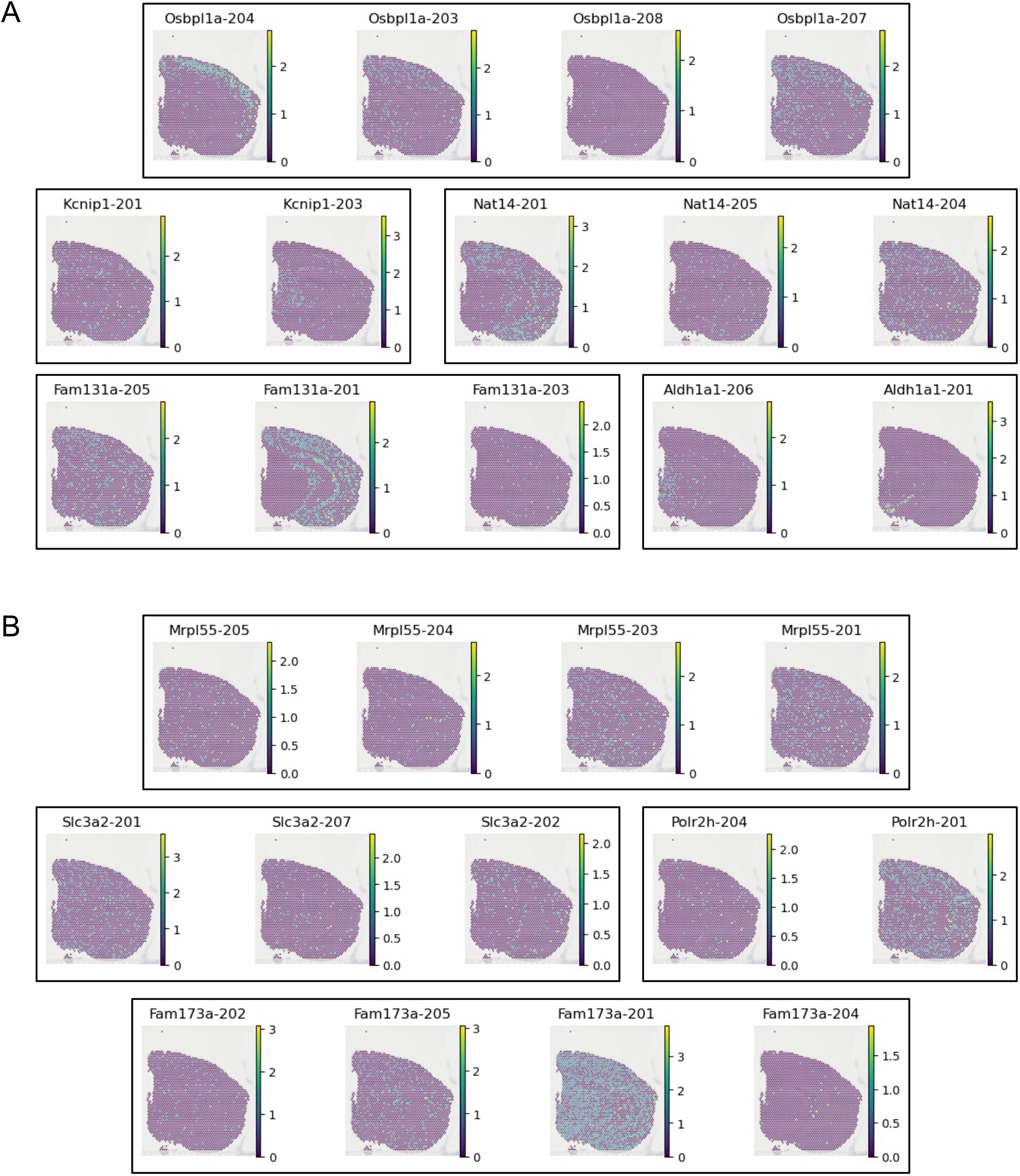
Examples of newly discovered SVP genes (A) and missed original genes (B) in SiT CBS data, related to Figure 3. SiT paper reported 61 genes (shared by replicates CBS1 and CBS2) with ≥2 isoforms called as markers of different brain regions. Our reanalysis identified 150 SVP genes shared between replicates, including 64 additional genes not called significant in neither sample in original study. We note that SiT’s region-marker approach is not technically a spatial variability test but closer to our differential usage test. Accordingly, 6 of the original 61 genes were not called as SVP in neither sample by SPLISOSM. Further validation is needed to confirm the authenticity of these missed regional processing examples, given data sparsity. **(A)** Log-normalized spatial isoform expression of newly discovered genes. **(B)** Log-normalized spatial isoform expression of 6 genes from original 61 not called SVP by SPLISOSM.

**Figure S3.**
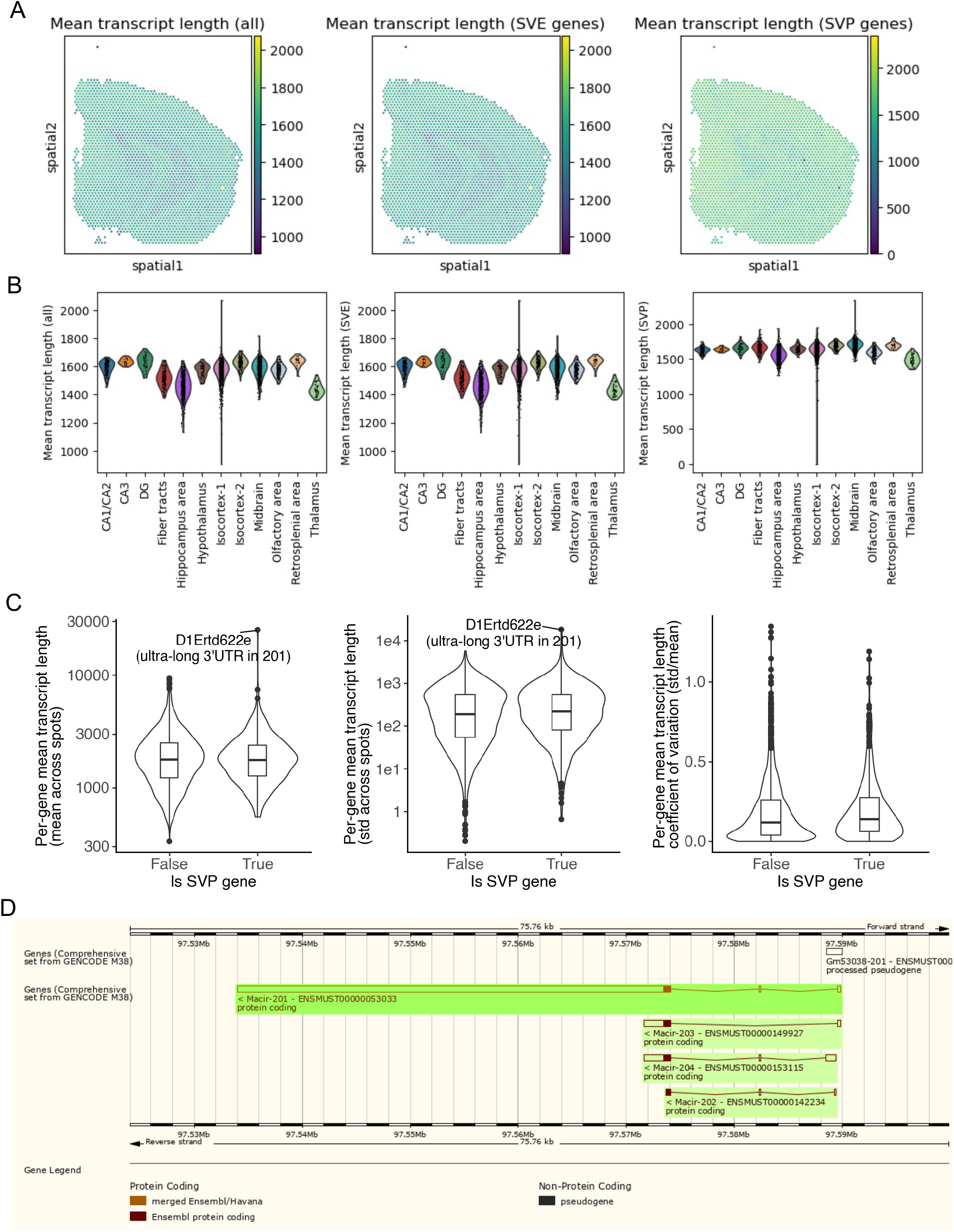
Spatial isoform usage variability is not an artifact from inhomogeneous RNA degradation, related to Figure 3. **(A)** Spatial distribution of per-spot average transcript length in SiT sample CBS2 for all QC-passed transcripts (left), transcripts from spatially variably expressed genes (middle), and spatially variably processed genes (right). **(B)** Per-brain-region distributions of per-spot average transcript length in CBS2 for all QC-passed transcripts (left), transcripts from spatially variably expressed genes (middle), and spatially variably processed genes (right). Each point represents a spatial spot. **(C)** Per-gene average transcript length distribution: mean across spots (left), standard deviation (middle), coefficient of variation (right). Each point represents a gene. SVP gene transcript length distributions not significantly different from background. **(D)** Outlier *D1Ertd622e* (*Macir*) results from collapsing full-splice-junction-match reads to 201 isoform with ultra-long 3’UTR, which itself is not a spatially variable error.

**Figure S4:**
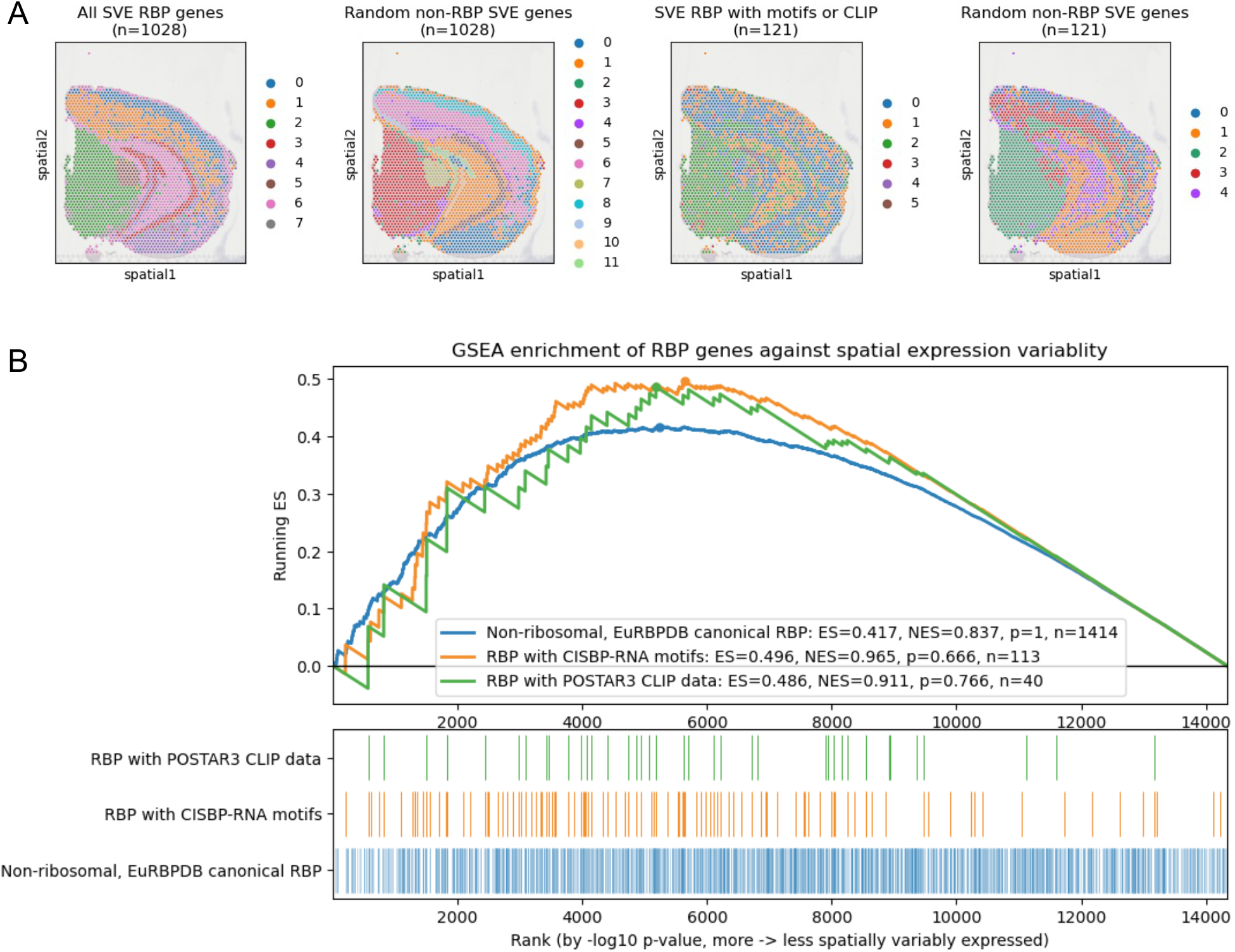
RBP gene expression in adult mouse contains spatial information but is not uniquely spatially specific, related to Figure 3H. **(A)** Short-read gene-expression-based Leiden clustering (resolution=0.9) using different gene sets. **(B)** Gene set enrichment analysis (GSEA) comparing spatial variability of RBP gene expression versus all other genes. Genes ranked by negative log HSIC-GC p-value computed using short-read gene counts. ES = enrichment score, NES = normalized enrichment score.

**Figure S5:**
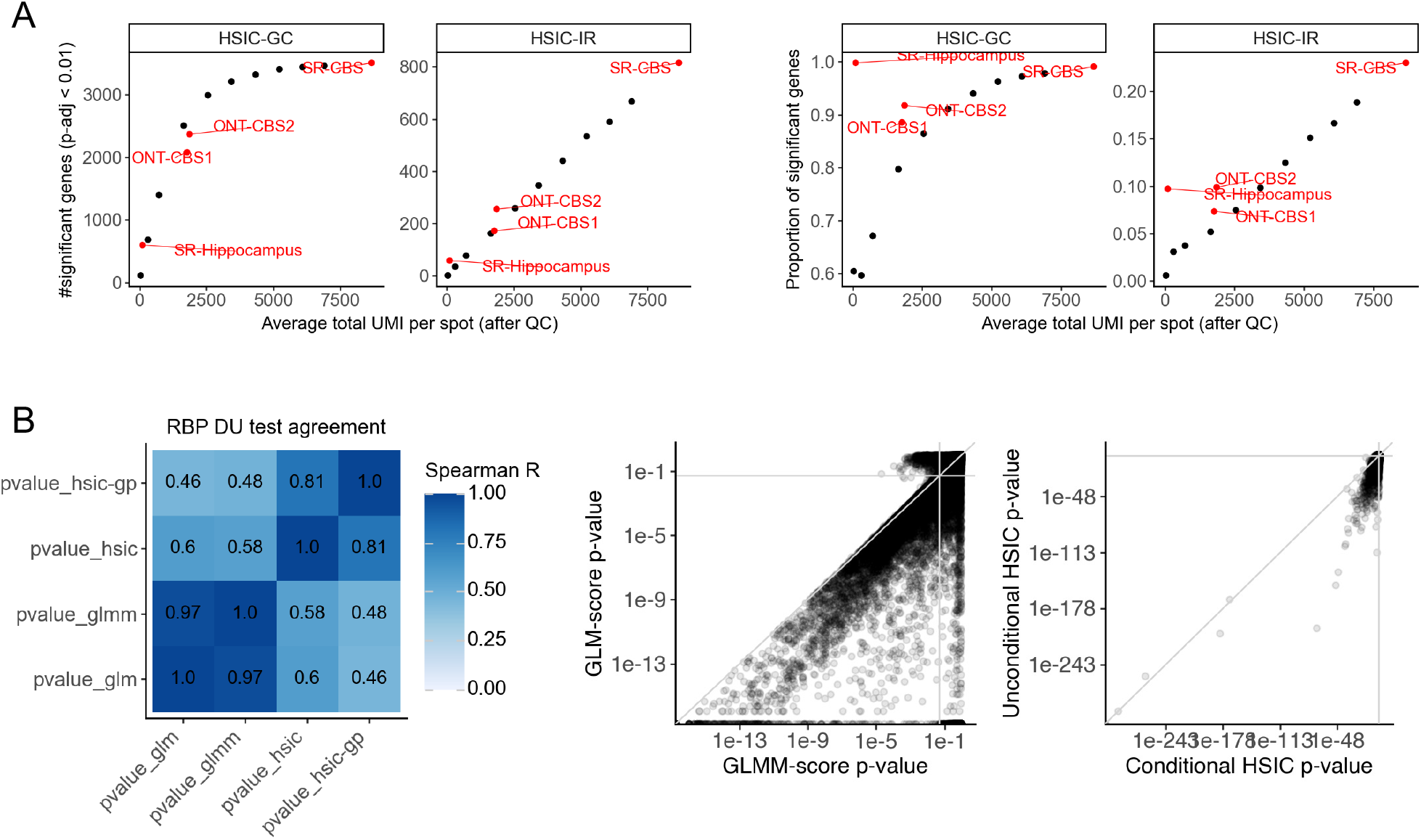
Additional analyses of spatially variable TREND events and their regulation in adult mouse brain, related to Figure 4 and Extended Data Figure 1. **(A)** Number (left) or proportion (right) of spatially variable genes versus sequencing depth in down-sampling experiments. Each black dot represents a short-read Visium CBS sample down-sampled to specific depth. SR-Hippocampus: Slide-seqV2 hippocampus sample with higher spatial resolution but fewer UMI per spot, from Extended Data Figure 1A. **(B)** Pairwise comparison of RBP DU test results. Each dot represents a tested RBP-SVP pair.

**Figure S6:**
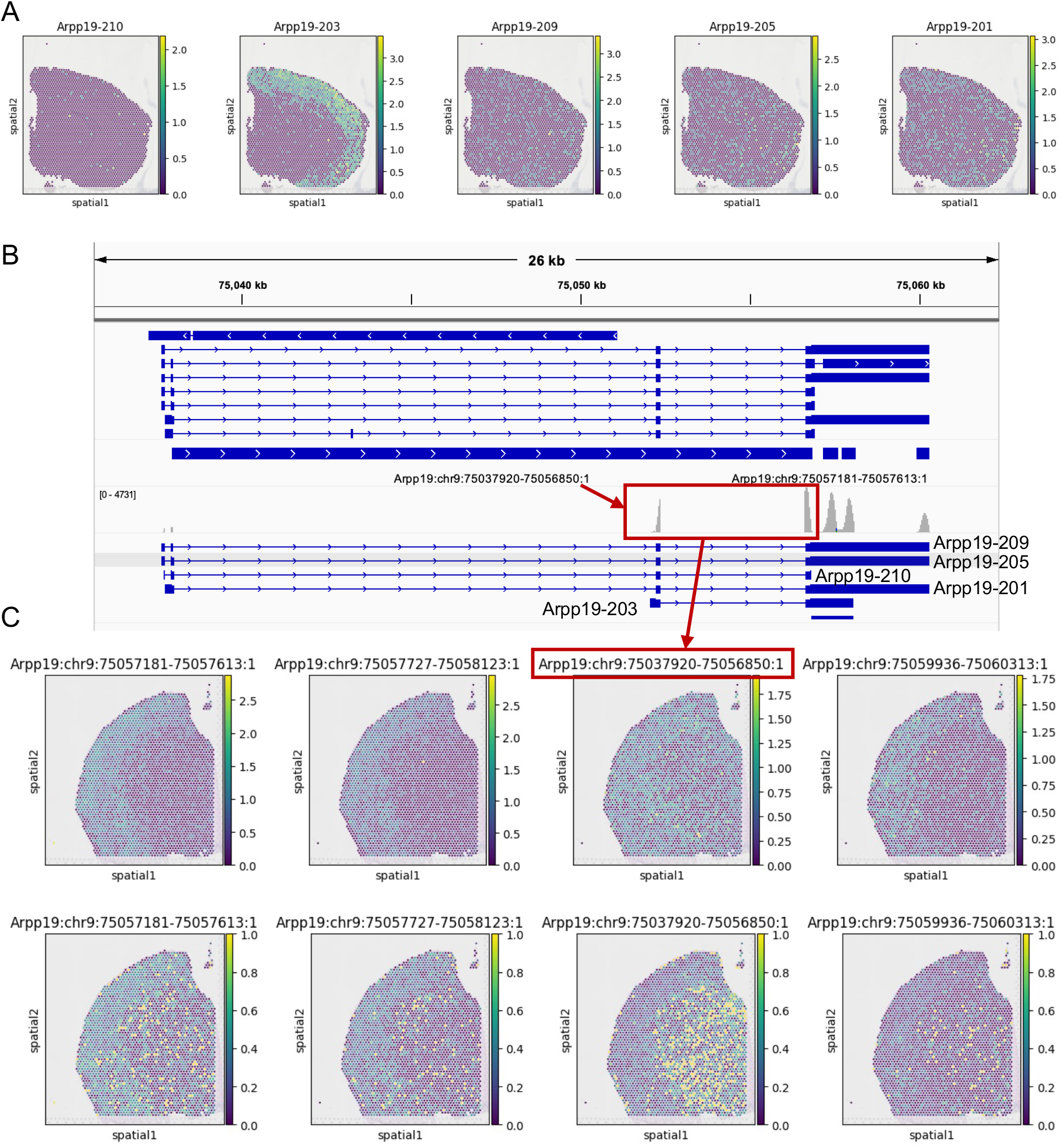
*Arpp19* spatially variable polyadenylation is coupled with its transcript start site selection, related to Extended Data Figure 2C. **(A)** Spatial log-normalized expression of *Arpp19* isoforms in SiT CBS2 sample. ***(B)*** *Arpp19* transcript structure and RNA-seq coverage in 10X Visium CBS sample. **(C)** Spatial distribution of *Arpp19* TREND event log-normalized expression (top) and usage ratios (bottom) in 10X Visium CBS sample. The highlighted upstream poly(A) site is ubiquitously used across brain region, in contrast to cortex-specific usage of downstream sites and Arpp19-203.

**Figure S7:**
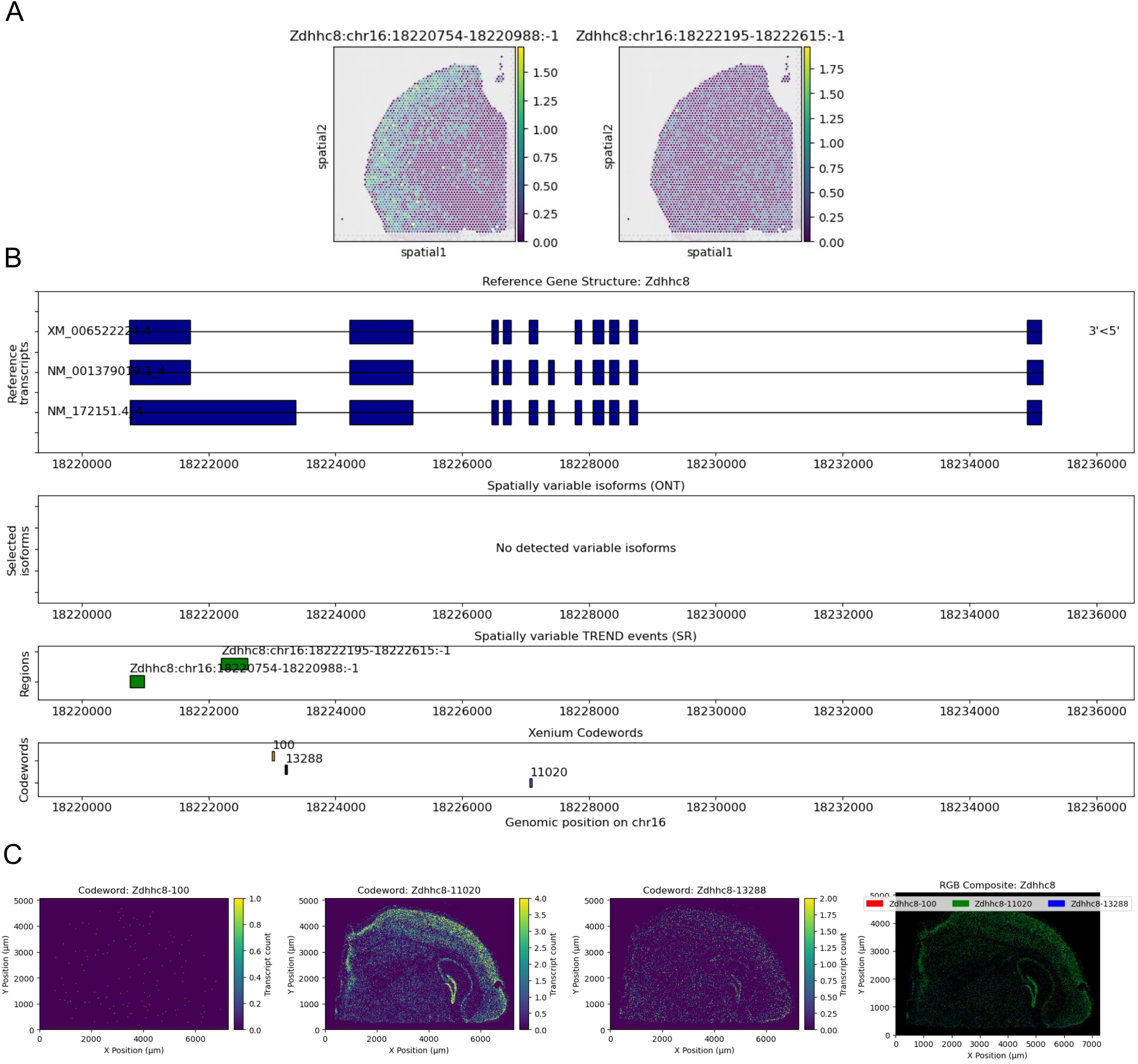
In situ validation of spatially variable last exon usage of *Zdhhc8*, related to Extended Data Figure 2. **(A)** Spatial distribution of log-normalized expression for *Zdhhc8* TREND events in Visium-SR CBS sample. **(B)** Gene structure of Zdhhc8 with annotated RefSeq isoforms and Xenium Prime 5K probe sets. Codewords 100 and 13288 target the longer last exon compatible with upstream TREND event. **(C)** Spatial density of Zdhhc8 codewords in Xenium Prime 5K mouse CBS sample. Codewords 100 and 13288 lack codeword 11020’s spatial specificity to cortical layers, consistent with upstream *Zdhhc8* TREND event expression pattern in (A).

**Figure S8:**
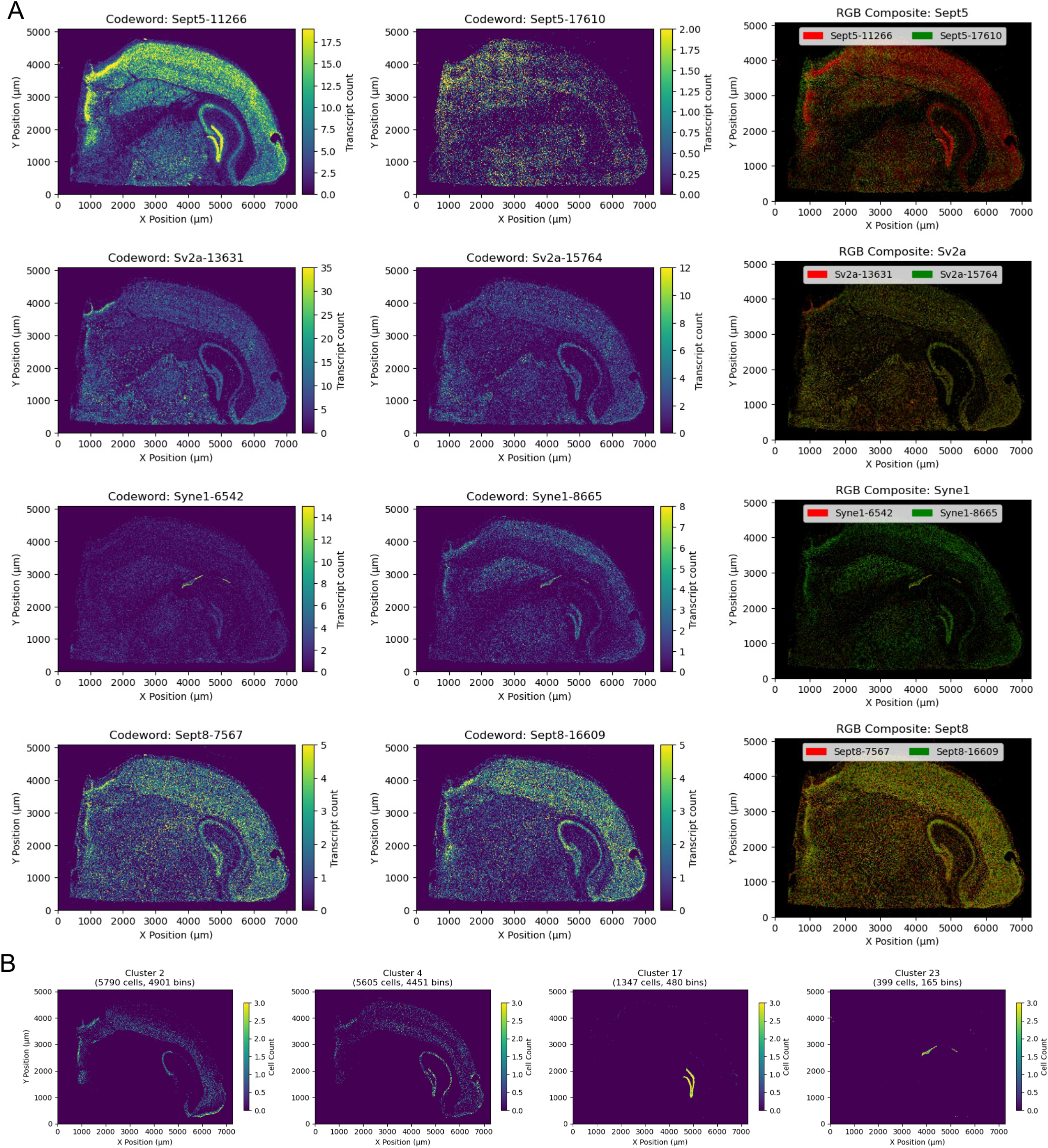
Additional examples of consensus SVP genes in the Xenium Prime 5K CBS dataset, related to Extended Data Figure 2. **(A)** Spatial codeword density of *Septin5, Sv2a, Syne1*, and *Septin8*. **(B)** Spatial distribution of cells from Clusters 2, 4, 17, and 23. All four genes in (A) were called as SVP in either ONT (*Septin5, Septin8*) or SR (*Sv2a, Syne1, Septin8*) CBS datasets, but variable events differ. *Septin8* shows variable last exon usage (Septin8-203/207 vs 204, Figure 5H) in ONT and SR data. However, Xenium Prime 5K panel targets two upstream exons, distinguishing Septin8-205 and 206 from others (203, 204, 207). *Septin8* codewords 7567 and 11609 are differentially used in DG (Cluster 17) and CA2-3 hippocampal regions.

**Figure S9.**
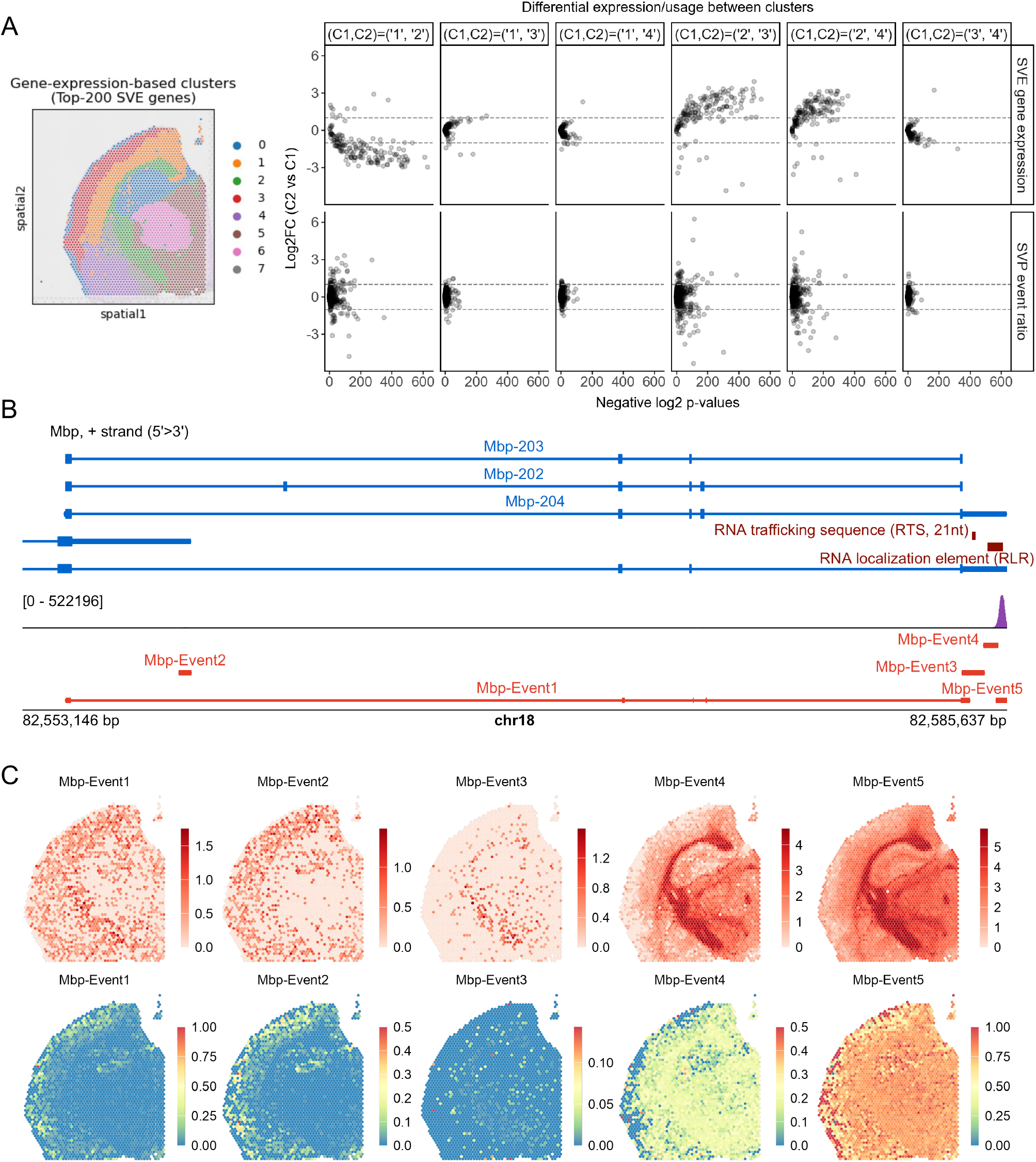
Spatially variable *Mbp* polyadenylation indicates distinct oligodendrocyte subtypes in cortical layers of adult mouse brain, related to Figure 4F. **(A)** Differential gene expression and TREND event usage analyses between expression-based brain regions (left). Clusters 1 and 3: upper (Layers 1-3) and lower (Layers 4-6) isocortical layers; Cluster 2: glia-enriched fiber tract. The uppermost cortical layer 1 is known to be nearly unmyelinated. **(B)** Transcript structure of *Mbp* TREND events with RNA-seq coverage in 10X Visium CBS sample. RNA trafficking sequence (RTS) and RNA localization element (RLR) reside in two wider translational regulatory regions. Events 1 and 2 generate transcripts without RTS and RLR, incapable of transport into myelin sheath. **(C)** Spatial distribution of *Mbp* TREND event log-normalized expression (top) and usage ratios (bottom). Uppermost cortical layers use Events 1, 2 and 5 but not Event 4. Among upstream Events 1-3, Event 2 is mostly absent in fiber tract area.

**Figure S10:**
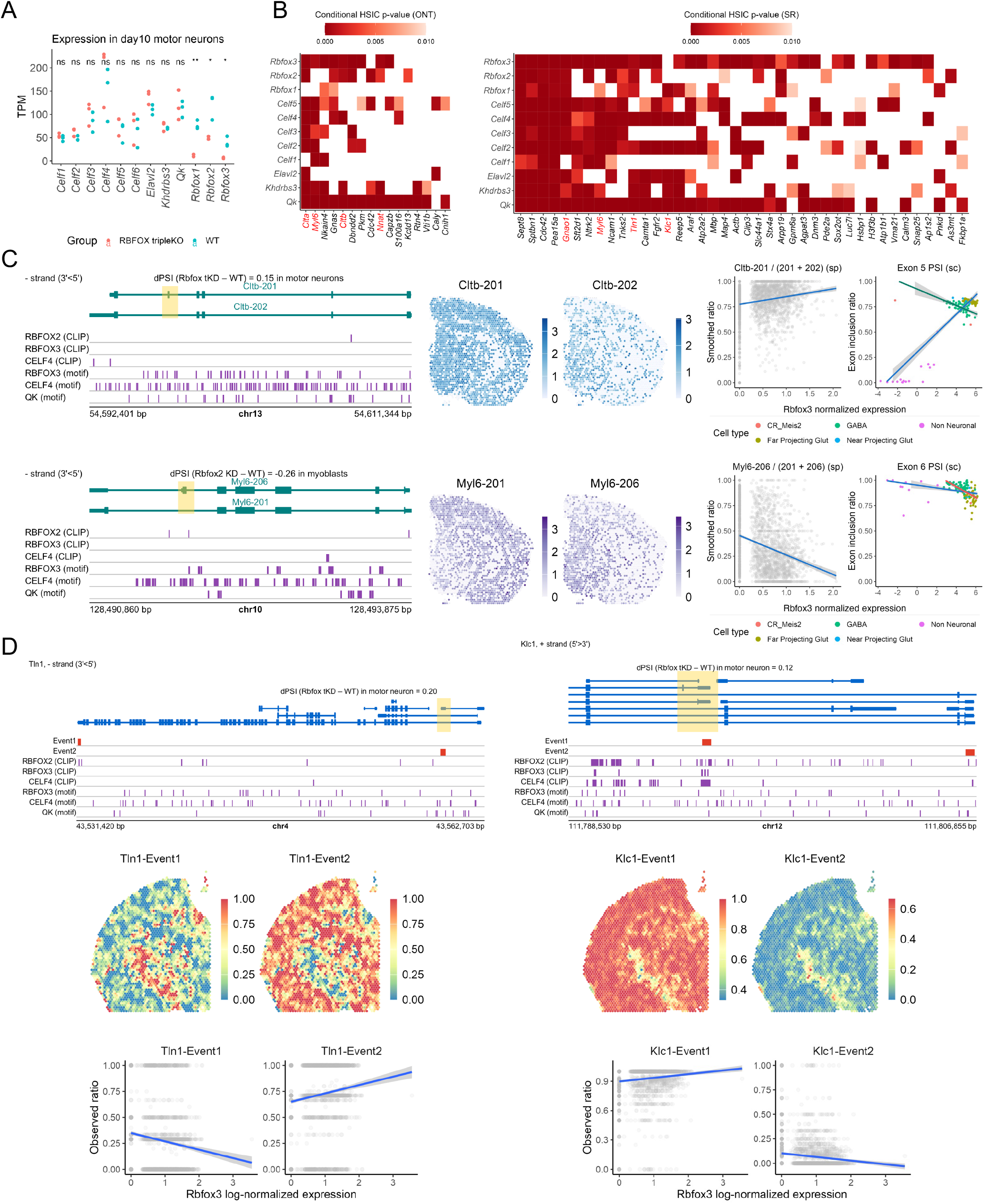
Discrepancy between knockout and observational data indicates cooperativity in RBFOX regulation, related to Extended Data Figure 5. **(A)** Expression changes of selected RBP genes after RBFOX triple knockout where RBFOX genes were disrupted by in-frame stop codons. Group means are compared using T-test. ns: p>0.05; **: p < 0.01. **(B)** Heatmaps displaying DU test p-values between selected RBPs and target SVP genes associated with multiple RBPs. Only significant pairs (p-value < 0.01) are colored. **(C)** Examples of two additional SVP genes whose observed isoform usage association with *Rbfox3* contradicts the triple knockout data. **(D)** Examples of two additional SVP genes whose observed last exon usage association with *Rbfox3* contradicts the triple knockout data.

**Figure S11:**
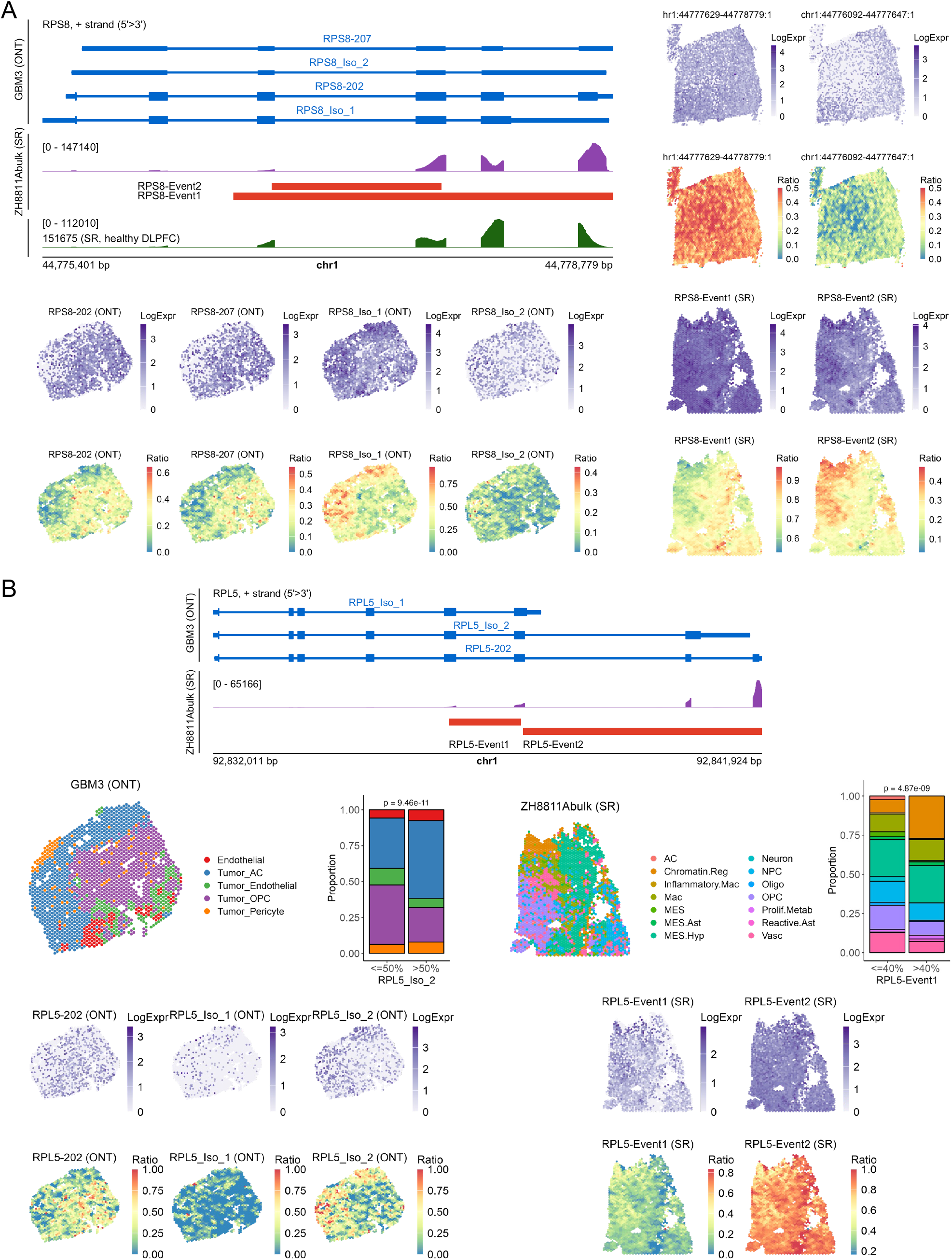
Spatial variable usage of reference and novel ribosomal transcripts in human glioma, related to Figure 6. ***(A)*** *RPS8* transcript structure in the ONT sample GBM3, read coverage in the SR sample ZH8811Abulk and the SR healthy DLPFC sample 151675, and the respective isoform or TREND spatial distribution in each sample. Event counts were locally smoothed for ratio visualization, same in **(B).** ***(B)*** *RPL5* transcript structure in the ONT sample GBM3, read coverage in the SR sample ZH8811Abulk and, and the respective isoform or TREND spatial distribution in each sample. P-value computed using Chi-square test.

## Supplementary Notes on SPLISOSM

## 1 Introduction

Transcript diversity through splicing and alternative 3’end usage is essential for cellular function, yet how this diversity is coordinated across tissues and spatial environments remains poorly understood. SPLISOSM (Spatial Isoform Statistical Modeling) addresses this gap by detecting isoform-level spatial patterns in transcriptomics data with provable guarantees and robust performance on sparse datasets.

This technical note details SPLISOSM’s two core capabilities: (1) identifying genes with significant spatial variation in isoform usage (spatial variability test), and (2) discovering regulatory factors driving these patterns while accounting for spatial confounding (differential association test). Our methods balance statistical rigor with computational efficiency to handle datasets with tens of thousands of spatial locations.

We present our approach in two complementary frameworks. Section 2 describes our non-parametric kernel-based methods, introducing the Hilbert-Schmidt Independence Criterion (HSIC) and two key theoretical innovations. First, we prove that all low-rank kernel approximations inevitably sacrifice statistical power (**Theorem 1**), which led us to develop improved full-rank and low-rank kernel designs that maximize detection sensitivity. Second, we establish a mathematically sound approach to handle undefined ratios in compositional data (**Theorem 2**), addressing a critical challenge in analyzing sparse spatial transcriptomics data. Our framework also introduces a novel conditional testing approach that effectively controls for spatial confounding effects. Section 3 outlines parametric alternatives using Generalized Linear Mixed Models (GLMM), including some of our unsuccessful attempts and the working example of GLMM differential usage test based on the score statistic.

Through these specialized statistical innovations, SPLISOSM enables researchers to investigate how transcript diversity contributes to cellular specialization and tissue organization across development, homeostasis, and disease states.

## 2 Non-parametric spatial pattern discovery using kernel independence test

### 2.1 Hilbert-Schmidt independence criterion (HSIC)

The Hilbert-Schmidt Independence Criterion (HSIC) provides a powerful framework for testing statistical independence between random variables[GBSS05]. Unlike methods that rely on linear correlations, HSIC can detect complex, non-linear relationships through kernel functions. In this section, we introduce HSIC and discuss its application to spatial pattern discovery.

#### 2.1.1 From correlation to dependence testing

For finite-dimensional random variables *X* ∈ ℝ^*m*^ and *Y* ∈ ℝ^*p*^, a common approach to assess dependence is through the cross-covariance matrix *C*_*XY*_ = *C*_*ij*_ = Cov(*X*_*i*_, *Y*_*j*_). The squared Frobenius norm of this matrix (the unnormalized RV coefficient[RE76])

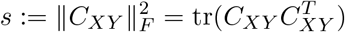

equals zero if and only if *X* and *Y* are uncorrelated across all dimensions.

However, for non-Gaussian distributions, uncorrelatedness is weaker than independence. HSIC addresses this limitation by mapping variables into reproducing kernel Hilbert spaces (RKHSs), allowing it to detect more general forms of dependence.

#### 2.1.2 Mathematical definition and empirical computation

Let 𝒳 and 𝒴 be separable spaces with probability distributions *P*_*X*_ and *P*_*Y*_. Given positive definite kernels *k* : 𝒳 × 𝒳 → ℝ and *l* : 𝒴 × 𝒴 → ℝ with corresponding RKHSs ℱ and 𝒢, we can implicitly map data points into (infinite-dimensional) feature spaces using the canonical feature map *ϕ* : 𝒳 → ℱ defined by *ϕ*(*X*) = *k*(*x*, ·). Accordingly, the cross-covariance operator *C*_*XY*_ : 𝒢 → ℱ generalizes the finite-dimensional cross-covariance matrix to these infinite-dimensional spaces. HSIC is defined as the squared Hilbert-Schmidt norm of this operator

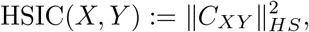

which can be equivalently expressed as

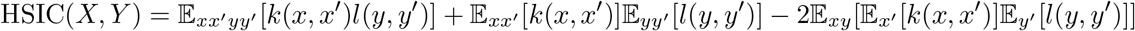

where expectations are taken over independent pairs drawn from the joint distribution.

Given observed data pairs 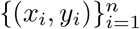, the empirical HSIC estimator is

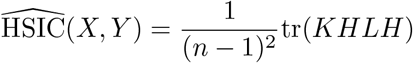

where *K*_*ij*_ = *k*(*x*_*i*_, *x*_*j*_), *L*_*ij*_ = *l*(*y*_*i*_, *y*_*j*_), and 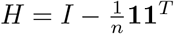 is the centering matrix.

A key theoretical result states that, with universal kernels *k* and *l*, HSIC(*X, Y*) = 0 if and only if *X* and *Y* are statistically independent (Theorem 4 of [GBSS05]). Moreover, we can define a test statistic based on the empirical HSIC estimator, which follows an asymptotic distribution of *χ*^2^ mixture under the null hypothesis of independent *X* and *Y* (Theorem 4 of [ZPJS12]),

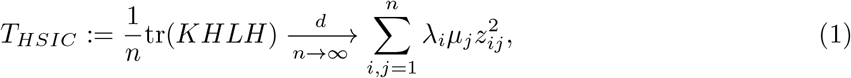

where *λ*_*i*_ and *µ*_*j*_ are eigenvalues of centered kernels *HKH* and *HLH*, and *Z*_*ij*_ are i.i.d. standard Gaussian variables.

#### 2.1.3 Application to spatial pattern detection

The analytical form of the asymptotic null distribution in Equation (1) enables a calibrated, permutation-free test for spatial pattern discovery in transcriptomics data. In this context,

- *X* represents the biological variable of interest (e.g. gene expression, isoform usage)
- *Y* represents spatial coordinates of cells/spots
- *K* is the variable kernel matrix encoding cross-feature relationships
- *L* is the spatial kernel matrix encoding spatial relationships

SPARK-X[ZSZ21] first applied HSIC to detect spatial variability in gene expression *X* using a custom spatial kernel *L* for coordinate *Y* ∈ R^2^ and a linear projection kernel for gene expression *X* ∈ ℝ. While computationally efficient and statistically powerful on sparse data, it has two major limitations:

1. It cannot handle multivariate spatial variables (e.g. isoform usage *X* ∈ [0, 1]^*q*^ of a given gene) and non-Euclidean data (e.g. region annotation *X* ∈ {*c*_1_, …, *c*_*k*_} of with *k* categories).
2. Its low-rank spatial kernel design leads to significant power loss.

In SPLISOSM, we address both limitations by implementing a multivariate spatial variability test for isoform expression and relative usage with theory-guided spatial kernel design.

For notational simplicity, we use *I*(*X*; *K*_*sp*_) to denote the general spatial variability testing procedure with predefined spatial kernel *K*_*sp*_, where the test statistic 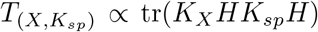 is evaluated against a specific null distribution (e.g. as in Equation (1)). We also do not distinguish between a random variable *X* ∈ 𝒳’s n-sample realization *X* = [*x*_1_, …, *x*_*n*_]^*T*^ ∈ 𝒳 ^*n*^, and a random field (set of *n* 𝒳-valued spatial random variables) *X* = {***x***_***i***_ ∈ 𝒳} ∈ 𝒳 ^*n*^ indexed by spatial location. This is because given the observation *X* = [*x*_1_, …, *x*_*n*_]^*T*^, we can always construct a random field where {***x***_***i***_ = *x*_*i*_} are point masses that deterministically vary with location. Depending on the context, a spatial pattern can refer to both the realization, or the distribution of a spatial random field.

### 2.2 Low-rank approximation of spatial kernel

While HSIC provides a powerful framework for detecting spatial patterns, its practical application faces computational challenges. The memory and computation requirements grow quadratically with sample size *n*, with kernel eigendecomposition for computing the null distribution in Equation (1) having computational complexity of *O*(*n*^3^). This motivates the use of low-rank approximations to improve scalability. Several approaches have been proposed to address this challenge[ZFGS18], including block-wise testing (splitting *n* data points into *b* blocks of size *n/b < n* to compute block-wise statistics) and low-rank kernel approximations (e.g., Nyström’s method with *m* inducing points). However, to the best of our knowledge, the theoretical implications on test power of these approximations have not been thoroughly analyzed.

In the following sections, we address this gap by demonstrating the fundamental limitations of low-rank kernel approximations in the context of spatial pattern detection (**Theorem 1, Section 2.2.2**). *Our theoretical results recommend using full-rank spatial kernels when possible, which is typically manageable for current spatial transcriptomics datasets (n* ≤ 10000)^1^. Nevertheless, we recognize the need for more efficient methods as datasets continue to grow in size and resolution. In response, we propose an alternative low-rank approximation approach based on Graph Fourier Transform (GFT) that balances computational efficiency and statistical power (**Section 2.2.3**).

#### 2.2.1 Preliminaries on kernel eigenvalues

We first introduce the following useful results on how transformations affect kernel eigenvalues.

##### Lemma 1 (Poincaré separation theorem or Cauchy interlacing theorem).

*Let A be an n* × *n symmetric matrix with eigenvalues λ*_1_ ≥ *λ*_2_ ≥ … ≥ *λ*_*n*_. *For any n* × *m matrix P with orthonormal columns (P*^*T*^ *P* = *I*_*m*_*) where m < n, let B* = *P*^*T*^ *AP with eigenvalues µ*_1_ ≥ *µ*_2_ ≥ … ≥ *µ*_*m*_. *Then the following interlacing property holds:*

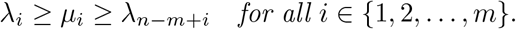

*Proof*. The Poincaré separation theorem follows directly from the Courant-Fischer min-max theorem. Note ⟨*Bx, x*⟩ = ⟨*P*^*T*^ *APx, x*⟩ = ⟨*A*(*Px*), *Px*⟩. The eigenvalues of *A* and *B* are thus linked and interlaced through their respective Rayleigh quotients.

□

Using the separation theorem, we can derive the eigenvalue interlacing properties that will be crucial for our analysis of low-rank (Section 2.2.2) and missing-data kernel tests (Section 2.3).

##### Corollary 1.1 (Eigenvalues of the centered kernel).

*Let K be an n* × *n kernel matrix, and* 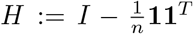 *the centering matrix. Denote λ*_1_ ≥ *λ*_2_ ≥ …≥ *λ*_*n*_ *and µ*_1_ ≥ *µ*_2_ ≥ … ≥ *µ*_*n*_ *the eigenvalues of K and HKH, respectively. Then we have the eigenvalue interlacing*

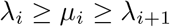

*for i* ∈ {1, …, *n* − 1}. *Specifically, λ*_*n*−1_ ≥ *µ*_*n*−1_ ≥ *λ*_*n*_ ≥ *µ*_*n*_ = 0.

*Proof*. The centering matrix *H* has rank *n*−1 with eigenvalues 1 (multiplicity *n*−1) and 0 (multiplicity 1). Thus, *H* = *UU*^*T*^ where *U* is an *n* × (*n* − 1) semi-orthogonal matrix with *U*^*T*^ *U* = *I*_*n*−1_. This gives *HKH* = *U* (*U*^*T*^ *KU*)*U*^*T*^. The matrices *B* := *U*^*T*^ *KU* and *HKH* = *UBU*^*T*^ share the same non-zero eigenvalues, as for any eigenpair (*λ, ν*) of *HKH*, we have *B*(*U*^*T*^ *ν*) = *U*^*T*^ (*UBU*^*T*^)*ν* = *λ*(*U*^*T*^ *ν*). Applying Lemma 1 with *P* = *U*^*T*^, we obtain the interlacing property. Furthermore, since *H***1** = **0**, we have *HKH***1** = **0**, confirming that *µ*_*n*_ = 0.

##### Corollary 1.2 (Eigenvalues of the principal submatrix).

*Let K be an n* × *n kernel matrix with eigenvalues λ*_1_ ≥ *λ*_2_ ≥ …≥ *λ*_*n*_. *For any principal submatrix K*_*m*_ *of K with size m* × *m (assuming K*_*m*_ *contains the first m rows and columns of K, m < n) and eigenvalues µ*_1_ ≥ *µ*_2_ ≥ … ≥ *µ*_*m*_, *the following interlacing property holds:*

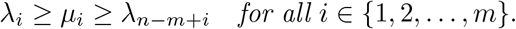

*Proof*. Let *P* be the *n* × *m* matrix formed by taking the first *m* columns of the *n* × *n* identity matrix. Then *P*^*T*^ *P* = *I*_*m*_ and *K*_*m*_ = *P*^*T*^ *KP*. The result follows directly from applying Lemma 1. □

##### Proposition 2.1.

*The principal submatrix of a kernel matrix is also a kernel (positive semidefinite)*.

#### 2.2.2 Low-rank kernel independence test has limited power

We now present our first major result on the fundamental limitation of independence tests with low-rank kernels. Our analysis reveals that certain spatial patterns remain undetectable when using low-rank spatial kernels, regardless of the testing procedure. This has important implications for existing methods like SPARK-X[ZSZ21], which we will discuss in detail.

##### Theorem 1 (Limited power of low-rank kernel tests).

*Consider a one-dimensional target variable X* ∈ ℝ^*n*^ *observed at n locations. Let K*_*sp*_ *be an n* × *n spatial kernel matrix of rank d < n* − 1, *and let K*_*X*_ := *XX*^*T*^ *be the linear target kernel matrix used in the spatial kernel independence test I*(*X*; *K*_*sp*_). *Then there exists a non-constant spatial pattern X*_0_ *for which I*(*X*_0_; *K*_*sp*_) *has no detection power. Moreover, such undetectable patterns form a subspace of dimension at least n* − *d*.

*Proof*. The unscaled test statistic with *K*_*sp*_ and a linear kernel *K*_*X*_ can be expressed as

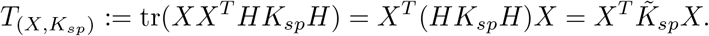

Using the eigendecomposition of 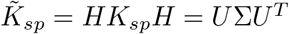, we get

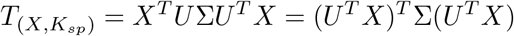

where *U* is an *n* × *n* orthogonal matrix and Σ = diag{*λ*_1_ ≥ *λ*_2_ ≥ … ≥ *λ*_*n*_}. From Corollary 1.1, we know that Σ has at least *n* − *d* ≥ 2 zero entries *λ*_*d*+1_ = *λ*_*d*+2_ = … = *λ*_*n*_ = 0, since the rank of *K*_*sp*_ is *d*. For *T*_(*X,K*)_ = 0, we need to construct *X* such that *U*^*T*^ *X* only has non-zero elements in positions corresponding to zero eigenvalues in Σ. This can be achieved by defining *X*_0_ = *UY* where *Y* = [**0**_*d*_, *y*_*d*+1_, …, *y*_*n*_]^*T*^ ∈ ℝ^*n*^ and **0**_*d*_ is a zero vector of length *d*. Moreover, *X*_0_ can be chosen to be non-constant as long as *Y* is not proportional to *U*^*T*^ **1** ∈ ℝ^*n*^, which is always possible since it has more than one degree of freedom.

Let *U*_*trunc*_ be the *n* × (*n* − *d*) sub-matrix of *U* containing the last *n* − *d* eigenvectors of *HK*_*sp*_*H*:

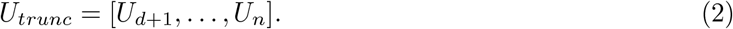

Following our construction, we see that 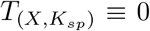 for any *X* in the image space of *U*_*trunc*_. Since *U* has orthonormal columns, we have dim(Img(*U*_*trunc*_)) = dim(Col(*U*_*trunc*_)) = *n* − *d*, which completes the proof. □

The result in Theorem 1 naturally extends to multi-dimensional data with non-linear kernels. To see that, we note that any *n* × *n* target kernel matrix *K*_*X*_ can be expressed as an inner product *K*_*X*_ = ⟨*ϕ*(*X*), *ϕ*(*X*)⟩ = ΨΨ^*T*^ where Ψ ∈ ℱ := span{*ϕ*(*X*)} represents the transformed data matrix in feature space. The corresponding unscaled test statistic becomes

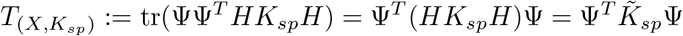

which recovers the linear kernel case in Theorem 1 when Ψ ≡ *X*. Following the same logic as before, we can construct a feature-space pattern Ψ_0_ ∈ ℱ that yield zero test statistic (e.g. Ψ_0_ ∈ Img(*U*_*trunc*_)). However, to obtain the corresponding spatial pattern *X*_0_, we need to map Ψ_0_ from feature space ℱ back to data space 𝒳, which is the classical pre-image problem and may not always have an exact solution (See [SS01] Chapter 18 for details). In the general case, the existence of undetectable spatial pattern depends on the specific choices of *K*_*sp*_ and the target kernel *K*_*X*_.

Under Theorem 1, the space of undetectable patterns grows larger as the rank of the spatial kernel decreases. The following corollary further characterizes this limitation.

##### Corollary 1.3 (Orthogonal decomposition of spatial patterns).

*Any spatial pattern X* ∈ ℝ^*n*^ *can be decomposed into two orthogonal components*

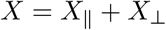

*such that* 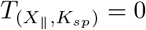 *with linear target kernel, X*_⊥_ ∈ *V* ⊂ ℝ^*n*^, *and* dim(*V*) ≤ *d* = *rank*(*K*_*sp*_).

*Proof*. We choose *X*_‖_ to be the projection of *X* onto Img(*U*_*trunc*_) and *X*_⊥_ to be the residual, which lies in Img(*U*_*trunc*_)^⊥^. This residual space is also the image space of the sub-matrix containing the first *d* columns of *U*.

Specifically, we have

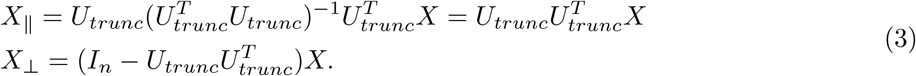

□

In essence, Equations (2) and (3) provide us with a kernel-specific projection 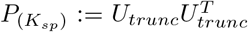 that transforms an arbitrary spatial pattern *X* to the no-power space of the variability test with *K*_*sp*_. The information loss under this projection is determined by the rank of *K*_*sp*_.

To illustrate these theoretical findings, we examine SPARK-X, a widely used method for detecting spatially variable gene expression that employs a low-rank spatial kernel.

##### Definition 2.1 (SPARK-X kernel[ZSZ21]).

*Let S be the n* × 2 *(or* 3*) centered and standardized 2D (or 3D) spatial coordinates. The SPARK-X spatial kernel is defined as:*

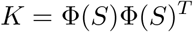

*where* F = {*ϕ*_0_, …, *ϕ*_10_} *is a set of 11 predefined transformation functions ϕ* : ℝ^2^ → ℝ^2^, *and* Φ(*S*) *is of dimension n* × 22. *Specifically:*

- *The first transformation is the projection ϕ*_0_(*S*) = *S*(*S*^*T*^ *S*)^−1*/*2^
- *The next 5 are per-dimension Gaussian transformations with varying bandwidths w*_*i*_

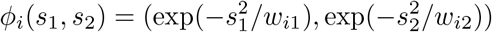
- *The final 5 are per-dimension cosine transformations with varying frequencies γ*_*i*_

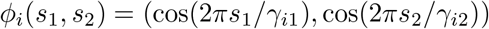

##### Definition 2.2 (SPARK-X spatially variable expression test[ZSZ21]).

*Let K be the n* × *n SPARK-X spatial kernel, and X* ∈ ℝ^*n*^ *the centered and standardized expression of a given gene. We define the joint SPARK-X test statistic as*

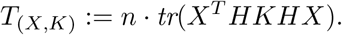

*Under the null hypothesis that spatial coordinates and gene expression are independent, T*_(*X,K*)_ *asymptotically follows a mixture of χ*^2^ *distributions. The original SPARK-X paper tests the 11 spatial transformations in K independently and then combines them into a single p-value. Since T*_(*X,K*)_ *is the sum of 11 non-negative statistics, T*_(*X,K*)_ = 0 *implies that all individual test statistics are zero*.

In typical spatial transcriptomics datasets, we have *n* ≫ 22 ≥ rank(*K*). Applying Theorem 1, we can conclude that the space of spatial patterns where SPARK-X has no statistical power is much larger than the space where it can detect patterns. By performing eigendecomposition of the low-rank matrix *HKH* = *H*F(*S*)Φ(*S*)^*T*^ *H*, we can transform arbitrary spatial patterns to become “invisible” to SPARK-X with minimal information loss, as illustrated in Figure 1.

**Figure 1.**
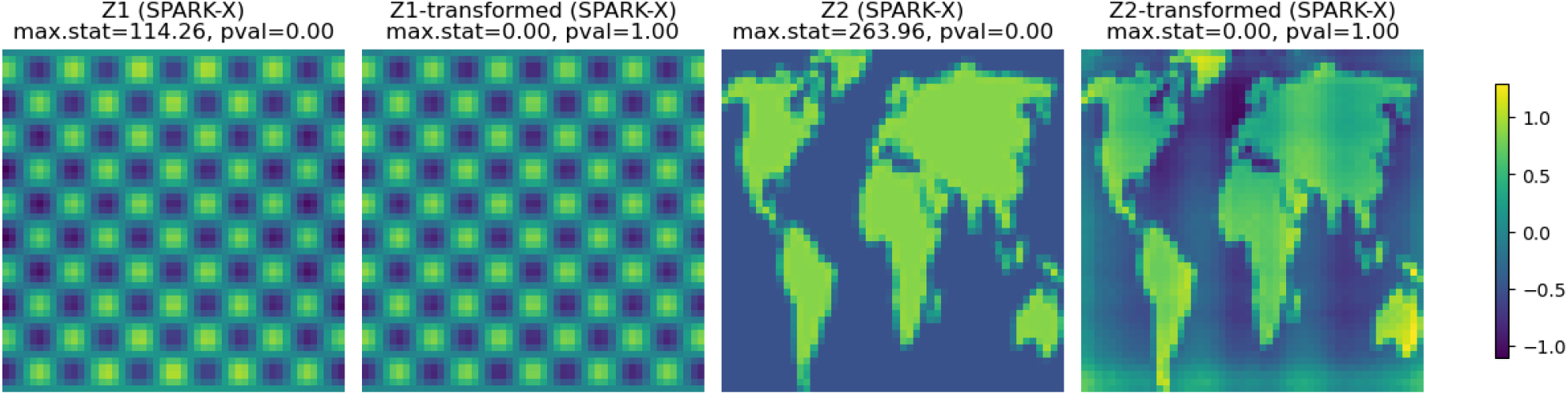
Transformation of arbitrary spatial patterns to make them undetectable by SPARK-X.

#### 2.2.3 Kernel approximation using Graph Fourier Transform

The general trade-off between computational efficiency and statistical power is not limited to low-rank approximations. A common approach to scaling up *I*(*X*; *K*_*sp*_) is to inject sparsity through thresholding

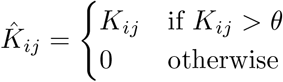

where *θ* determines the spatial range of dependencies. The resulting banded matrix kernel 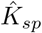 is typically full-rank and sparse, reducing both space complexity (to sub-quadratic) and computational complexity of eigendecomposition (to at least quadratic)^2^. However, this efficiency gain comes at the cost of statistical power 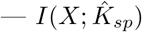 cannot detect global patterns that extend beyond the context defined by *θ*.

Our analysis raises an important question: *if some power loss is inevitable with approximation, can we design spatial kernels where the undetectable pattern space contains the least biologically interesting patterns?* We propose a principled approach based on Graph Fourier Transform (GFT), which decomposes spatial patterns into frequency components and prioritizes the most informative frequencies rather than relying on preselected transformations. This preserves both local and global pattern detection capability while maintaining computational efficiency.

We begin by defining a full-rank spatial kernel with favorable properties.

##### Definition 2.3 (Smoother ICAR kernel[SRF

^+^**23])**. *Let S be the n* × 2 *(or* 3*) spatial coordinates of spots/cells. Let W be the n* × *n adjacency weight matrix of the spatial graph built from S, where W*_*ij*_ = 1 *if i and j are mutual k-nearest neighbors (otherwise 0), and W*_*ii*_ = 0^3^. *The Smoother ICAR (intrinsic conditional autoregressive) kernel is defined as:*

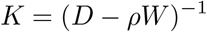

*where D* = *diag*{∑_*j*_ *W*_*ij*_} *is the degree matrix and ρ* ∈ [0, 1) *is the spatial autocorrelation coefficient ensuring invertibility*.

This kernel offers several advantages:

- *K* is positive definite and defines a multivariate Gaussian distribution on the spatial random field that can be factorized into the product of local conditional distributions
- *K* captures topological properties and is invariant to scaling, translation, and rotation of *S*
- *K*^−1^ is sparse with at most (*k* + 1)*n* non-zero entries, enabling efficient eigendecomposition
- The eigenvectors of *K* represent meaningful spatial frequency components (Proposition 2.2)

##### Proposition 2.2 (Connection to spectrum graph theory[Chu97]).

*Let L* = *D* − *W be the graph Laplacian of spatial graph G, and K* = (*D* − *ρW*)^−1^ *the ICAR kernel. Let* 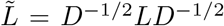 *and* 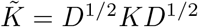 *denote the normalized Laplacian and normalized spatial kernel, respectively*. 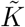 *and* 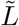 *share the same eigenvectors, which form the basis for the Graph Fourier Transform*.

*The eigenvalues λ*_*i*_ *of* 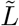 *(called the spectrum of G) are conventionally ordered as* 0 = *λ*_1_ ≤ *λ*_2_ ≤ … ≤ *λ*_*n*_ ≤ 2, *and the corresponding eigenvectors (called eigenfunctions) represent spatial patterns with increasing spatial frequency. In particular, the first eigenvector associated with λ*_1_ = 0 *is the square root of node degree u*_1_ = *D*^1*/*2^**1**. *Moreover, for ρ* ∈ [0, 1), *if* 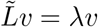, *then* 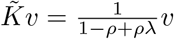.

*Proof*. We can rewrite the inverse of the normalized ICAR kernel as 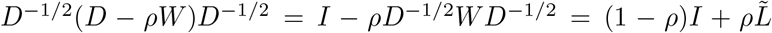. For any eigenvector *v* of 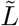 with eigenvalue *λ*, we have 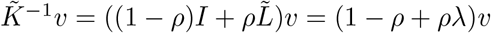, implying 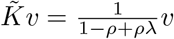.

□

The Rayleigh quotient of the normalized graph Laplacian for a spatial pattern *X* ∈ ℝ^*n*^

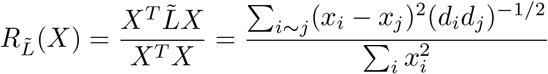

quantifies the smoothness of *X*. Higher values indicate rapid changes between adjacent nodes (higher frequency), while lower values indicate smoother variation (lower frequency).

By the min-max theorem, the eigenvectors of 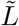 (and thus of 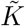) correspond to spatial patterns of increasing frequency. For a regular graph (i.e. every node has the same number of neighbors), the first eigenvector is constant with zero frequency, while each subsequent eigenvector minimizes the Rayleigh quotient subject to being orthogonal to all previous eigenvectors, creating a natural ordering of spatial frequencies (eigenvalues). Given the *n* eigenvectors *U* = [*u*_1_, …, *u*_*n*_], we can decompose any spatial pattern *X* ∈ ℝ^*n*^ through Graph Fourier Transform

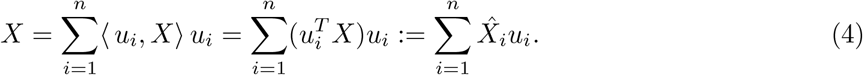

##### Definition 2.4 (Low-rank GFT kernel).

*Let* 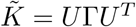 *be the eigendecomposition of the normalized ICAR kernel, where* Γ = *diag*(*γ*_1_, …, *γ*_*n*_) *with γ*_1_ ≥ *γ*_2_ ≥ … ≥ *γ*_*n*_. *The rank-r GFT kernel is defined as*

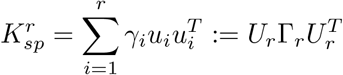

*where u*_*i*_ *is the i-th column of U representing the i-th lowest-frequency component of the spatial graph*.

From Equation (4), we see that the spatial variability test statistic 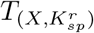 with linear target kernel is the sum of squares of Fourier coefficients weighted by inverse frequency *γ* under a low-pass filter *γ*_*i*_ = 0 for *i* ∈ (*r, n*]

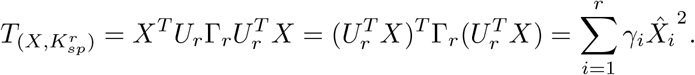

By scaling *γ* (e.g. exponential normalization[CLJ^+^24]), we can encourage the spatial kernel 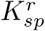 to prioritize specific frequencies and patterns. Note 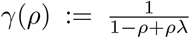 itself is parametrized by the autocorrelation coefficient *ρ*. For regular graphs, centering *X* removes the first coefficient 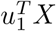 in Equation (4). Moreover, the no-power space 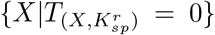 is spanned by the zero-frequency component and the *n* − *r* high-frequency eigenfunctions. Figure 2 visualizes the effect of kernel rank *r* on test power. As we increase the rank of the GFT-based approximation, the no-power transformation space primarily contains high-frequency patterns that are often less biologically meaningful in spatial transcriptomics data.

Together, our GFT-based kernel design provides a principled way to approximate the spatial kernel while preserving the most biologically relevant patterns. By retaining only the low-frequency components (corresponding to the largest eigenvalues of *K*), we can construct approximations that maintain sensitivity to global and smooth spatial patterns while reducing computational complexity.

**Figure 2.**
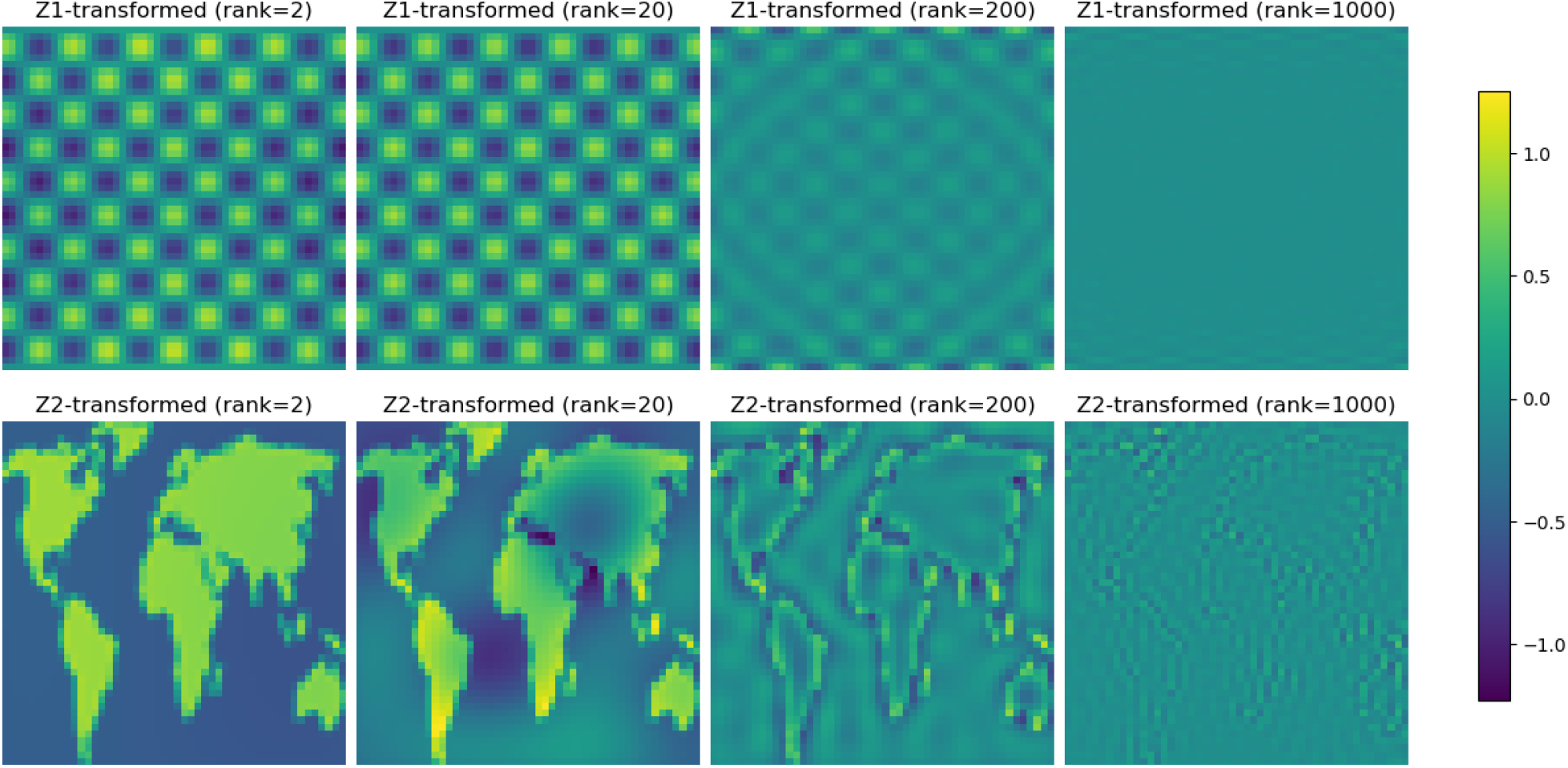
No-power transformation of spatial patterns on a 50 × 50 grid regarding GFT kernels with varying ranks. As rank increases, the undetectable patterns shift toward higher spatial frequencies, which are typically less biologically relevant in the context of spatial transcriptomics.

### 2.3 Kernel design and missing data handling for compositional data

After establishing our HSIC framework for spatial pattern detection and designing effective spatial kernels, we now address the challenges of analyzing isoform usage ratios — SPLISOSM’s primary focus. Specifically, we introduce an efficient independence testing approach for compositional data with varying sparsity patterns while maintaining test validity (**Theorem 2, Section 2.3.2**).

#### 2.3.1 Challenges of analyzing isoform usage ratio in spatial transcriptomics

Isoform usage ratios {*r*_1_, *r*_2_, …, *r*_*q*_} ∈ *S*^*q*^ for a gene with *q* isoforms form compositional data constrained to the simplex

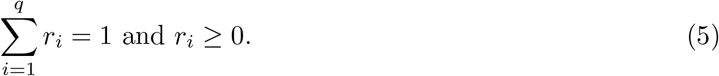

By analyzing ratios rather than raw expression, we focus on the relative transcript preference resulting from alternative processing of pre-mRNAs. Since increased preference for one isoform necessarily decreases preference for others, ratio variability should be treated holistically and tested at the gene level. Moreover, the simplex constraint in Equation (5) invalidates standard Euclidean data analysis methods. Traditional approaches rely on log-ratio transformations such as the additive log ratio (alr)

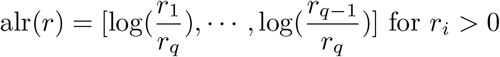

to map the simplex *S*^*q*^ to an isomorphic unconstrained space ℝ^*q*−1^. However, these transformations fail when encountering zeros, and workarounds like pseudo-counts introduce bias and distort relationships between observations.

Spatial transcriptomics presents two additional challenges:

1. **Zero coverage and undefined ratios**: The total UMI count for a gene at a particular spot is often zero^4^, leading to undefined ratios (NaN values).
2. **Discretization effect**: Even when observed, the ratios often appear binary due to limited sampling while the underlying biological preferences are more continuous[BANYL20], creating excessive zeros that further complicate log-ratio transformations.

#### 2.3.2 Efficient kernel independence test for ratios with undefined values

The simplest and unbiased approach to handle undefined ratios is to exclude spots with zero gene coverage from analysis. For kernel testing, this requires constructing a reduced spatial kernel *K*_*m*_ containing only the *m* spots with non-zero expression. However, this approach has two limitations: it discards potentially valuable biological information about regional gene expression, and becomes computationally expensive when testing thousands of genes with different sparsity patterns.

We develop an efficient testing framework that uses a single global spatial kernel *K*_*n*_ for all genes while properly controlling statistical properties. Our key insight is that gene-specific spatial kernels *K*_*m*_ can be derived as the principal submatrices of the global kernel: 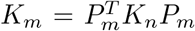, where *P*_*m*_ is the projection matrix selecting spots with non-zero coverage of a given gene. This ensures that zero-coverage spots still contribute to the spatial structure through their connections to observed spots. However, computing the spectrum of *K*_*m*_ for each gene remains expensive. We therefore introduce a method to efficiently approximate the gene-specific test using the global kernel.

##### Definition 2.5 (Zero-Padded Centered (ZPC) kernel).

*Let X*_*m*_ ∈ ℝ^*m*×*q*^ *be isoform ratios at m spots where ratios are defined, and let X*_*n*_ *be the extension of X*_*m*_ *with n*−*m additional rows containing undefined values (*nan*). Given a kernel function k* : *S*^*q*^ × *S*^*q*^ → ℝ *for compositional data, the ZPC kernel* 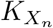 *is defined as:*

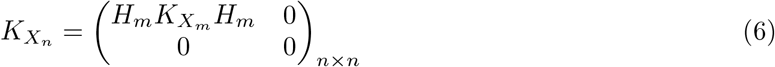

*where* 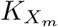 *with elements* 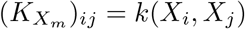 *is the kernel matrix for the m spots with defined ratios, and* 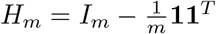 *centers this matrix*.

For a kernel *k*(*x, y*) = ⟨*ϕ*(*x*), *ϕ*(*y*)⟩ with feature map *ϕ* (e.g., log-ratio transformation), this construction effectively replaces undefined ratios with the mean in feature space 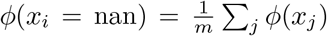 for defined *x*_*j*_. It enables an efficient testing procedure with rigorous statistical guarantees:

##### Theorem 2 (Mean-replacement preserves test validity).

*Let X*_*m*_ ∈ ℝ^*m*×*q*^ *represent isoform ratios at m spots where ratios are defined, X*_*n*_ *be its extension with undefined values, K*_*m*_ *be the centered compositional kernel for X*_*m*_, *and K*_*n*_ *be the corresponding ZPC kernel*.

*Let L*_*n*_ *be a spatial kernel for all n spots, and* 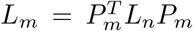 *be the m* × *m principal submatrix corresponding to spots with defined ratios. Consider the HSIC-based statistics*

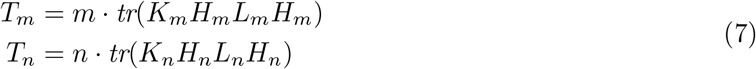

*where* 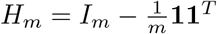 *and* 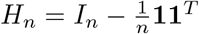 *are centering matrices*.

*The following properties hold:*

1. 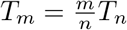, *allowing computation of the gene-specific statistic from the global statistic*.
2. *The asymptotic null threshold* 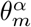 *where* 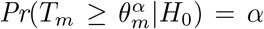 *can be bounded using only the spectrum of L*_*n*_.

*Proof*. First, we establish the relationship between *T*_*m*_ and *T*_*n*_. Note the projection matrix *P*_*m*_ satisfies

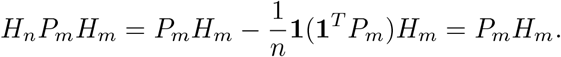

Removing the all-zero rows and columns in *K*_*n*_ that do not contribute to *T*_*n*_, we have

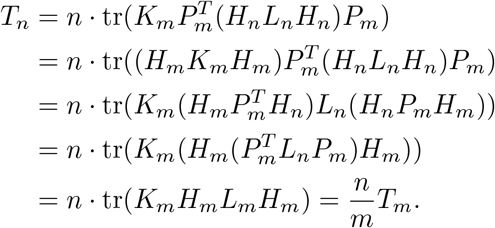

For the second claim, consider the eigenvalues of centered spatial kernels 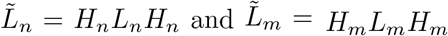, denoted as *λ*_1_ ≥ *λ*_2_ ≥ … ≥ *λ*_*n*_ and *µ*_1_ ≥ *µ*_2_ ≥ … ≥ *µ*_*m*_. Specifically, 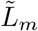 is a centered principal submatrix of 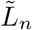

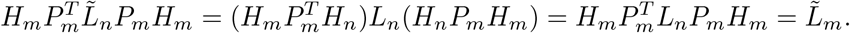

Applying Corollaries 1.1 and 1.2, we get the eigenvalue interlacing

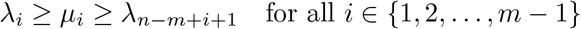

and *λ*_*m*_ ≥ *µ*_*m*_ = *λ*_*n*_ = 0 := *λ*_*n*+1_. Since *K*_*m*_ and its extension *K*_*n*_ share the same non-zero eigenvalues, for simplicity we assume it to be one (with multiplicity one).

Under the null hypothesis, *T*_*m*_ asymptotically follows a distribution 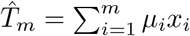 whare 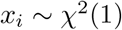 are independent. The interlacing property implies

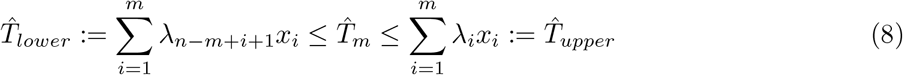

Therefore, the null threshold 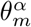 is bounded by corresponding quantiles of 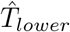 and 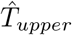, which depend only on the spectrum of the global kernel *L*_*n*_ and the number of non-zero spots *m*.

□

Theorem 2 provides a computationally efficient framework for testing spatial patterns in isoform usage. Rather than computing gene-specific kernels and their spectra for thousands of genes, we can: (1) compute *T*_*m*_ directly from *T*_*n*_ using the relationship 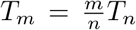, and (2) approximate the null distribution of *T*_*m*_ using the spectrum of the global spatial kernel *L*_*n*_.

In practice, using the upper bound from Equation (8) yields a conservative test with controlled false positive rate but reduced power. For exploratory analysis, we can alternatively approximate the null distribution of *T*_*m*_ with

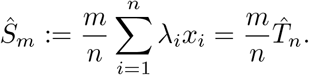

This essentially corresponds to a two-step procedure where we first replace undefined values with feature means after transformation, then test with *n* spots as if all ratios were observed. The quality of this approximation relates to the bounds in Equation (8) through the majorization relationship

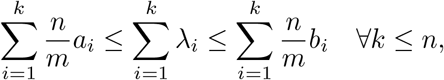

where *a* = {*λ*_*n*−*m*+2_ ≥ … ≥ *λ*_*n*_ = 0 = …= 0} ∈ R^*n*^ and *b* = {*λ*_1_ ≥ … ≥ *λ*_*m*_ ≥ 0 = … = 0} ∈ R^*n*^ are the weights used in constructing the bounds in Equation (8).

Our empirical simulations show that *Ŝ*_*m*_ yields similar power and well-calibrated type I error compared to the ground truth gene-specific null 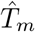.

#### 2.3.3 Kernel transformation for near-binary ratios

Having addressed undefined values with our ZPC kernel, we now consider the optimal kernel function for compositional data with near-binary ratios. As noted earlier, spatial transcriptomics data often exhibit discrete, binary-like ratios because of the limited sequencing depth.

We evaluated several transformation approaches for the kernel function *k*(*x, y*) = ⟨*ϕ*(*x*), *ϕ*(*y*)⟩, including standard log-ratio transformations (alr, clr, ilr) and a recently proposed radial transformation[PYPA22] 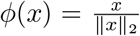. All log-based methods required pseudocounts (i.e. replacing zeros with *ϵ*s and then renormalizing ratios to sum to one) that distort the original composition. Moreover, the extreme values produced by transforming near-binary ratios dominated the analysis after centering, whose scale is only controlled by the pseudocount and does not have biological significance.

Indeed, our empirical evaluations demonstrated that the simple linear kernel *k*(*x, y*) = ⟨*x, y*⟩ without transformation provided optimal performance. This approach naturally accommodates zeros, avoids extreme value problems, and maintains appropriate distance relationships between near-binary observations. When combined with our ZPC kernel construction, it creates a robust, parameter-free framework for detecting spatial patterns in isoform usage that balances statistical power with proper type I error control. In contrast, the radial transformation always leads to inflated p-values under the null.

### 2.4 Conditional kernel independence test for spatial association analysis

After detecting spatial variability in isoform usage, we now aim to identify the molecular mechanisms underlying the observed patterns. Our HSIC framework naturally extends to test associations between isoform usage (*X*) and biological covariates (*Y*) such as spatial domains or RNA binding protein expression. While HSIC testing is non-directional, biological knowledge often suggests causal interpretations — spatial variation in splicing factors likely drives region-specific splicing patterns rather than vice versa.

To our knowledge, SPLISOSM provides the first differential association testing framework that explicitly *conditions on spatial autocorrelation*, controlling for false positives that would otherwise arise from shared spatial structure. These association tests can integrate into broader causal discovery frameworks to systematically identify regulatory networks that drive isoform usage dynamics.

#### 2.4.1 From unconditional to conditional kernel independence test

With linear kernels, the HSIC statistic 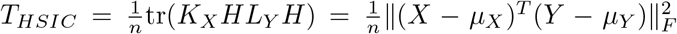 is proportional to the RV coefficient, which measures multivariate linear correlations between *X* and *Y*.

For binary *Y* with *m* positive samples, we have

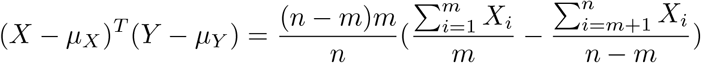

effectively functioning as a multivariate extension of the two-sample t-test.

However, this approach cannot distinguish true biological associations from spatial confounding — *X* and *Y* may share spatial patterns simply due to mutual dependence on spatial coordinates without direct relationship. Traditional gene regulatory network approaches handle confounding by conditioning on cell type (*X* ⊥ *Y* |*C*), but the continuous nature of spatial data creates a fundamental challenge: for each location *S*_*i*_, we have only one observation (*X*_*i*_, *Y*_*i*_), making direct estimation of conditional distributions impossible.

#### 2.4.2 Conditional kernel independence test

To overcome this challenge, we adapt the conditional independence test from [ZPJS12], which shares information across observations by learning regression functions *f*_*X*_ : *S* → *X* and *f*_*Y*_ : *S* → *Y*, then testing the independence of the residuals.

For a linear ridge regression with L2 regularization *λ*, we have the estimator

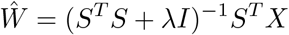

where *S* ∈ ℝ^*n*×2 or 3^ represents spatial coordinates, *X* ∈ ℝ^*n*×*d*^ the spatial variable, and *W* ∈ ℝ^2 or 3×*d*^ the regression coefficient mapping *S* to *X*. The residual 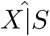 becomes

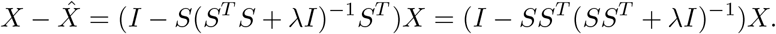

Substituting the linear kernel *SS*^*T*^ with general kernels *K*_*S*_ extends this to non-linear relationships

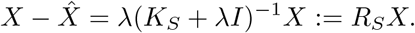

This formulation allows us to construct a kernel for the conditional *X*|*S* as *K*_*X*|*S*_ := *R*_*S*_*K*_*X*_*R*_*S*_ where *R*_*S*_ = *λ*(*K*_*S*_ + *λI*)^−1^, given a spatial kernel *K*_*S*_ and the target kernel *K*_*X*_. The parameter *λ* controls conditioning strength: *λ* = 0 implies *X* is completely explained by *S*, leaving no information for *X*|*S*; while *λ* → ∞ implies 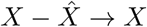 which gives the unconditional test. In practice, we estimate *λ* and *K*_*S*_ individually for each variable using Gaussian Process Regression with RBF and white kernels.

Using again linear kernels for *X*|*S* and *Y* |*S*, our conditional association test statistic becomes

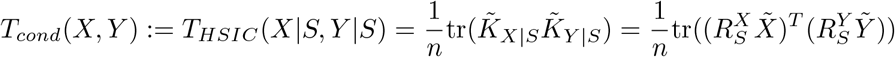

where 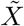 and 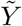 are column-wise centered, and 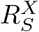 and 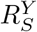 are the estimated residual operators from Gaussian Process regression. The asymptotic null is computed similarly following Equation (1).

### 2.5 Implementation details

This section outlines SPLISOSM’s implementation for nonparametric spatial variability (SV) and differential isoform usage (DU) tests. We first present the construction of the full-rank ICAR kernel, followed by the core algorithms for three full-rank SV tests: HSIC-GC (gene expression), HSIC-IR (isoform usage ratio), and HSIC-IC (isoform expression). We then describe their computationally efficient low-rank versions using the GFT kernel approximation (Definition 2.4). Finally, we detail the implementation of HSIC-based unconditional and conditional association tests for differential isoform usage analysis.

#### 2.5.1 Full-rank spatial variability tests

##### Algorithm 1 Construction of ICAR Spatial Kernel (Definition 2.3)

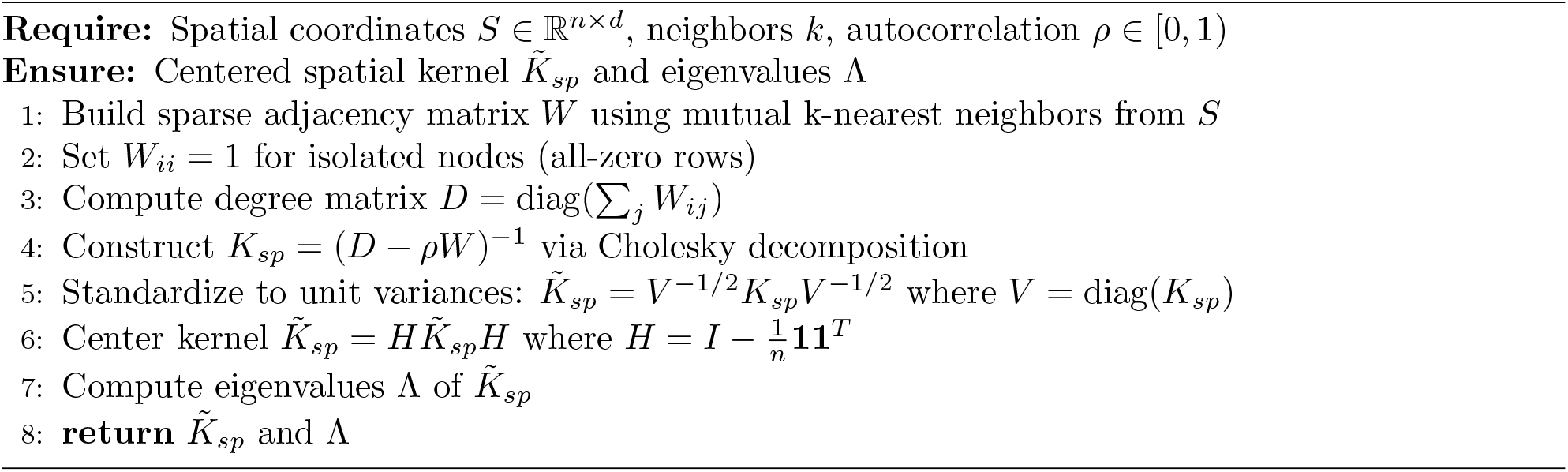

The computation bottleneck of Algorithm 1 is the construction of the dense *n* × *n* kernel (*O*(*n*^2^) space, *O*(*n*^3^) time) and its eigendecomposition (*O*(*n*^3^) time). For 10X Visium data (*n* ≈ 5000), this requires around 1GB RAM and 11 seconds on an M1 MacBook Air.

##### Algorithm 2 HSIC-GC: Spatial Variability Test for Gene Expression

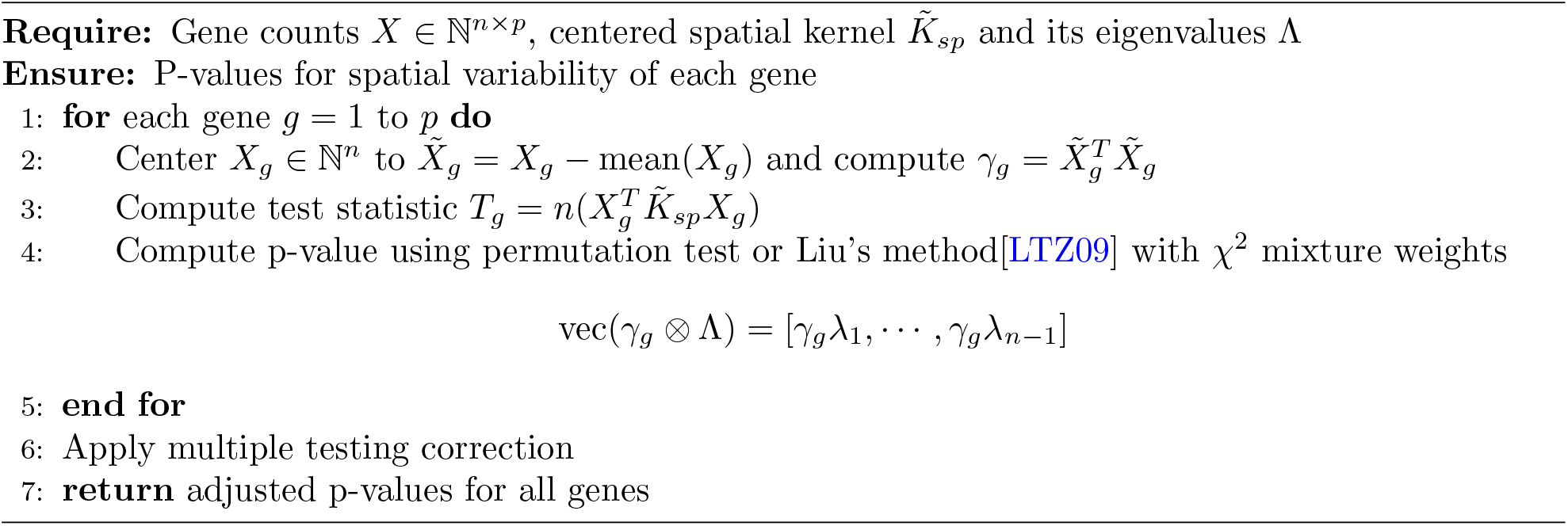

The key difference between HSIC-GC (Algorithm 2) and SPARK-X[ZSZ21] is the design of the spatial kernel. Our ICAR kernel (from Algorithm 1) is full-rank and has greater statistical power. Also, HSIC-GC does not test specific spatial patterns (i.e. the 11 transformations in SPARK-X) individually.

##### Algorithm 3 HSIC-IC: Spatial Variability Test for Isoform Expression

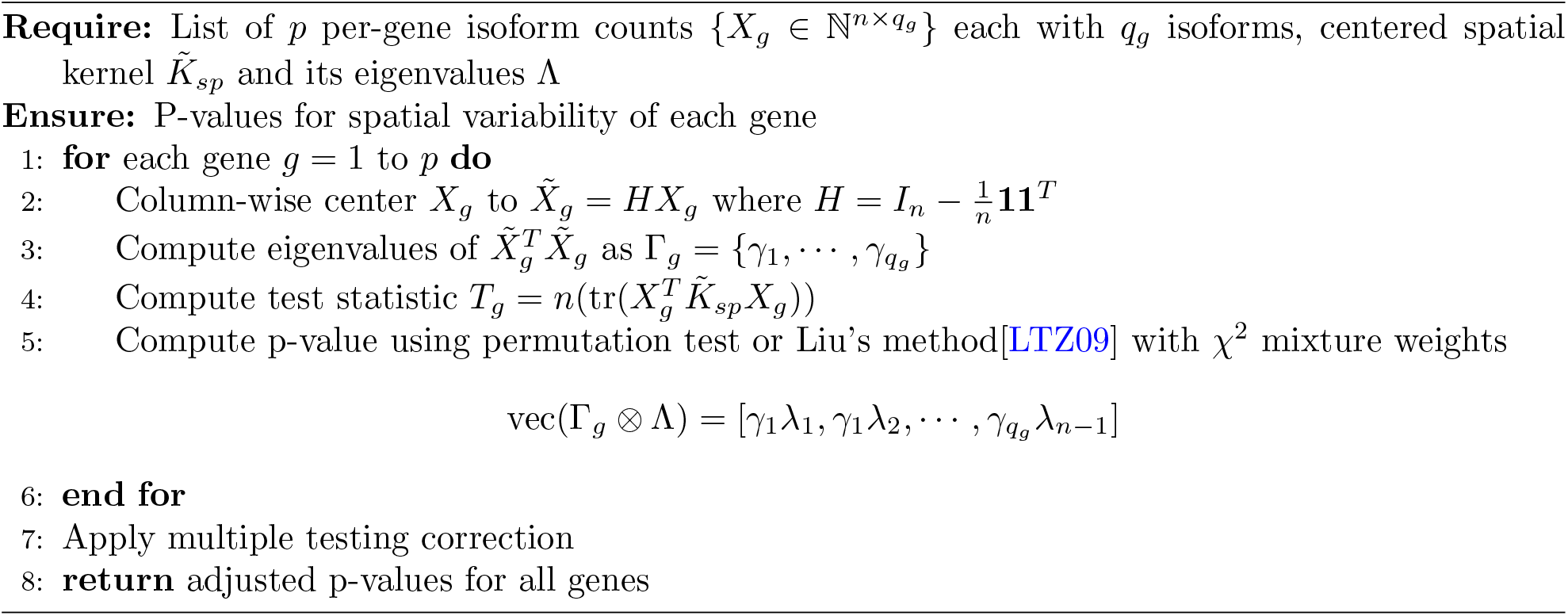

HSIC-IC (Algorithm 3) is the multivariate generalization of HSIC-GC. With linear kernel *K*_*X*_ = *XX*^*T*^, it is equivalent to testing each isoforms (columns) in *X* individually and then combining them into a single p-value per gene with a specific asymptotic null. By design, it can also test spatial variability in other multivariate quantities like pathway or gene program activity.

##### Algorithm 4 HSIC-IR: Spatial Variability Test for Isoform Usage Ratio

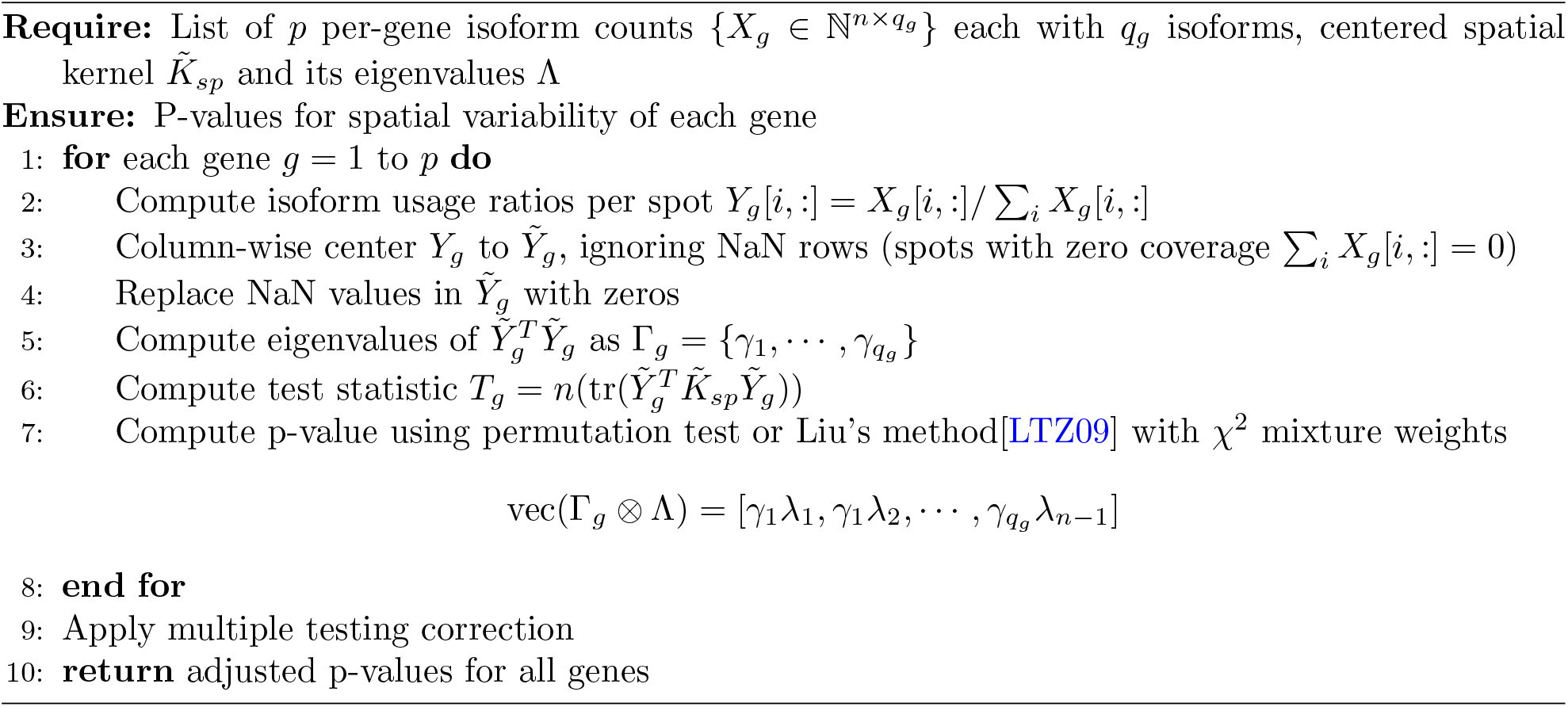

HSIC-IR (Algorithm 4) is the generalization of HSIC-IC for compositional data where NaN values are replaced by the mean (zeros after centering) to handle zero coverage. The linear kernel shows comparable results to more complex compositional kernels (i.e. log-ratio transformations) in practice. By design, it can also test spatial variability in other multivariate quantities with missing data points.

#### 2.5.2 GFT-based low-rank spatial variability tests

##### Algorithm 5 Construction of Low-rank GFT Spatial Kernel (Definition 2.4)

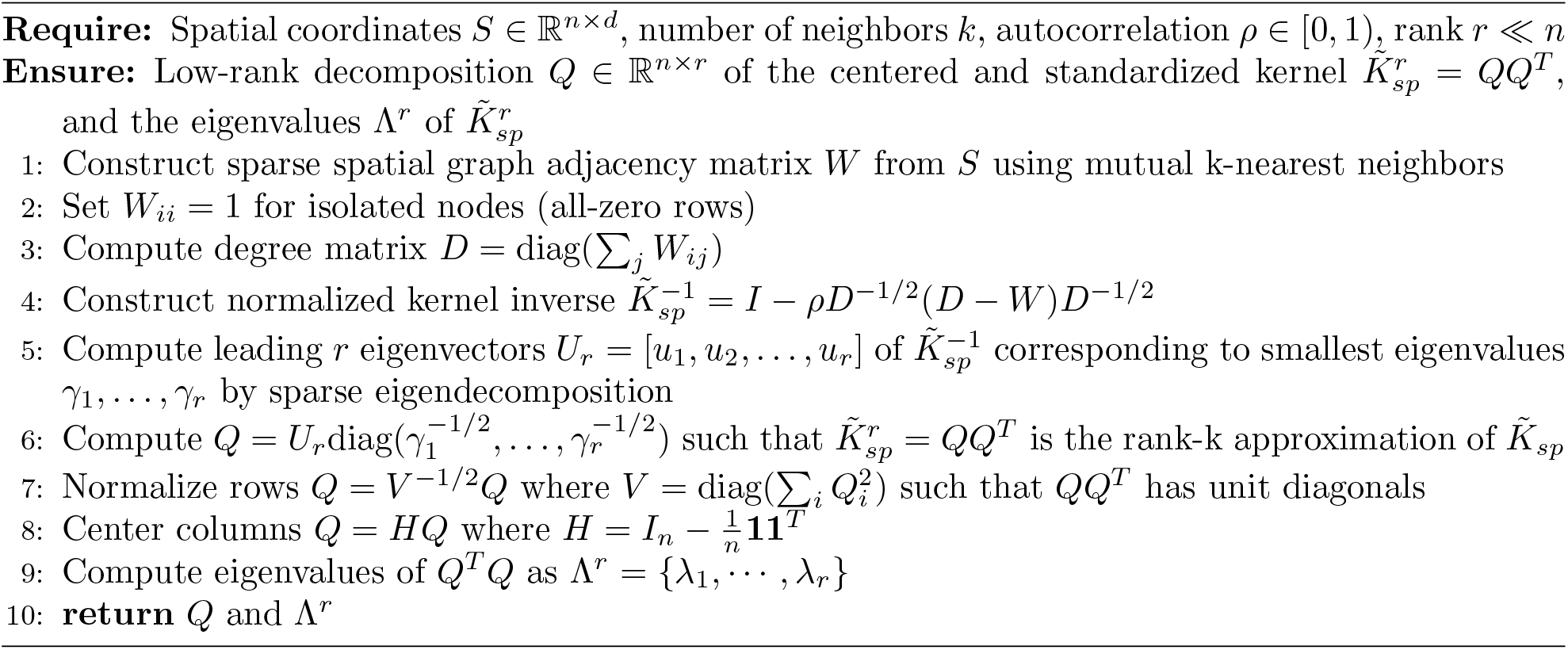

The computational bottleneck of Algorithm 5 is now the sparse eigendecomposition with complexity *O*(*rkn*), which scales linearly with sample size. Replacing 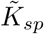 with *QQ*^*T*^ and Λ with Λ^*r*^ in Algorithms 2, 3, and 4 gives their low-rank versions. Note the dense kernel *QQ*^*T*^ is never explicitly formed since 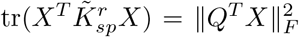. For Slideseq-V2 data (*n* ≈ 50000, *r* = 100), the low-rank SV tests require around 2GB RAM and takes 1 minute to complete on an M1 MacBook Air. See Figure 3 for performance comparison on the mouse hippocampus dataset.

**Figure 3.**
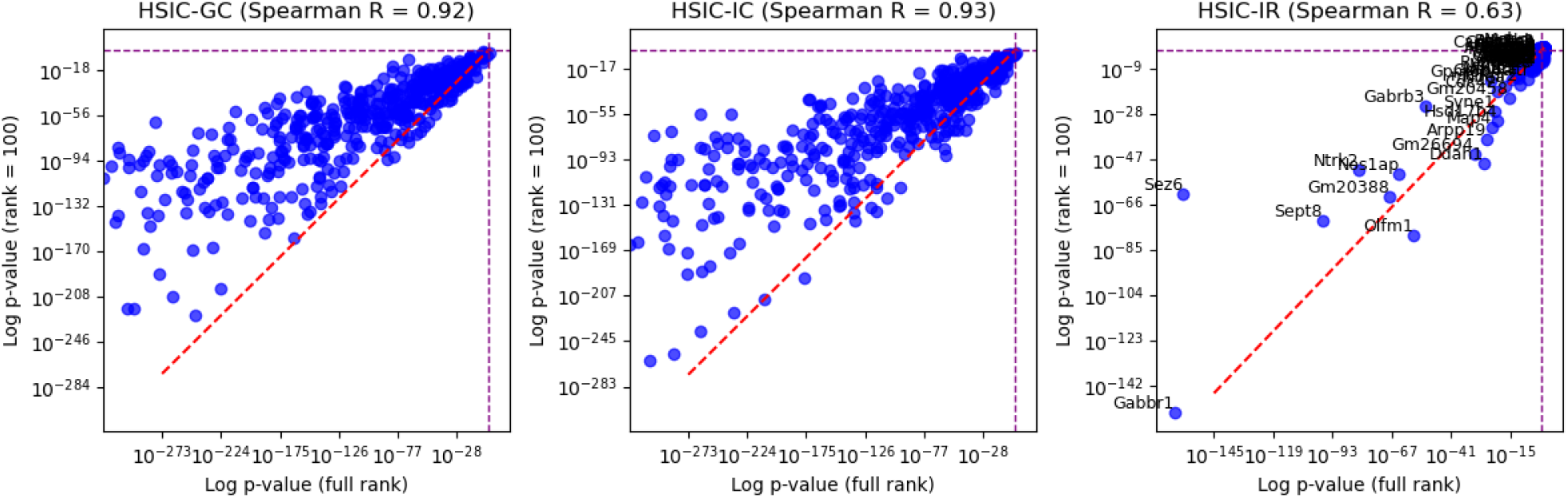
Performance of low-rank spatial variability tests on a Slideseq-V2 dataset of mouse hip-pocampus (*n* = 53208, *r* = 100). Low-rank tests produce consist ranking of genes by their variability but have reduced statistical power.

#### 2.5.3 HSIC-based differential isoform usage tests

##### Algorithm 6 Unconditional Differential Isoform Usage Test

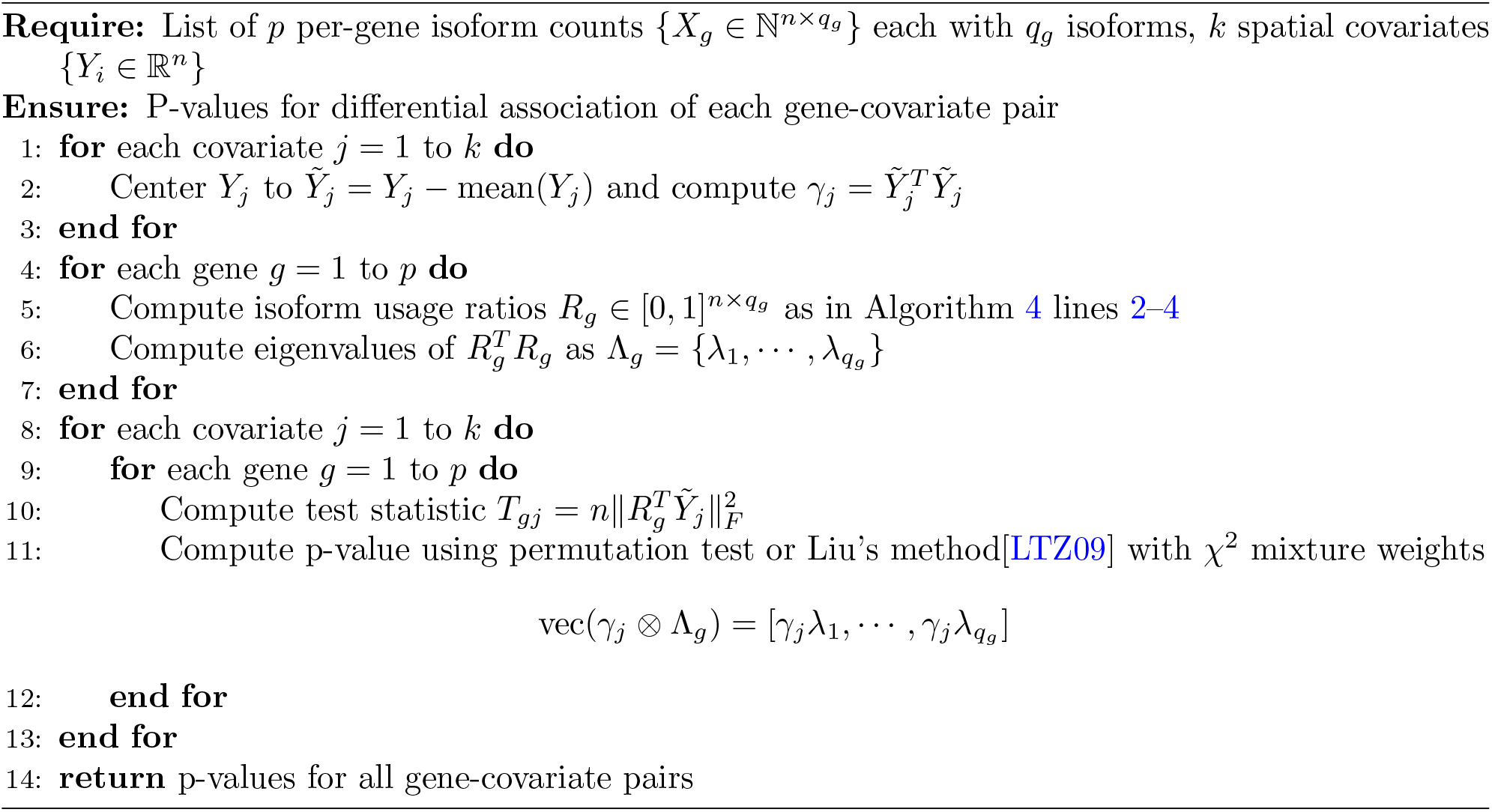

Algorithm 6 tests for associations between isoform usage patterns and biological covariates without accounting for spatial confounding. This direct approach identifies strong associations but may include false positives due to shared spatial structure. With linear kernels it is both fast and memory efficient.

##### Algorithm 7 Conditional Differential Isoform Usage Test

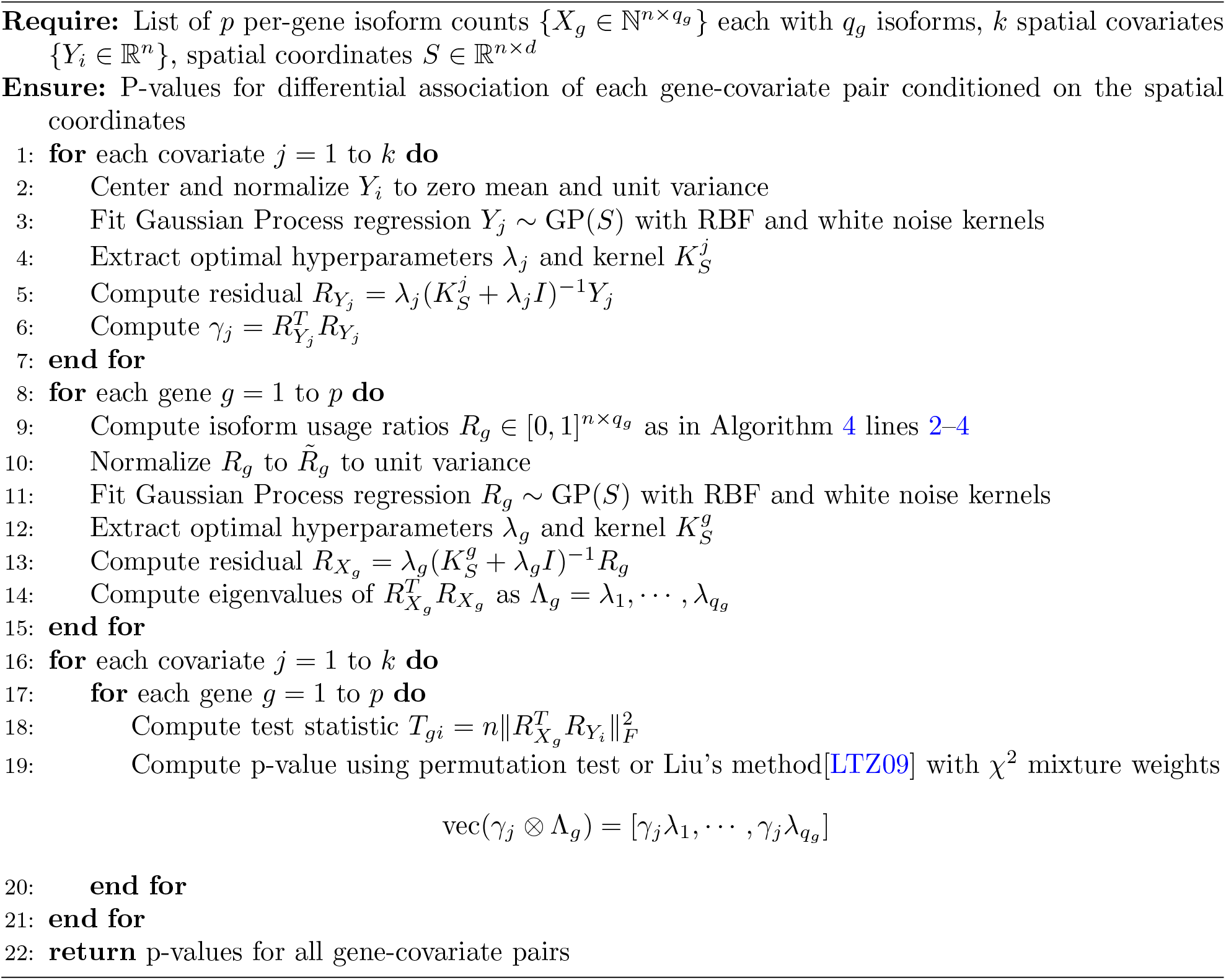

Algorithm 7 represents a key innovation in SPLISOSM, which aims to identify associations that cannot be explained by shared spatial structure alone. The Gaussian Process regression with RBF kernels captures complex, non-linear spatial patterns, while the hyperparameter *λ* automatically determines the appropriate level of conditioning based on how strongly each variable correlates with spatial location. Our empirical results show that a global *λ* and spatial kernel *K*_*S*_ is not sufficient to reduce inflated p-values under variable autoregressive noise (Figure 4, the first column). The computational bottleneck is thus the fitting of per-gene and per-covariate Gaussian Process models.

**Figure 4.**
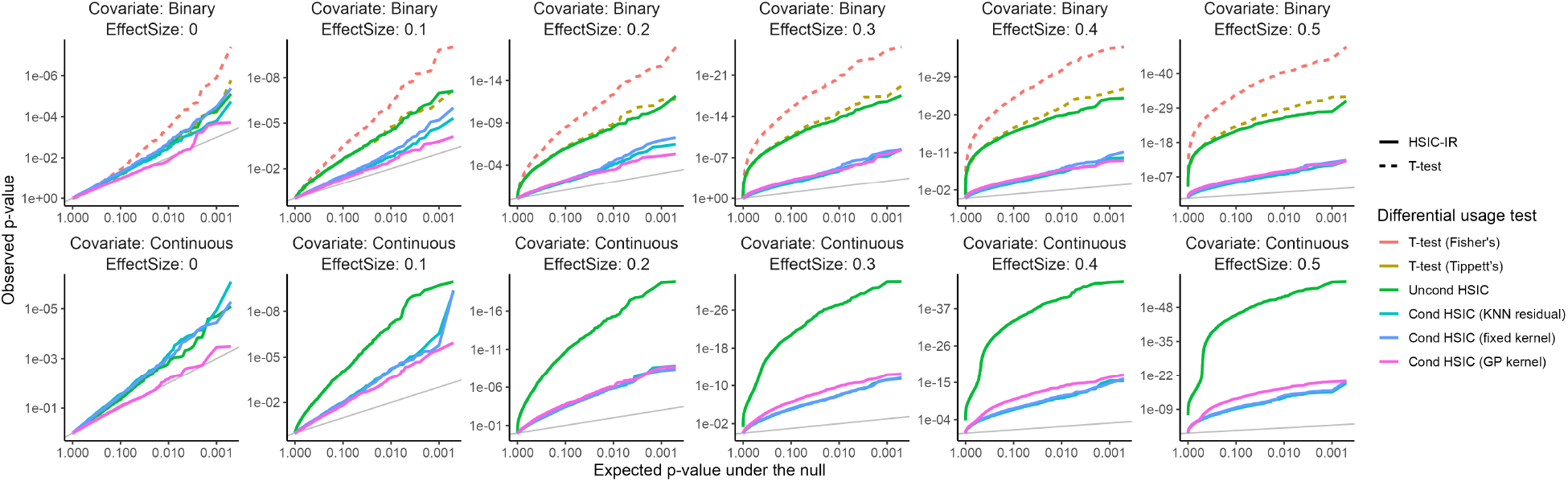
Only the conditional HSIC with the adaptive GP regression model (pink line) can reduce the inflated p-value under the null hypothesis (the first column).

### 2.6 Comparison with existing spatial isoform analysis approaches

While SPLISOSM represents the first statistically rigorous framework for spatial isoform variability testing, several prior studies have attempted to analyze transcript diversity from spatial transcriptomics data using various approaches. Here we provide a brief review clarifying their limitations and explaining why these methods are not directly comparable to SPLISOSM.

#### 2.6.1 Differential usage-based approaches

These methods require predefined spatial regions or annotations to compare isoform usage between domains, which aligns closer to SPLISOSM’s differential usage test, though none explicitly consider spatial autocorrelation.

##### **sl-ISO-Seq** [JPM^+^21]

This is one of the first studies presenting spatial long-read transcriptomics data. For isoform switching detection, the authors aggregated all isoform counts per predefined brain region and applied a gene-level chi-squared test for differential usage.

##### **SiT** [LBT^+^23]

This is one of the first studies presenting spatial long-read transcriptomics data. The SiT analysis pipeline first identified isoform markers for each region, then designated genes with different isoform markers across regions as variably spliced.

##### **Patho-DBiT** [BZG^+^24]

When analyzing spatial splicing, the authors aggregated all reads within anatomical regions and applied rMATS, a bulk RNA-seq differential splicing tool.

##### **Longcell** [FKR^+^25]

Although primarily improving the preprocessing of single-cell and spatial long-read data, Longcell also comes with a single-cell-based differential usage test based on a hierarchical Beta-Binomial model. However, without explicitly accounting for spatial autocorrelation, the test would again suffer from inflated false positive rates from spurious associations.

#### 2.6.2 Heuristic-based approaches

These methods reduce multivariate compositional isoform usage to univariate per-gene metrics, fundamentally limiting their statistical power compared to SPLISOSM’s multivariate approach.

##### **stAPAminer** [JTZ^+^23]

This tool quantifies annotated poly(A) site expression from spatial transcriptomics data but employs several questionable statistical practices for isoform variability analysis: (1) it applies local spatial smoothing as an imputation step, artificially making every gene appear spatially variable, (2) it multiplies usage ratios by 10 and treats them as count data, and (3) it applies SPARK, which was designed for univariate gene expression rather than compositional isoform ratios. These limitations severely compromise the validity of its statistical inference.

##### **SpliZ-based spatial analysis** [OS23]

This more rigorous approach computes a univariate SpliZ score per gene at each spatial location from short-read sequencing data, then quantifies spatial variability using Moran’s I. This approach has limited sensitivity because: (1) SpliZ scores are computed only from junction-spanning reads, which drastically reduces the number of analyzable genes in sparse data, (2) collapsing multivariate usage to a single score loses information. This explains why their analysis identified fewer than 20 spatially variable splicing genes in adult mouse brain, compared to over 1,000 identified by SPLISOSM.

## 3 Parametric spatial pattern discovery using generalized linear mixed model (GLMM)

While non-parametric kernel independence tests show robust performance in spatial pattern detection, parametric approaches can provide complementary insights through explicit data modeling. For example, we may fit a probabilistic model to infer the noise-free latent isoform usage preferences, generate useful visualizations, and link isoform patterns to spatial phenotypes in a generative process.

This section introduces a generalized linear mixed model (GLMM) framework for spatial isoform analysis. However, unlike univariate linear mixed models commonly used for gene expression[STS18, SZZ20], multivariate GLMM is substantially harder to fit. Our empirical evidence shows that *GLMM-based spatial variability detection is not reliable on sparse data* due to inaccurate model fitting especially for random effects. We therefore recommend using this approach *only* for differential usage testing and visualization.

### 3.1 From generalized linear model (GLM) to generalized linear mixed model (GLMM)

Conventional alternative splicing analysis typically models gene isoform expression using a multinomial distribution

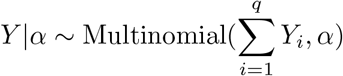

where 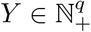 represents observed counts of *q* isoforms and *α* ∈ [0, 1]^*q*^ the latent isoform usage ratios. The relationship between isoform preferences and covariates can be expressed through a generalized linear model

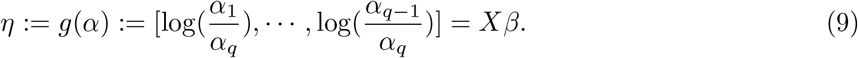

Here, *X* ∈ ℝ^*k*^ represents covariates, *β* ∈ ℝ^*k*×(*q*−1)^ contains regression coefficients, and *g* is the log-ratio transformation that maps constrained ratios *α* from the simplex *S*^*q*^ to unconstrained space *η* ∈ ℝ^*q*−1^. Through this link function, the expected counts 𝔼 [*Y*] := *µ* = *α* ∑_*i*_ *Y*_*i*_ = *g*^−1^(*η*) ∑ _*i*_ *Y*_*i*_ become a function of the model parameters, with partial derivatives 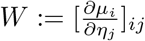 forming a (*q* − 1) × (*q* − 1) matrix.

Standard GLM approaches assume spatial patterns in *Y* derive solely from covariates *X*, potentially leading to inflated estimates and false discoveries. To address this (analogous to the challenges addressed by our conditional HSIC tests), we extend GLM to a generalized linear mixed model (GLMM) with spatial random effects *ν*:

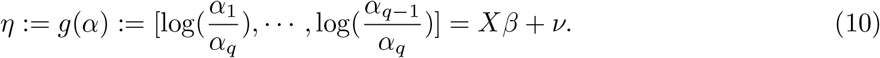

For simplicity, we assume independent random effects across isoforms

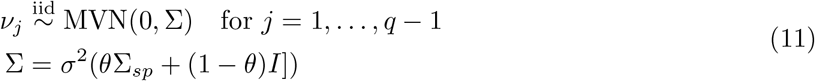

where Σ mixes spatial covariance Σ_*sp*_ (e.g., the ICAR kernel) and white noise.

### 3.2 Problem formulation

Under the GLMM framework, we can formulate the key tasks of isoform analysis as:

- **Spatial variability testing**: Given isoform-level quantification of *n* observations 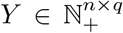 (with *X* = ∅), test whether the spatial variance component is non-zero

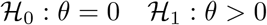
- **Differential usage testing** Given isoform-level quantification of *n* observations 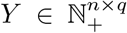 and covariates *X* ∈ ℝ^*n*×*k*^, test whether the effect size *β* is zero while accounting for spatial autocorrelation

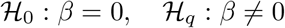

Note that the two objectives require fitting different GLMM models (with or without fixed effects).

The GLMM also enables estimation of underlying latent isoform usage ratios

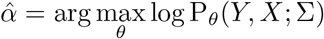

where *θ* = {*β, ν*} includes both fixed and random effects.

### 3.3 Model inference

#### 3.3.1 Fitting GLM using iteratively reweighted least squares (IWLS)

Denote *L*(*β*) := *f* (*Y, X, β*) the log-likelihood function of the GLM model. To compute the (vectorized) maximized likelihood estimator (MLE) 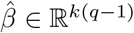 for GLM, we apply Newton’s method

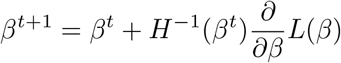

where *H*(*β*^*t*^) is the Hessian matrix 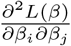 of shape *k*(*q* − 1) × *k*(*q* − 1) evaluated at *β*^*t*^. With *n* observations, this update can be reformulated as iteratively reweighted least squares (IWLS)

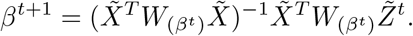

The weight matrix *W* = diag(*W*_1_, *W*_2_, …, *W*_*n*_) is block diagonal, where 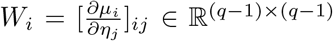 contains partial derivatives of the mean-link relationship computed for the *i*-th observation. 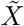 is the expanded design matrix of size *n*(*q* − 1) × *k*(*q* − 1) repeating *X* ∈ ℝ^*n*×*k*^, and 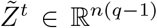 is the vectorized version of the working variable *Z*^*t*^ ∈ ℝ^*n*×(*q*−1)^

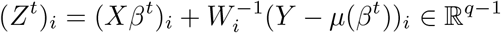

for each observation *i* = 1, …, *n*.

#### 3.3.2 Fitting GLMM using Laplace approximation

For GLMM, we need to integrate out random effects *ν* to obtain maximum likelihood estimates

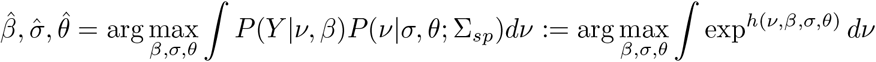

where *h*(*ν, β, σ, θ*) is the log joint likelihood function. Since *Y* follows a non-Gaussian distribution, this integral is computationally intractable. Instead, we approximate the marginal log-likelihood function using second-order Laplace approximation at the mode 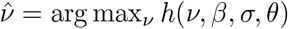, yielding

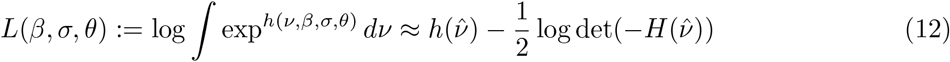

where *H*(*ν*) is the Hessian matrix of *h*(*ν*) with shape *n*(*q* − 1) × *n*(*q* − 1).

The joint likelihood (and thus Hessian) can be decomposed into multinomial likelihood and multivariate Gaussian (MVN) prior components following *h*(*ν*) = log P_*ν,β*_(*Y*) + log P_*σ,θ*_(*ν*; Σ_*sp*_). Denote *ν* ∈ ℝ^*n*×(*q*−1)^ with *n* observations in matrix form. For the multinomial term, note *ν*_*i*:_ and *ν*_*j*:_ ∈ R^*q*−1^ of any two spatial locations *i* and *j* are independent. Within a particular location *i*, let 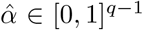 be the first *q* − 1 elements of the usage ratio *α*_*i*_(*ν*_*i*_, *β*) ∈ [0, 1]^*q*^, we have

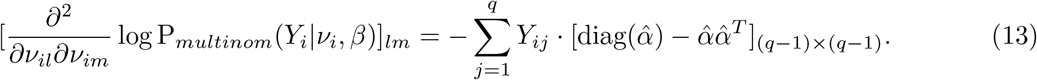

For the MVN prior term, note *ν*_:*l*_ and *ν*_:*m*_ ∈ R^*n*^ of any two isoforms *l* and *m* are independent. Within a particular isoform *l*, we have

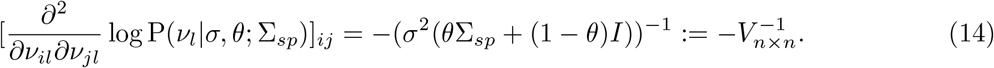

Combining Equations (13) and (14) we have all the non-zero entries of the Hessian *H*(*ν*).

Given 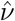 and 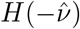, we maximize Equation (12) using gradient descent. 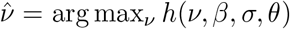 is updated by either Newton’s method (with 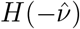) every *k* epoch, or by gradient descent where we jointly optimize *ν, β, σ, θ*

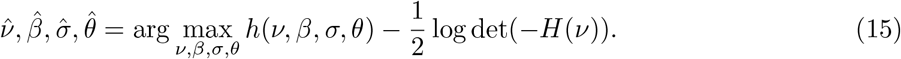

Due to frequent Hessian evaluation, the optimization is computationally intensive and can be numerically unstable. Since our primary interest is in likelihood ratios for hypothesis testing rather than absolute likelihoods, we also implement an h-likelihood approach[JL21] that optimizes the joint likelihood directly

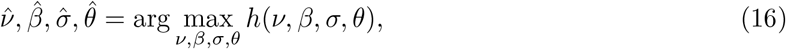

reflecting the assumption that, under the null hypothesis, 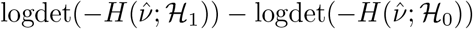 is asymptotically neglectable[LN96]. We solve (16) using gradient descent with a small lower bound on *σ* to avoid zero variance.

### 3.4 Hypothesis testing

We encountered several setbacks when exploring GLMM-based hypothesis testing for spatial isoform analysis. The accurate estimation of model parameters, particularly random effects, proves difficult with sparse spatial transcriptomics data. Consequently, most testing procedures depending on alternative model inference (involving spatial variance proportion *θ* or fixed effect size *β*) were unreliable without substantial modifications. The GLM and GLMM differential usage tests based on the score statistic, which is estimated under the null (*β* = 0), are relatively calibrated. We document these exploratory findings below for reference in future research.

#### 3.4.1 Inaccurate random effect estimation in GLMM

After model fitting, we can estimate underlying isoform usage ratios as

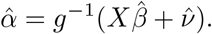

While random effects improve ratio inference compared to simple GLM (Figure 5), GLMM-full (with learnable *θ*) estimates are only marginally better than models using just white noise (GLMM-null *θ* = 0), suggesting difficulties in capturing spatial structure. Furthermore, the estimated variance 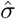 often shrinks toward zero, reflecting the challenge of estimating dispersion *ν* that contains *n* · (*q* − 1) parameters from sparse observations where multinomial alone may sufficiently explain data variation.

Due to these vanishing random effects, GLMM models fail to correct inflated fixed effect estimates *β* when both fixed and random effects are present in the model (Figure 6). Estimates deviate substantially from ground truth even when no covariate-driven pattern exists in the simulation.

#### 3.4.2 Likelihood-ratio tests compromised by local optima

For spatial variability testing, we fit a GLMM model without covariates (*X* = ∅). When covariate-induced variability exists, the model attributes it to spatial noise components as expected 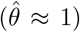, suggesting the capability of the *θ*-based SV test to capture effects from unknown latent factors (e.g. cell types). However, the marginal likelihood suffers from approximation errors and local optima (Figure 7), causing the likelihood ratio statistic 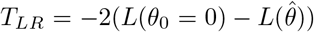 to deviate significantly from its asymptotic *χ*^2^ distribution.

We implement a permutation test by randomly shuffling spatial coordinates to calculate empirical p-values from *T*_*LR*_. Still, the GLMM-based approach remains computationally expensive (even more so with permutation) and exhibits considerably lower power than our non-parametric HSIC-based test.

#### 3.4.3 Effective differential usage testing with score statistics

Despite these unsuccessful attempts, the observation that GLMM models can properly attribute spatial variability to random effects when estimated without covariates indicates their capability to control for spatial confounding when testing under the null hypothesis *β*_0_ = 0. We validate this idea by comparing differential usage test performance between Wald and score tests

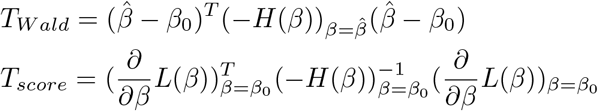

**Figure 5.**
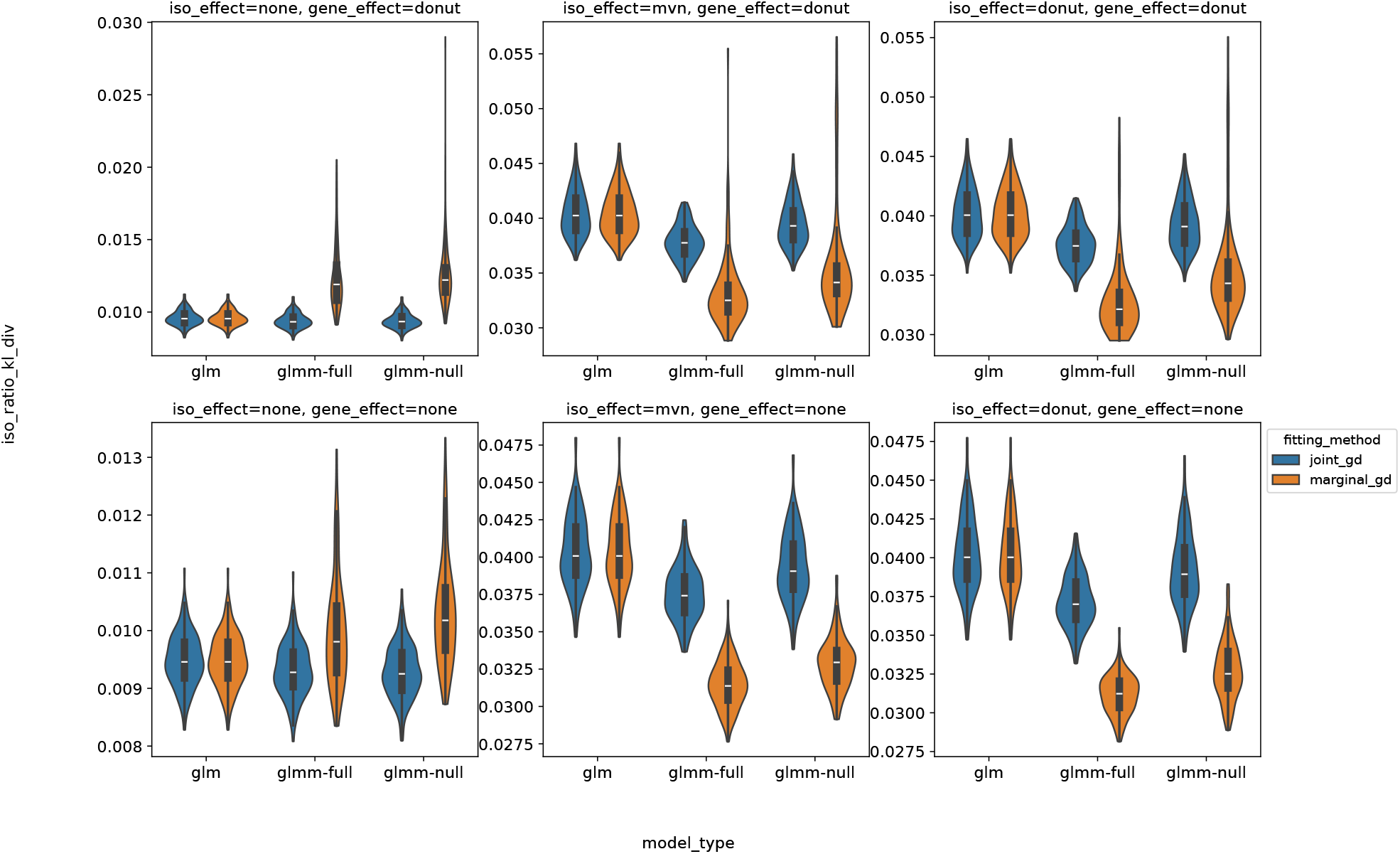
Per-spot isoform ratio inference accuracy with covariates *X*. Y-axis shows KL divergence to ground truth ratios (lower is better). The GLM model uses a global ratio for all spots, GLMM-full incorporates learnable spatial variance, while GLMM-null uses only white noise (*θ* = 0). ‘joint-gd’ and ‘marginal-gd’ refer to different optimization approaches in Equations (16) and (12) (only applicable to GLMM models).

**Figure 6.**
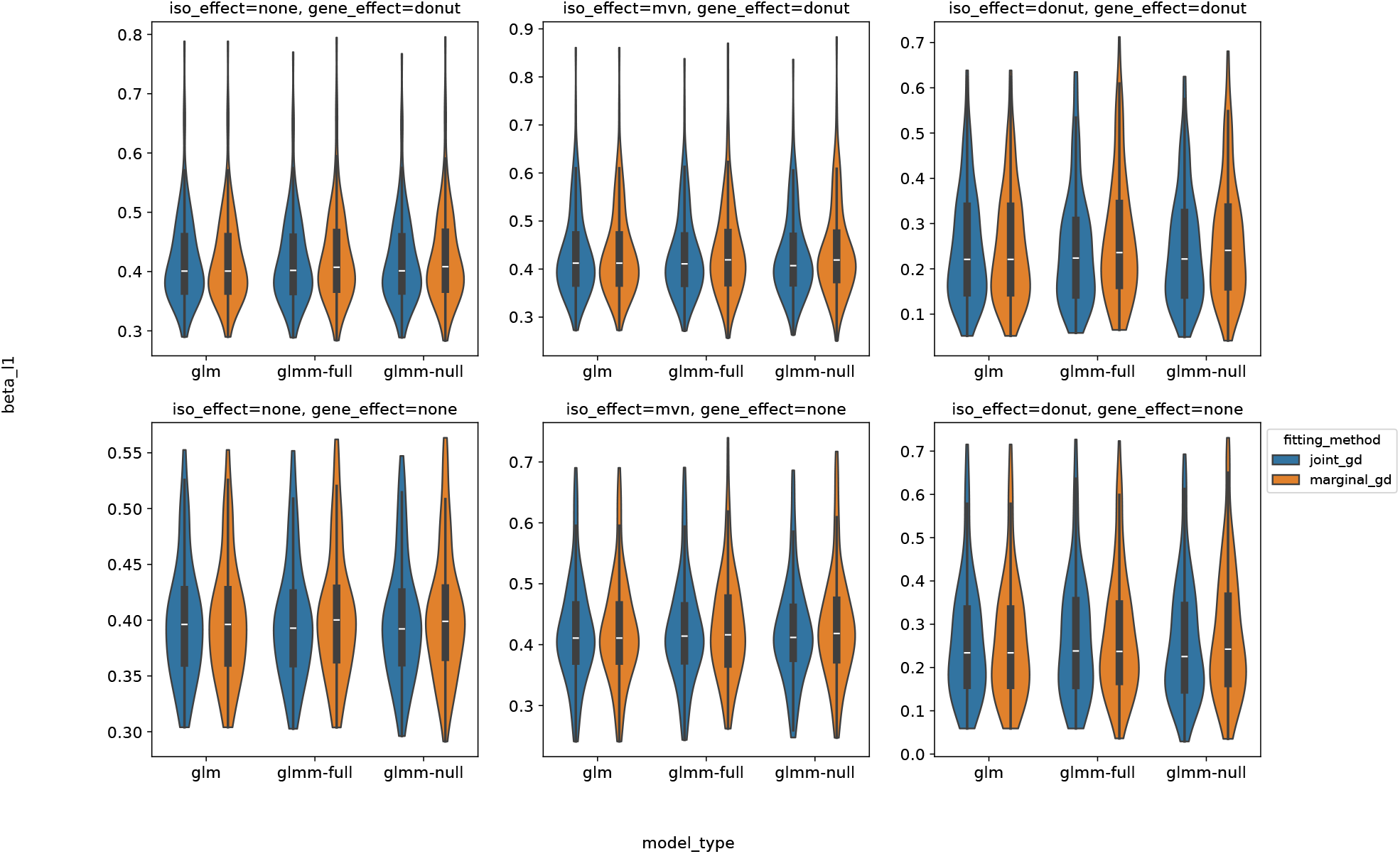
Fixed effect size *β* estimation accuracy. Y-axis shows L1 loss to ground truth (lower is better). The left and center columns represent data with no covariate-driven patterns, yet all models, including GLM, learn spurious non-zero *β* from confounding spatial variability.

**Figure 7.**
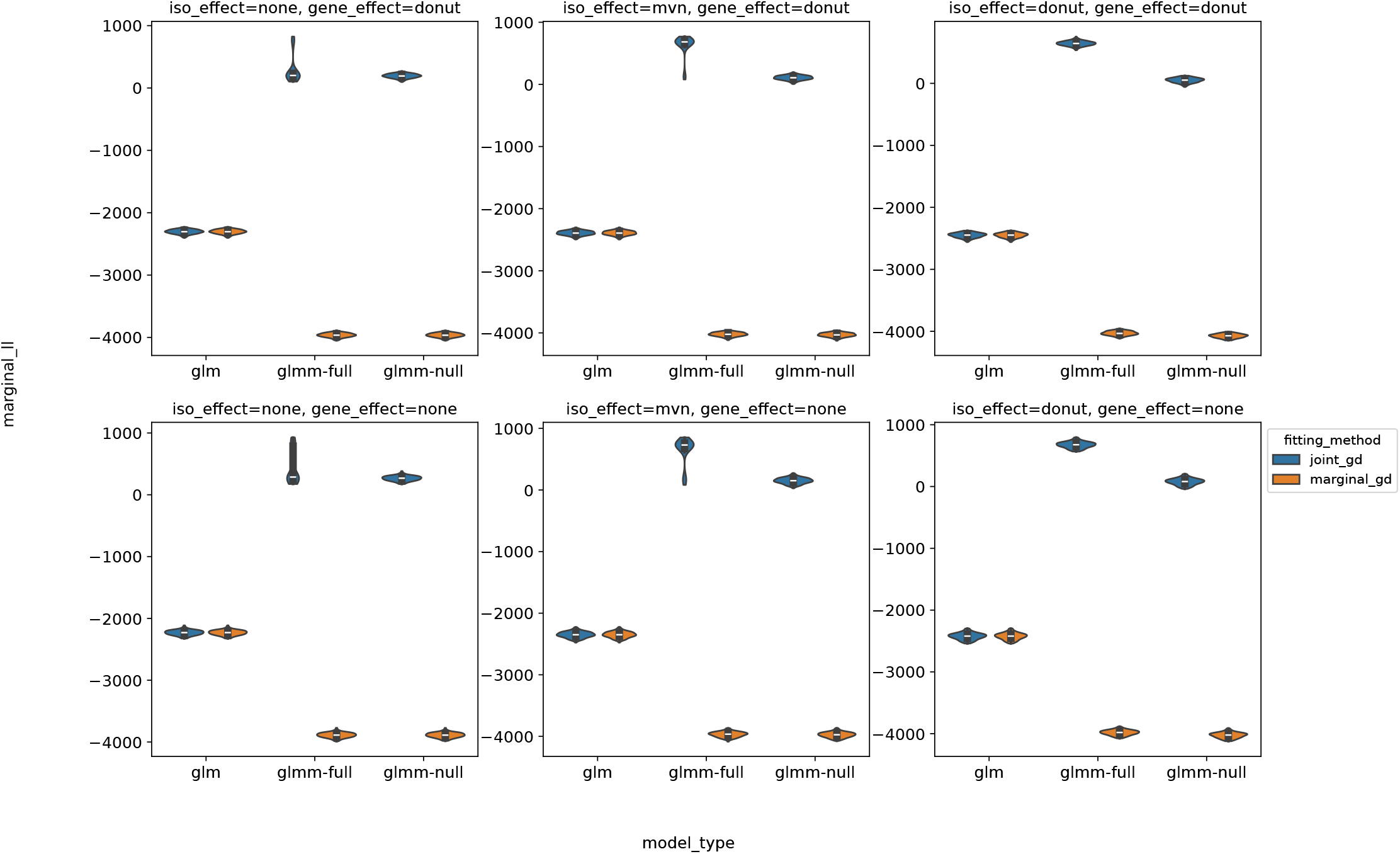
Marginal log-likelihood computed from Equation (12) (higher is better). The ‘joint-gd’ approach can achieves artificially high likelihood by minimizing variance *σ*, leading to exploding point density *h*(*β, σ, θ*; *ν* ≃ 0). Even with ‘marginal-gd’, solutions suffer from local optima, sometimes even producing negative likelihood ratios. Similar optimization challenges have been reported in spatial variability tests for gene expression[BGJ^+^21] where the integral over random effects is approximated using variational inference.

**Figure 8.**
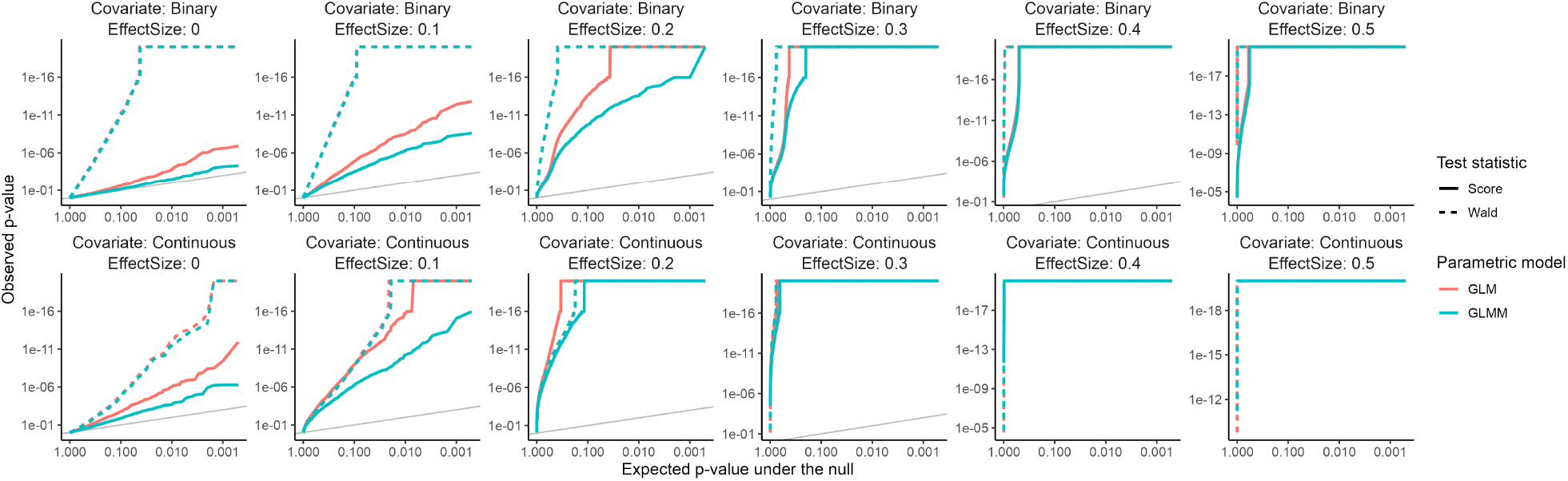
GLMM effectively reduces p-value inflation arising from spatial confounding (the first column) when coupled with the score test and fitted under the null hypothesis H_0_ : *β* = 0. In contrast, the Wald test requires estimation under the alternative (ℋ_1_ : *β* ≠ 0), thus suffering from inflated 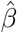 estimates from both GLM and GLMM models.

where *ℋ* (*β*) is the Hessian matrix (negative Fisher information) and *L*(*β*) the log-likelihood function.

As shown in Figure 8, the score test with GLMM successfully controls for spatial confounding effects, while the Wald test suffers from inflated estimates regardless of model choice. We observed consistent results with different optimization approaches (e.g. (15) or (16) for GLMM). In particular, the approximation-free GLM estimates of *β* from IWLS are similarly inaccurate, and the resulting GLM-Wald test produces the worst performance.

## Footnotes

1 For 10X Visium data (*n ≤* 4992), computing and storing a dense *K*_*sp*_ requires approximately 5000 *×* 5000 *×* 4 bytes (float32) *≈* 100 MB RAM.

2 This approach resembles block-based HSIC[ZFGS18] where *K* is partitioned into blocks and *K*_*ij*_ ≡ 0 for entries across different blocks, which also suffers from power loss in empirical experiments.

3 Some nodes may not connected to any other nodes in a k-mutual-neighbors graph. We set *W*_*ii*_ = 1 for these isolated nodes to avoid all-zero rows and columns in the Laplacian. After inverse, isolated nodes will have zero cross-covariance with other nodes in *K*

4 Approximately 50% of spots in 10X Visium data with Illumina sequencing (80% spots with Oxford Nanopore) have zero captured UMI of a median gene.

## Reference

1. Chen, M. & Manley, J. L. Mechanisms of alternative splicing regulation: insights from molecular and genomics approaches. Nat Rev Mol Cell Biol 10, 741–54 (2009).

2. Tian, B. & Manley, J. L. Alternative polyadenylation of mRNA precursors. Nat Rev Mol Cell Biol 18, 18–30 (2017).

3. Wang, E. T. et al. Alternative isoform regulation in human tissue transcriptomes. Nature 456, 470–6 (2008).

4. Weyn-Vanhentenryck, S. M. et al. Precise temporal regulation of alternative splicing during neural development. Nat Commun 9, 2189 (2018).

5. Zhang, X. et al. Cell-Type-Specific Alternative Splicing Governs Cell Fate in the Developing Cerebral Cortex. Cell 166, 1147–1162 e15 (2016).

6. Ule, J. et al. Nova regulates brain-specific splicing to shape the synapse. Nat Genet 37, 844–52 (2005).

7. Beffert, U. et al. Modulation of synaptic plasticity and memory by Reelin involves differential splicing of the lipoprotein receptor Apoer2. Neuron 47, 567–79 (2005).

8. Li, Y. I. et al. RNA splicing is a primary link between genetic variation and disease. Science 352, 600–4 (2016).

9. Walker, R. L. et al. Genetic Control of Expression and Splicing in Developing Human Brain Informs Disease Mechanisms. Cell 179, 750-771.e22 (2019).

10. Raj, T. et al. Integrative transcriptome analyses of the aging brain implicate altered splicing in Alzheimer’s disease susceptibility. Nat Genet 50, 1584–1592 (2018).

11. Liu, Z. et al. Mutations in the RNA Splicing Factor SF3B1 Promote Tumorigenesis through MYC Stabilization. Cancer Discov 10, 806–821 (2020).

12. Zhang, Y., Qian, J., Gu, C. & Yang, Y. Alternative splicing and cancer: a systematic review. Signal Transduct Target Ther 6, 78 (2021).

13. Jbara, A. et al. RBFOX2 modulates a metastatic signature of alternative splicing in pancreatic cancer. Nature 617, 147–153 (2023).

14. Stahl, P. L. et al. Visualization and analysis of gene expression in tissue sections by spatial transcriptomics. Science 353, 78–82 (2016).

15. Liu, Y. et al. High-Spatial-Resolution Multi-Omics Sequencing via Deterministic Barcoding in Tissue. Cell 183, 1665-1681.e18 (2020).

16. Stickels, R. R. et al. Highly sensitive spatial transcriptomics at near-cellular resolution with Slide-seqV2. Nat Biotechnol 39, 313–319 (2021).

17. Joglekar, A. et al. A spatially resolved brain region- and cell type-specific isoform atlas of the postnatal mouse brain. Nat Commun 12, 463 (2021).

18. Lebrigand, K. et al. The spatial landscape of gene expression isoforms in tissue sections. Nucleic Acids Research 51, e47 (2023).

19. Chen, K. H., Boettiger, A. N., Moffitt, J. R., Wang, S. & Zhuang, X. Spatially resolved, highly multiplexed RNA profiling in single cells. Science 348, aaa6090 (2015).

20. Cohen, L. et al. Whole-transcriptome-scale and isoform-resolved spatial imaging of single cells in complex tissues. 2025.08.27.672533 Preprint at 10.1101/2025.08.27.672533 (2025).

21. Ji, G. et al. stAPAminer: Mining Spatial Patterns of Alternative Polyadenylation for Spatially Resolved Transcriptomic Studies. Genomics Proteomics Bioinformatics 21, 601–618 (2023).

22. Olivieri, J. & Salzman, J. Analysis of RNA processing directly from spatial transcriptomics data reveals previously unknown regulation. bioRxiv https://doi.org/10.1101/2023.03.13.532412 (2023) xdoi:10.1101/2023.03.13.532412.

23. Fu, Y. et al. Single cell and spatial alternative splicing analysis with Nanopore long read sequencing. Nat Commun 16, 6654 (2025).

24. Buen Abad Najar, C. F., Yosef, N. & Lareau, L. F. Coverage-dependent bias creates the appearance of binary splicing in single cells. eLife 9, e54603 (2020).

25. Isaev, K. & Knowles, D. A. Investigating RNA splicing as a source of cellular diversity using a binomial mixture model. in 163–175 (PMLR, 2024).

26. Rich, J. M. et al. The impact of package selection and versioning on single-cell RNA-seq analysis. bioRxiv (2024).

27. Edsgard, D., Johnsson, P. & Sandberg, R. Identification of spatial expression trends in single-cell gene expression data. Nat Methods 15, 339–342 (2018).

28. Svensson, V., Teichmann, S. A. & Stegle, O. SpatialDE: identification of spatially variable genes. Nat Methods 15, 343–346 (2018).

29. Sun, S., Zhu, J. & Zhou, X. Statistical analysis of spatial expression patterns for spatially resolved transcriptomic studies. Nat Methods 17, 193–200 (2020).

30. Zhu, J., Sun, S. & Zhou, X. SPARK-X: non-parametric modeling enables scalable and robust detection of spatial expression patterns for large spatial transcriptomic studies. Genome Biology 22, 184 (2021).

31. Dries, R. et al. Giotto: a toolbox for integrative analysis and visualization of spatial expression data. Genome Biol 22, 78 (2021).

32. Chen, C., Kim, H. J. & Yang, P. Evaluating spatially variable gene detection methods for spatial transcriptomics data. Genome Biol 25, 18 (2024).

33. Gretton, A. et al. A kernel statistical test of independence. Advances in neural information processing systems 20, (2007).

34. Zhang, K., Peters, J., Janzing, D. & Schölkopf, B. Kernel-based conditional independence test and application in causal discovery. arXiv preprint arXiv:1202.3775 (2012).

35. Su, J. et al. Smoother: a unified and modular framework for incorporating structural dependency in spatial omics data. Genome Biol 24, 291 (2023).

36. Booeshaghi, A. S. et al. Isoform cell-type specificity in the mouse primary motor cortex. Nature 598, 195–199 (2021).

37. Joglekar, A. et al. Single-cell long-read sequencing-based mapping reveals specialized splicing patterns in developing and adult mouse and human brain. Nat Neurosci 27, 1051–1063 (2024).

38. Patowary, A. et al. Developmental isoform diversity in the human neocortex informs neuropsychiatric risk mechanisms. Science 384, eadh7688 (2024).

39. Olivieri, J. E. et al. RNA splicing programs define tissue compartments and cell types at single-cell resolution. Elife 10, (2021).

40. Zhao, W. et al. POSTAR3: an updated platform for exploring post-transcriptional regulation coordinated by RNA-binding proteins. Nucleic Acids Res 50, D287–D294 (2022).

41. Wheeler, E. C. et al. Integrative RNA-omics Discovers GNAS Alternative Splicing as a Phenotypic Driver of Splicing Factor–Mutant Neoplasms. Cancer Discovery 12, 836–855 (2022).

42. Miura, P., Shenker, S., Andreu-Agullo, C., Westholm, J. O. & Lai, E. C. Widespread and extensive lengthening of 3′ UTRs in the mammalian brain. Genome Res. 23, 812–825 (2013).

43. Ogorodnikov, A. et al. Transcriptome 3′end organization by PCF11 links alternative polyadenylation to formation and neuronal differentiation of neuroblastoma. Nat Commun 9, 5331 (2018).

44. 10x Genomics. Adult Mouse Brain Coronal Section (Fresh Frozen) obtained from BioIVT Asterand, Spatial Gene Expression dataset analyzed using Space Ranger 2.1.0. (2023).

45. Alfonso-Gonzalez, C. et al. Sites of transcription initiation drive mRNA isoform selection. Cell 186, 2438-2455.e22 (2023).

46. 10x Genomics. Mouse Brain Coronal Section (Fresh Frozen) from healthy 9-week-old C57BL/6 mouse, In Situ Gene Expression dataset analyzed using Xenium Onboard Analysis 3.0.0. (2024).

47. Hwang, H. W. et al. cTag-PAPERCLIP Reveals Alternative Polyadenylation Promotes Cell-Type Specific Protein Diversity and Shifts Araf Isoforms with Microglia Activation. Neuron 95, 1334–1349 e5 (2017).

48. Fisher, E. & Feng, J. RNA splicing regulators play critical roles in neurogenesis. Wiley Interdiscip Rev RNA 13, e1728 (2022).

49. Jacko, M. et al. Rbfox Splicing Factors Promote Neuronal Maturation and Axon Initial Segment Assembly. Neuron 97, 853–868 e6 (2018).

50. Rehfeld, F. et al. The RNA-binding protein ARPP21 controls dendritic branching by functionally opposing the miRNA it hosts. Nat Commun 9, 1235 (2018).

51. Chatrikhi, R. et al. RNA Binding Protein CELF2 Regulates Signal-Induced Alternative Polyadenylation by Competing with Enhancers of the Polyadenylation Machinery. Cell Rep 28, 2795–2806 e3 (2019).

52. Van Nostrand, E. L. et al. A large-scale binding and functional map of human RNA-binding proteins. Nature 583, 711–719 (2020).

53. Gazzara, M. R. et al. Ancient antagonism between CELF and RBFOX families tunes mRNA splicing outcomes. Genome Res. 27, 1360–1370 (2017).

54. Berto, S., Usui, N., Konopka, G. & Fogel, B. L. ELAVL2-regulated transcriptional and splicing networks in human neurons link neurodevelopment and autism. Human Molecular Genetics 25, 2451–2464 (2016).

55. McGurk, M. P., McWatters, D. C. & Burge, C. B. KATMAP: Inferring splicing factor activity and regulatory targets from knockdown data. bioRxiv 2024.06. 25.600605 (2024).

56. Maynard, K. R. et al. Transcriptome-scale spatial gene expression in the human dorsolateral prefrontal cortex. Nat Neurosci 24, 425–436 (2021).

57. Ito, H. et al. Sept8 controls the binding of vesicle-associated membrane protein 2 to synaptophysin. Journal of Neurochemistry 108, 867–880 (2009).

58. Yuan, Y. et al. Cell type-specific CLIP reveals that NOVA regulates cytoskeleton interactions in motoneurons. Genome Biology 19, 117 (2018).

59. Greenwald, A. C. et al. Integrative spatial analysis reveals a multi-layered organization of glioblastoma. Cell 187, 2485–2501 e26 (2024).

60. Ren, Y. et al. Spatial transcriptomics reveals niche-specific enrichment and vulnerabilities of radial glial stem-like cells in malignant gliomas. Nat Commun 14, 1028 (2023).

61. Vandiedonck, C. et al. Pervasive haplotypic variation in the spliceo-transcriptome of the human major histocompatibility complex. Genome Res. 21, 1042–1054 (2011).

62. Puttick, C. et al. MHC Hammer reveals genetic and non-genetic HLA disruption in cancer evolution. Nat Genet 56, 2121–2131 (2024).

63. Kahles, A. et al. Comprehensive Analysis of Alternative Splicing Across Tumors from 8,705 Patients. Cancer Cell 34, 211-224.e6 (2018).

64. Song, X. et al. A Single-Cell Atlas of RNA Alternative Splicing in the Glioma-Immune Ecosystem. bioRxiv 2025.03. 26.645511 (2025).

65. Stassen, O. et al. GFAPdelta/GFAPalpha ratio directs astrocytoma gene expression towards a more malignant profile. Oncotarget 8, 88104–88121 (2017).

66. Uceda-Castro, R. et al. GFAP splice variants fine-tune glioma cell invasion and tumour dynamics by modulating migration persistence. Sci Rep 12, 424 (2022).

67. Larionova, T. D. et al. Alternative RNA splicing modulates ribosomal composition and determines the spatial phenotype of glioblastoma cells. Nat Cell Biol 24, 1541–1557 (2022).

68. Fuhrmann, D. C., Mondorf, A., Beifuß, J., Jung, M. & Brüne, B. Hypoxia inhibits ferritinophagy, increases mitochondrial ferritin, and protects from ferroptosis. Redox Biology 36, 101670 (2020).

69. Iijima, T., Hidaka, C. & Iijima, Y. Spatio-temporal regulations and functions of neuronal alternative RNA splicing in developing and adult brains. Neurosci Res 109, 1–8 (2016).

70. Chang, Y. et al. Graph Fourier transform for spatial omics representation and analyses of complex organs. Nat Commun 15, 7467 (2024).

71. Pedregosa, F. et al. Scikit-learn: Machine Learning in Python. Journal of Machine Learning Research 12, 2825–2830 (2011).

72. Liu, H., Tang, Y. & Zhang, H. H. A new chi-square approximation to the distribution of non-negative definite quadratic forms in non-central normal variables. Computational Statistics & Data Analysis 53, 853–856 (2009).

73. Benjamini, Y. & Hochberg, Y. Controlling the False Discovery Rate - a Practical and Powerful Approach to Multiple Testing. J R Stat Soc B 57, 289–300 (1995).

74. Liao, J. Y. et al. EuRBPDB: a comprehensive resource for annotation, functional and oncological investigation of eukaryotic RNA binding proteins (RBPs). Nucleic Acids Res 48, D307–D313 (2020).

75. Ray, D. et al. A compendium of RNA-binding motifs for decoding gene regulation. Nature 499, 172–7 (2013).

76. Tremblay, B. J.-M. universalmotif: An R package for biological motif analysis. Journal of Open Source Software 9, 7012 (2024).

77. Bailey, T. L., Johnson, J., Grant, C. E. & Noble, W. S. The MEME Suite. Nucleic Acids Res 43, W39–49 (2015).

78. Luo, Y. et al. New developments on the Encyclopedia of DNA Elements (ENCODE) data portal. Nucleic Acids Res 48, D882–D889 (2020).

79. Kowalski, M. H. et al. Multiplexed single-cell characterization of alternative polyadenylation regulators. Cell 187, 4408–4425 e23 (2024).

80. Trincado, J. L. et al. SUPPA2: fast, accurate, and uncertainty-aware differential splicing analysis across multiple conditions. Genome Biol 19, 40 (2018).

81. Gohr, A. & Irimia, M. Matt: Unix tools for alternative splicing analysis. Bioinformatics 35, 130–132 (2019).

82. Patrick, R. et al. Sierra: discovery of differential transcript usage from polyA-captured single-cell RNA-seq data. Genome Biology 21, 167 (2020).

83. Tasic, B. et al. Shared and distinct transcriptomic cell types across neocortical areas. Nature 563, 72–78 (2018).

84. Moakley, D. F. et al. Reverse engineering neuron-type-specific and type-orthogonal splicing-regulatory networks using diverse cellular transcriptomes. Cell Reports 44, 115898 (2025).

85. Song, J. et al. DEMINERS enables clinical metagenomics and comparative transcriptomic analysis by increasing throughput and accuracy of nanopore direct RNA sequencing. Genome Biol 26, 76 (2025).

## References

[BANYL20] Carlos F Buen Abad Najar, Nir Yosef, and Liana F Lareau. Coverage-dependent bias creates the appearance of binary splicing in single cells. Elife, 9:e54603, 2020.

[BGJ+21] Nuha BinTayyash, Sokratia Georgaka, S. John, Sumon Ahmed, Alexis Boukouvalas, James Hensman, and Magnus Rattray. Non-parametric modelling of temporal and spatial counts data from rna-seq experiments. Bioinformatics, 37(21): 3788–3795, 2021.

[BZG+24] Zhiliang Bai, Dingyao Zhang, Yan Gao, Bo Tao, Daiwei Zhang, Shuozhen Bao, Archibald Enninful, Yadong Wang, Haikuo Li, Graham Su, et al. Spatially exploring rna biology in archival formalin-fixed paraffin-embedded tissues. Cell, 187(23): 6760–6779, 2024.

[Chu97] Fan R. K. Chung. Spectral Graph Theory. American Mathematical Society, Providence, RI, 1997.

[CLJ+24] Yuzhou Chang, Jixin Liu, Yi Jiang, Anjun Ma, Yao Yu Yeo, Qi Guo, Megan McNutt, Jordan E Krull, Scott J Rodig, Dan H Barouch, et al. Graph fourier transform for spatial omics representation and analyses of complex organs. Nature Communications, 15(1): 7467, 2024.

[FKR+25] Yuntian Fu, Heonseok Kim, Sharmili Roy, Sijia Huang, Jenea I Adams, Susan M Grimes, Billy T Lau, Anuja Sathe, Hanlee P Ji, and Nancy R Zhang. Single cell and spatial alternative splicing analysis with nanopore long read sequencing. Nature Communications, 16(1): 6654, 2025.

[GBSS05] Arthur Gretton, Olivier Bousquet, Alex Smola, and Bernhard Schölkopf. Measuring statistical dependence with hilbert-schmidt norms. In Proceedings of the 16th international conference on Algorithmic Learning Theory, pages 63–77, 2005.

[JL21] Shaobo Jin and Youngjo Lee. A review of h-likelihood and hierarchical generalized linear model. Wiley Interdisciplinary Reviews: Computational Statistics, 13(5):e1527, 2021.

[JPM+21] Anoushka Joglekar, Andrey Prjibelski, Ahmed Mahfouz, Paul Collier, Susan Lin, Anna Katharina Schlusche, Jordan Marrocco, Stephen R Williams, Bettina Haase, Ashley Hayes, et al. A spatially resolved brain region-and cell type-specific isoform atlas of the postnatal mouse brain. Nature communications, 12(1): 463, 2021.

[JTZ+23] Guoli Ji, Qi Tang, Sheng Zhu, Junyi Zhu, Pengchao Ye, Shuting Xia, and Xiaohui Wu. stapaminer: mining spatial patterns of alternative polyadenylation for spatially resolved transcriptomic studies. Genomics, Proteomics & Bioinformatics, 21(3): 601–618, 2023.

[LBT+23] Kevin Lebrigand, Joseph Bergenstråhle, Kim Thrane, Annelie Mollbrink, Konstantinos Meletis, Pascal Barbry, Rainer Waldmann, and Joakim Lundeberg. The spatial landscape of gene expression isoforms in tissue sections. Nucleic Acids Research, 51(8):e47–e47, 2023.

[LN96] Youngjo Lee and John A Nelder. Hierarchical generalized linear models. Journal of the Royal Statistical Society: Series B (Methodological), 58(4): 619–656, 1996.

[LTZ09] Huan Liu, Yongqiang Tang, and Hao Helen Zhang. A new chi-square approximation to the distribution of non-negative definite quadratic forms in non-central normal variables. Computational Statistics & Data Analysis, 53(4): 853–856, 2009.

[OS23] Julia Olivieri and Julia Salzman. Analysis of rna processing directly from spatial transcriptomics data reveals previously unknown regulation. BioRxiv, 2023.

[PYPA22] Junyoung Park, Changwon Yoon, Cheolwoo Park, and Jeongyoun Ahn. Kernel methods for radial transformed compositional data with many zeros. In International Conference on Machine Learning, pages 17458–17472. PMLR, 2022.

[RE76] Paul Robert and Yves Escoufier. A unifying tool for linear multivariate statistical methods: the rv-coefficient. Journal of the Royal Statistical Society Series C: Applied Statistics, 25(3): 257–265, 1976.

[SRF+23] Jiayu Su, Jean-Baptiste Reynier, Xi Fu, Guojie Zhong, Jiahao Jiang, Rydberg Supo Escalante, Yiping Wang, Luis Aparicio, Benjamin Izar, David A Knowles, et al. Smoother: a unified and modular framework for incorporating structural dependency in spatial omics data. Genome Biology, 24(1): 291, 2023.

[SS01] Bernhard Scholkopf and Alexander J. Smola. Learning with Kernels: Support Vector Machines, Regularization, Optimization, and Beyond. MIT Press, Cambridge, MA, USA, 2001.

[STS18] Valentine Svensson, Sarah A Teichmann, and Oliver Stegle. Spatialde: identification of spatially variable genes. Nature methods, 15(5): 343–346, 2018.

[SZZ20] Shiquan Sun, Jiaqiang Zhu, and Xiang Zhou. Statistical analysis of spatial expression patterns for spatially resolved transcriptomic studies. Nature methods, 17(2): 193–200, 2020.

[ZFGS18] Qinyi Zhang, Sarah Filippi, Arthur Gretton, and Dino Sejdinovic. Large-scale kernel methods for independence testing. Statistics and Computing, 28: 113–130, 2018.

[ZPJS12] Kun Zhang, Jonas Peters, Dominik Janzing, and Bernhard Schölkopf. Kernel-based conditional independence test and application in causal discovery. arXiv preprint arXiv:1202.3775, 2012.

[ZSZ21] Jiaqiang Zhu, Shiquan Sun, and Xiang Zhou. Spark-x: non-parametric modeling enables scalable and robust detection of spatial expression patterns for large spatial transcriptomic studies. Genome biology, 22(1): 184, 2021.

